# Transforming inert cycloalkanes into α,ω-diamines through designed enzymatic cascade catalysis

**DOI:** 10.1101/2022.11.15.516597

**Authors:** Zhongwei Zhang, Lin Fang, Fei Wang, Yu Deng, Zhengbin Jiang, Aitao Li

## Abstract

Aliphatic α,ω-diamines (DAs) are important monomer precursors in polyamide plastic manufacturing. However, the dominant industrial process for DA synthesis involves energy-intensive, multistage chemical reactions that are harmful to the environment. For instance, 1,6-hexanediamine (HMD), one of most prominent monomers in nylon-66 synthesis, is mainly synthesized with currently high technological control by butadiene hydrocyanation, which suffers from the use of highly toxic hydrogen cyanide, unsatisfactory selectivity and a complex separation process. Thus, the development of sustainable green DA synthetic routes is highly desired. Herein, we report an efficient one-pot *in vivo* biocatalytic cascade for the transformation of cycloalkanes into DAs with the aid of advanced techniques, including the RetroBioCat tool for biocatalytic route design, enzyme mining for finding appropriate enzymes and microbial consortia construction for efficient pathway assembly. As a result, DAs are successfully produced by the developed microbial consortia-based biocatalytic system, especially HMD, and product concentrations as high as 16.5 mM and 7.6 mM are achieved when using cyclohexanol (CHOL) or cyclohexane (CH) as substrates, respectively. This also represents the highest HMD biosynthesis productivity to date. Other cycloalkanes also serve as substrates, indicating the generality of our approach.

## Introduction

Aliphatic linear α,ω-diamines (DAs) are large-volume commodity chemicals consumed principally as monomer precursors in polyamide plastic manufacturing that have a broad range of applications in the fabrication of engineering plastics, mechanical accessories, fibres, films and other applications^1,2^. Among them, 1,6-hexanediamine (HMD), which is produced annually on a *ca*. 1.2 Mt scale^3^, is the most prominent example in terms of production volume. In addition, HMD can be used as a nylon-66 monomer (made from HMD and adipic acid by polycondensation)^4,5^, with nylon-66 being one of the most important polyamides for the textile and plastic industries. The global nylon-66 market size was estimated to be $16.29 billion in 2019 and is expected to rise by 6.5% annually from 2019 to 2027^6^.

Today’s dominant industrial process for HMD production, which was developed by DuPont^7,8^, involves energy-intensive, multistage chemical reactions with butadiene as a starting material (**Fig. 1a**). Although successfully applied on a large scale, this process still suffers from the use of highly toxic hydrogen cyanide, severe reaction conditions and unsatisfactory selectivity. To overcome these problems, over many years and even decades, attempts have been made to find more environmentally benign processes by replacing the hazardous reagents or/and changing the raw material basis^9,10^. Some elegant approaches avoiding the use of cyanides and starting from readily available materials (e.g., 1,6-hexanediol) have been developed, but limitations such as high reaction temperature, elevated pressure and high catalyst loading still exist^11,12^. In contrast, biological methods offer the possibility of mild reaction conditions and excellent selectivity. Nevertheless, until now, only one biocatalytic route from adipic acid (AA) to HMD was reported *via* carboxylic acid reductases (CARs) and transaminases (TAs) involving the catalysis of two rounds of reduction/amination reactions, which gave only *ca*. 3 mM HMD with a 70% accumulation of 6-aminocaproic acid (6-ACA) intermediate^4^ as a by-product. Furthermore, although several patents have also been proposed for some nonnatural biological pathways for HMD biosynthesis^13^, none of them have been put into practice. Thus, a new biocatalytic route for the efficient synthesis of HMD remains a highly desirable goal.

**Figure 1.**
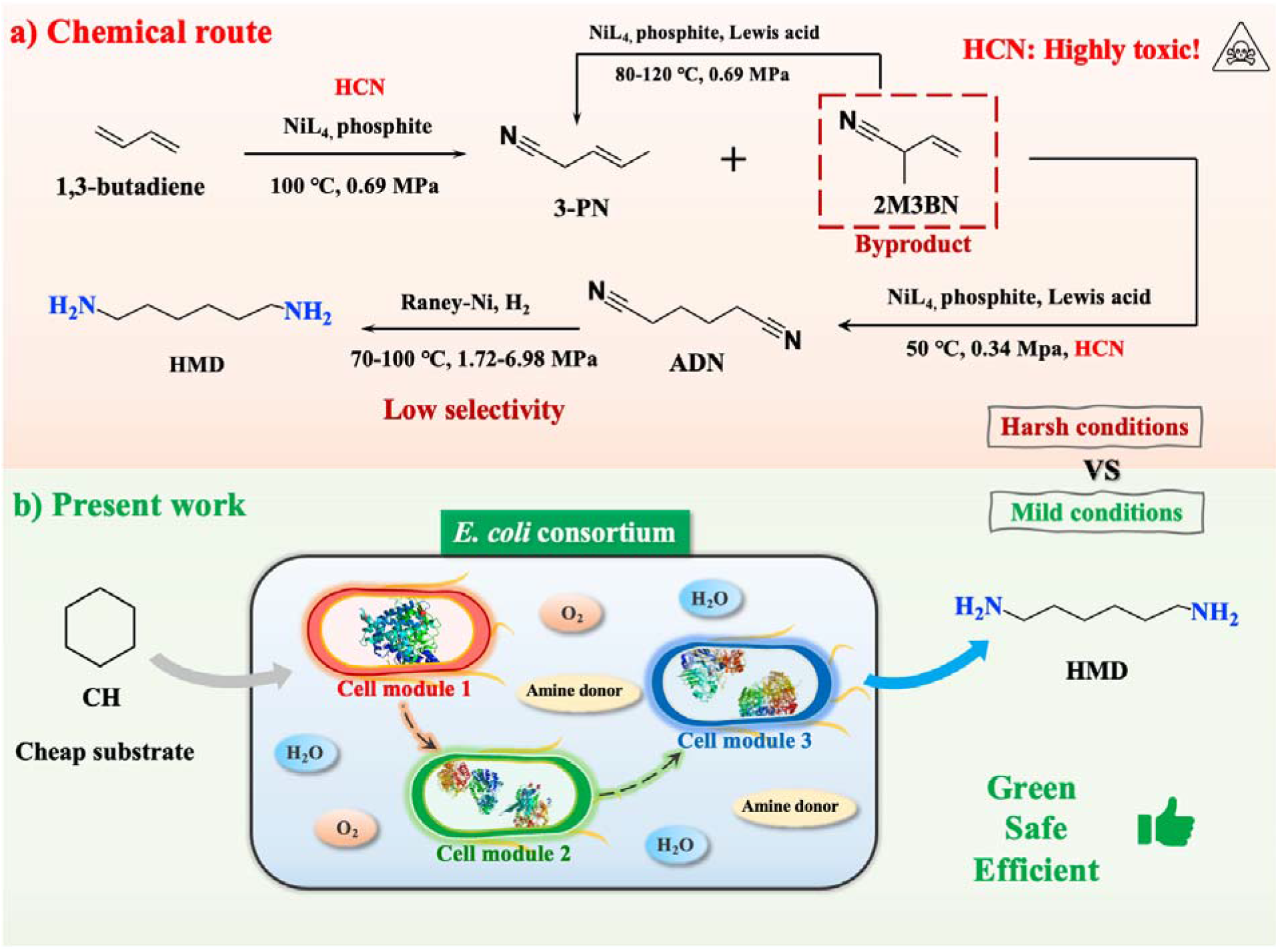
Industrial chemical and designed biocatalytic routes for HMD production. (a) Current industrial process for HMD synthesis involving multistage chemical reactions with butadiene as starting materials. **(b)** Designed one-pot biocatalytic cascade for HMD synthesis from cyclohexane (CH) using whole-cell catalysts in the form of *E. coli* consortium.

Currently, artificially designed cascade catalysis has attracted increasing attention^5,14–18^ because it enables one-pot efficient syntheses of targeted molecules from simple, inexpensive, and easily available starting materials. For instance, islatravir, which is an antiviral nucleoside analogue, was synthesized by Merck as an outstanding example of a five-enzyme *in vitro* cascade^19^. Terephthalic acid, which is used as a monomer precursor for polyethylene terephthalate fabrication, was produced through five enzyme-catalyzed *in vivo* consecutive oxidation reactions from *p*-xylene^17^. L-Homophenylalanine as a building block in chiral drug synthesis was recently produced with a designed enzymatic spontaneous chemical cascade, starting from simple benzaldehyde and pyruvate^20^. However, to achieve efficient targeted compound production in cascade catalysis, three issues concerning biocatalytic route design, enzyme selection and enzyme assembly should be addressed. With the rapid development of computational biology, enzymology and synthetic biology, an increasing number of techniques have been developed to address the three problems mentioned above, including (i) computational tools, such as retrobiosynthesis^21^, RetroPath2.0^22^ and RetroBioCat^23^, that have been established to guide retrosynthesis and facilitate the route design process; (ii) efficient enzyme screening methods, such as literature mining and genome mining, as well as advanced enzyme engineering methodologies, such as semirational design and machine learning, that have been employed to identify appropriate enzymes with desired properties^24–26^; and (iii) enzyme expression regulation strategies, such as promoter/RBS optimization, multiplasmid systems, gene copy modification, and microbial consortium-mediated pathway reconstitution, that have been used to solve the problems of unbalanced enzyme activity or expression ratios^27,28^ when assembling enzymes in a designed one-pot *in vivo/in vitro* biocatalytic cascade. Finally, the efficient production of target compounds can be achieved after optimizing the reaction conditions.

In this study, we designed an *in vivo* biocatalytic route for DA production based on retrosynthetic analysis. This biocatalytic cascade operates by means of a microbial consortia composed of three *E. coli* cell modules using simple and readily available cycloalkanes as substrates. With the conversion of cyclohexane (CH) to HMD as a model reaction **(Fig. 1b)**, to achieve a balanced enzyme activity or expression ratios in each *E. coli* cell module for facilitating HMD production, screening enzymes with desired properties and engineering cells with assigned functions were conducted and optimized. Finally, the efficient biosynthesis of HMD and related DAs from cycloalkanes or cyclohexanols was realized in a one-pot, ecologically and economically viable manner.

## Results

### Biocatalytic Cascade Design with Retrosynthesis

To achieve efficient DA biosynthesis, with the production of HMD as an example, biocatalytic retrosynthesis was carried out. The cascade route was designed using RetroBioCat (https://retrobiocat.com/) in either pathway explorer mode or network explorer mode. With pathway explorer mode, two routes starting from hexanediol (HDO) were discovered for HMD synthesis. As presented in **Fig. 2a**, in route 1, HDO can be subjected to two rounds of oxidation/amination with the corresponding alcohol dehydrogenase and transaminase to produce HMD in four enzymatic steps. In contrast, for route 2, although a green adiponitrile (ADN) production concept has been envisioned from HDO by the aldoxime dehydrogenase-catalyzed dehydration of dialdoximes^29^, the corresponding enzymes responsible for the nitrile reduction of AND to HMD are still unavailable. Thus, route 1 involving the conversion of HDO into HMD catalyzed by alcohol dehydrogenase and transaminase in a one-pot manner was considered. In addition, since efficient HDO biosynthesis from the inexpensive, readily available substrate cyclohexane (CH) or cyclohexanol (CHOL) was achieved in our previous study^14^, we envisioned a biocatalytic cascade for the production of HMD from CH or CHOL (**Fig. 2a**).

**Figure 2.**
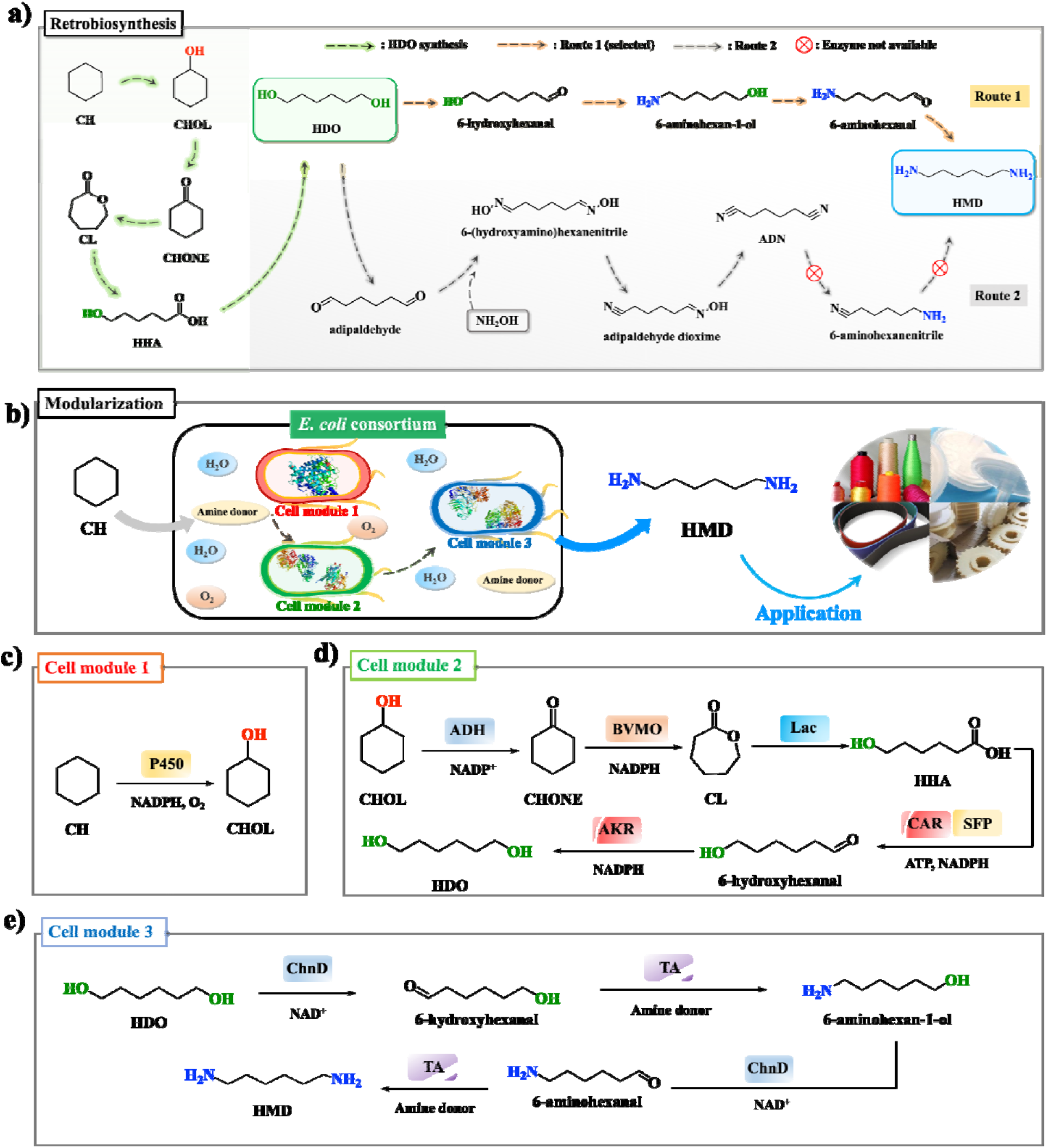
Design and modularization of a biocatalytic cascade for HMD synthesis. a) Retrobiosynthesis with a pathway explorer mode using RetroBioCat for HMD synthesis. b) Design of microbial consortium system consisting of three cell modules for converting cyclohexane (CH) to HMD in a one-pot manner. c) Cell module 1 involves P450-catalyzed hydroxylation of CH to cyclohexanol (CHOL). d) Cell module 2 includes ADH catalyzed oxidation of CHOL to cyclohexanone (CHONE), Baeyer-Villiger monooxygenase (BVMO) catalyzed oxidation of CHONE to ε-caprolactone (CL), lactonase (Lac) mediated hydrolysis of CL to HHA, followed by consecutive reduction with carboxylic acid reductase (CAR) (with the aid of phosphopantetheinyl transferase (SFP)) as well as endogenous aldo–keto reductase (AKR) to 1,6-hexanediol (HDO). e) Cell module 3 consists of two rounds of oxidation/animation reactions for converting HDO to HMD as a final product, catalyzed by an alcohol dehydrogenase (ChnD) and a transaminase (TA).

After designing this artificial biosynthetic route for HMD synthesis based on biocatalytic retrosynthesis, *in vivo* pathway reconstitution was adopted since costly steps (e.g., enzyme purification, addition of expensive cofactors) can thereby be avoided compared with *in vitro* approaches. In addition, to eliminate the potential expression burden and redox constraints caused by expressing multiple enzymes in a single cell, the concept of using a modularized microbial consortia was employed by distributing the enzymes among three cell modules (**Fig. 2b**). Specifically, cell module 1 includes a P450-catalyzed hydroxylation of inert CH to CHOL in the presence of the endogenous cofactor NADPH (**Fig. 2c**). Cell module 2 consists of an alcohol dehydrogenase (ADH)-catalyzed oxidation of CHOL to cyclohexanone (CHONE), a Baeyer-Villiger monooxygenase (BVMO)-mediated conversion of CHONE to ε-caprolactone (CL), a lactonase (Lac)-catalyzed hydrolysis of CL to 6-hydroxyhexanoic acid (HHA), and carboxylic acid reductase (CAR)- and endogenous aldo-keto reductase (AKR)-catalyzed consecutive reductions of HHA to HDO (**Fig. 2d**). Cell module 3 involves two rounds of oxidation/animation reactions for converting HDO to HMD, catalyzed by an alcohol dehydrogenase (ChnD) and a transaminase (TA) (**Fig. 2e**). Finally, cell engineering was conducted for optimal function, followed by cell combinations to form the microbial consortia that were used in HMD synthesis starting from the readily available substrate CHOL or CH. Therefore, by following a similar design, the biosynthesis of different DAs was achieved either from cycloalkanes or cycloalkanols.

### Engineering of *E. coli* Cell Module Catalysts

Next, the three basic cell modules were constructed and engineered in parallel, and *Escherichia coli* was used as a host cell system for expressing each module enzyme due to its fast growth, genetic tractability and availability of versatile genetic manipulation tools.

First, to engineer cell module 3, which was responsible for converting HDO to HMD, we used alcohol dehydrogenase (ChnD) from *Acinetobacter* sp. NCIMB9871^30^ and transaminases (TAs) originating from different sources. To find the best combination, a collection of *E. coli* whole-cell catalysts containing 59 transaminases were individually mixed with *E. coli* harbouring ChnD at a ratio of 1:1. The resulting cell mixtures were then employed to transform HDO into HMD in the presence of L/D-alanine (L/D-Ala) or isopropylamine as amine donors (**Supplementary Fig. 1**). The screening results showed that 5 out of 59 transaminases displayed excellent activity for HMD synthesis when coupled with ChnD (**Supplementary Figs 1 and 2**). Thus, the co-expression of ChnD with the five transaminases (CV, PP2159, SAV2614, PAK or SPO3471) in a single cell was then individually conducted (**Fig. 3b**). The plasmid pETDuet-1 carrying both TA and ChnD genes was transformed into *E. coli*, which resulted in five *E. coli* cells with different plasmid configurations (**Fig. 3a & 3b**). For comparison, the *E. coli* cells were then tested in the conversion of 20 mM HDO to HMD. As presented in **Fig. 3c**, except for *E. coli* (M3C), all cell catalysts clearly led to HMD production when using either L-Ala or isopropylamine as the amine donor. Among them, *E. coli* (M3E) showed the best catalytic performance and produced 17.0 mM HMD within 21 h. The results indicated that the engineered cell module 3 can realize biocatalytic HMD production from HDO, thus providing the possibility for subsequent HMD production from CH or CHOL.

**Figure 3.**
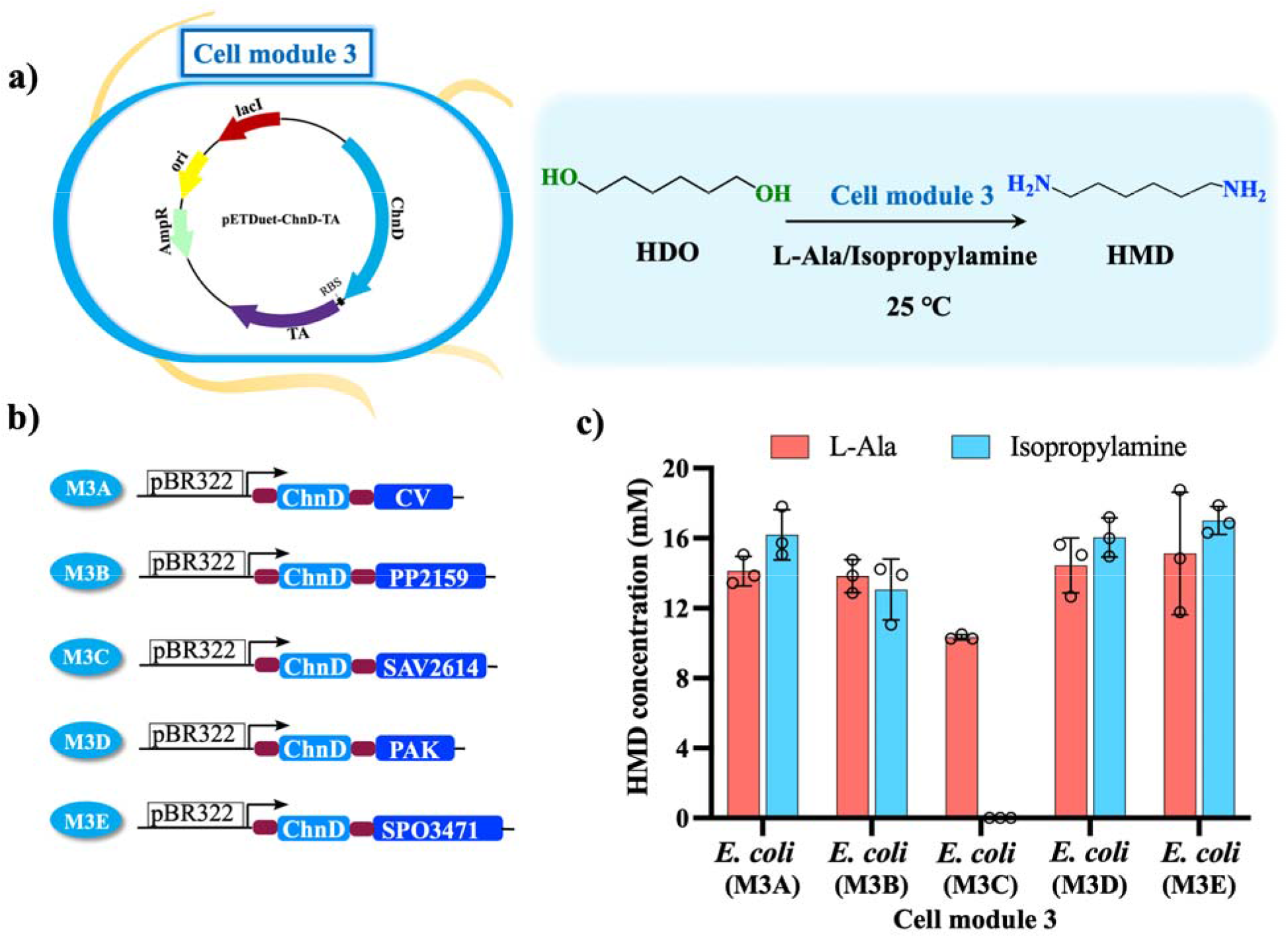
Biocatalytic HMD synthesis from HDO. a) Diagram of *E. coli*-catalyzed conversion of HDO to HMD. b) Construction of *E. coli* cells expressing alcohol dehydrogenase (ChnD) and transaminase (TA). pBR322: pETDuet-1; arrow: T7 promoter; red filed rectangle: RBS site. Light blue filled rectangle: ChnD gene; bule filled rectangle: transaminase gene (CV, PP2159, SAV2614, PAK and SPO3471). See **Supplementary Fig. 3** for SDS-PAGE of whole-cell proteins of cell module 3 expressed in *E. coli*. c) Engineered *E. coli* cell module 3 for biotransformation of HDO to HMD. Reaction conditions: *E. coli* cells expressing corresponding enzymes were resuspended in phosphate buffer (pH 8.0, 100 mM) at a cell density of 8 g CDW L^-1^, 20 mM HDO, reactions were performed at 25 °C, 220 rpm for 21 h, and the 100 mM L-Ala or isopropylamine used as amine donor.

In the construction of cell module 2 for converting CHOL to HDO, the co-expression of relevant enzymes in a single cell was attempted. In our previous study^14^, we achieved HDO synthesis from CHOL with a developed one-pot, one-step process consisting of two different cell module catalysts: the first cell transformed CHOL into HHA, and the second cell converted HHA to HDO (**Supplementary Fig. 4)**. To simplify this process, we speculated whether it could be possible to integrate all required enzymes into a single cell, hoping to realize direct HDO synthesis with comparable productivity. Theoretically, in this biocatalytic cascade, six enzymes need to be expressed in the same *E. coli* cell. However, based on the kinetic data of these enzymes^14^, it was found that lactonase (Lac)^31^ showed a much higher activity than the other enzymes. Thus, to balance its expression and activity, its integration into the genome for lower expression might be enough to achieve the required activity. In addition, since the endogenous aldo-keto reductase (AKR) responsible for aldehyde reduction in *E. coli* has been reported^32^, we suspected that endogenous AKR could also work for the reduction of 6-hydroxyhexanal to HDO, and the heterologous expression of AKR in *E. coli* could then be omitted (**Fig. 4b**). Therefore, plasmids carrying only four enzyme genes need to be constructed for heterologous expression in *E. coli*, namely Baeyer-Villiger monooxygenase (BVMO) from *Acinetobacter* sp. NCIMB9871^33^, alcohol dehydrogenase (ADH) from *Lactobacillus brevis*^34^, carboxylic acid reductase (CAR) from *Mycobacterium abscessus* ATCC 19977^35^, and phosphopantetheinyl transferase (SFP) from *Bacillus subtilis*^35^.

**Figure 4.**
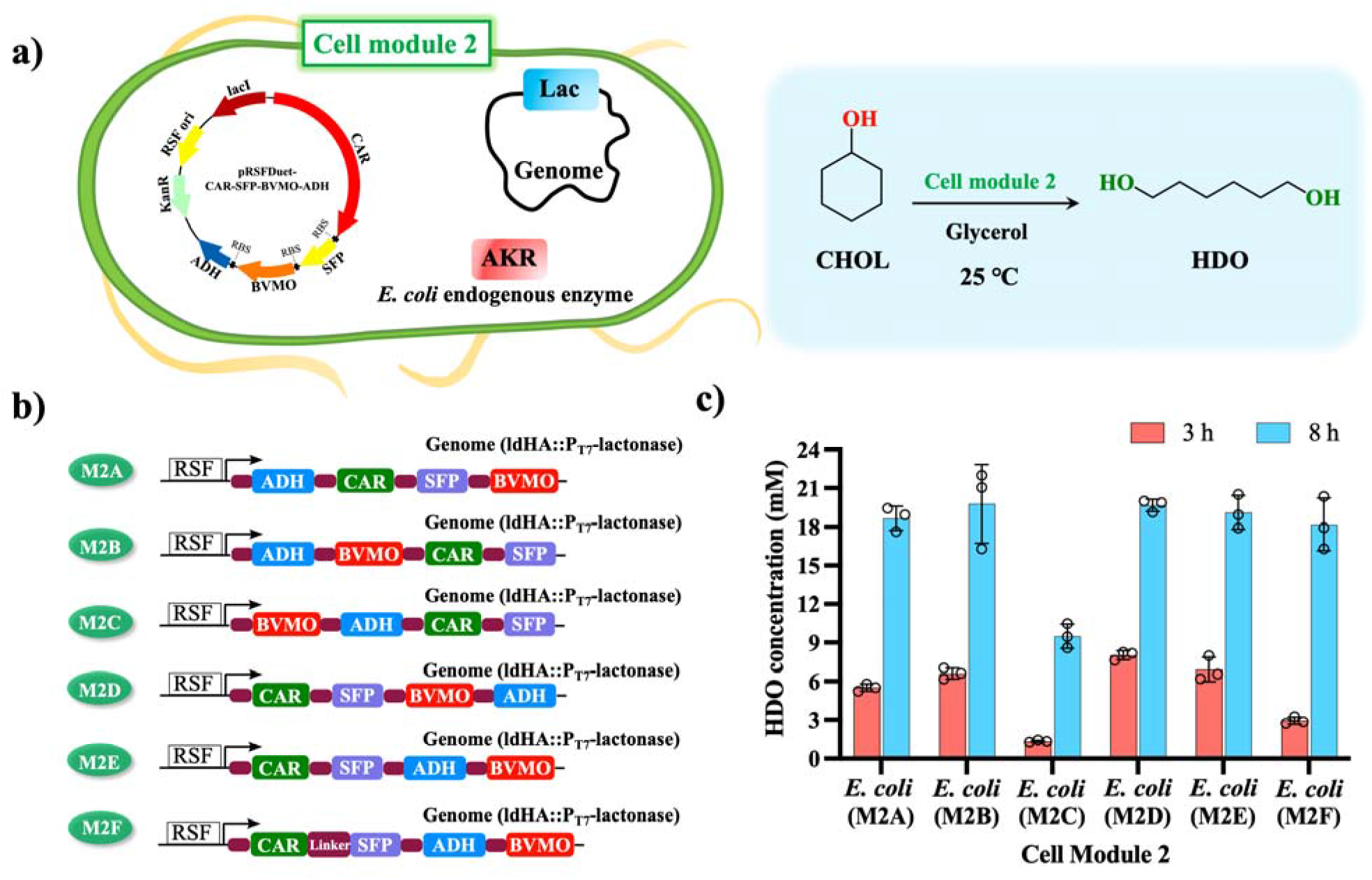
Biocatalytic HDO synthesis from CHOL. a) Diagram of constructed *E. coli*-catalyzed conversion of HDO from CHOL. b) Construction of *E. coli* cell expressing all necessary enzymes for converting CHOL to HDO. RSF: pRSFDuet-1; arrow: T7 promoter; dark red filed rectangle: RBS site. Light blue filled rectangle: ADH gene; green filled rectangle: CAR gene; purple filled rectangle: SFP gene; bright red filled rectangle: BVMO gene. The Lac gene was integrated to genome at ldHA site. See **Supplementary Fig. 6** for SDS-PAGE of whole-cell proteins of Module 2 expressed in *E. coli*. c) Engineered *E. coli* cell module 2 for the conversion of CHOL to HDO. Reaction conditions: *E. coli* cells expressing corresponding enzymes were resuspended in phosphate buffer (pH 8.0, 100 mM) at a cell density of 6 g CDW L^-1^, 20 mM CHOL, reactions were performed at 25 °C, 220 rpm for 8 h, cofactor NAD(P)H/ATP was provided by the *E. coli* host cells using 68 mM glycerol as an energy source.

Next, based on the different orders of enzyme genes in pRSFDuet-1 plasmids, the plasmids with different configurations (**Fig. 4b**) were transformed into *E. coli* cells with genomes that had been integrated with the Lac gene at the ldHA site, which resulted in five recombinant *E. coli* cells as cell module 2 catalysts. Furthermore, we also fused SFP with CAR to form a fused protein (**Fig. 4b, M2F**), hoping to further enhance the catalytic activity of CAR. As a result, a sixth recombinant *E. coli* cell was obtained. Then, for comparison, cells of the sixth type of *E. coli* were used as whole-cell catalysts for the conversion of CHOL to HDO. As shown in **Fig. 4c**, most of the engineered cell catalysts exhibited high catalytic performance. Among them, *E. coli* (M2B) and *E. coli* (M2D) yielded high product concentrations (6.6 mM and 8.0 mM) after 3 h of reaction, and 19.7 mM~19.8 mM HDO (corresponding to 98.5~99% yield) were obtained after 8 h of reaction. Although the remaining cell module catalysts, such as *E. coli* (M2A), *E. coli* (M2E) and *E. coli* (M2F), showed slightly lower catalytic activities, high product concentrations ranging from 18.2 to 19.1 mM were also obtained.

Compared with the two-cell biocatalytic system developed in our previous work^14^, the newly constructed single-cell catalyst system showed comparable catalytic performance for the conversion of CHOL to HDO, yet with less catalyst loading (**Supplementary Fig. 5**), which demonstrates the power of our cell engineering strategy. Thus, efficient CHOL conversion to HDO was successfully achieved with a single-cell catalyst, and *E. coli* (M2D) with the best catalytic performance was chosen for the following experiments. Moreover, since cell module 1, responsible for the hydroxylation of CH to CHOL, was constructed in our previous study, the following work of combining different cell modules (modules 1, 2 and 3) to form an *E. coli* consortium (EC) was conducted and tested using HMD synthesis from either CHOL or CH.

### One-pot HMD Biosynthesis from CHOL

With the three cell module catalysts in hand, we first tried a combination of cell modules 2 and 3 to form *E. coli* consortium 2_3 (EC2_3) for the biotransformation of CHOL to HMD. The biocatalytic cascade in a one-pot, two-step manner was first investigated. *E. coli* (M2D) with the best catalytic performance was used for converting CHOL to HDO. Once complete substrate conversion was achieved, each cell module 3 was individually added to initiate the reaction of HDO into HMD in the presence of L-Ala or isopropylamine (**Fig. 5a**). The results showed that EC2_3, composed of *E. coli* (M2D) and *E. coli* (M3D), gave the highest product concentration (11.6 mM HMD, 22 h) when using isopropylamine as an amine donor (**Fig. 5b**). Then, with this selected EC2_3, various reaction conditions, including glycerol concentration, cell density, and amine donor concentration, were optimized. It was found that the glycerol concentration influences the HMD productivity, and an optimal glycerol concentration of 68 mM was obtained when 20 mM CHOL was employed (**Supplementary Fig. 7a**). For the optimization of the catalyst loading and the ratio of modules 2 to 3 in the consortium, a ratio of 4:3 (modules 2:3) at a cell density of 14 g CDW L^-1^ showed the highest catalytic performance for HMD production (**Supplementary Fig. 7c**). In terms of the amount of amine donor used, with a fixed CHOL concentration of 20 mM, an optimal isopropylamine concentration of 80 mM was obtained (**Supplementary Fig. 7b)**. Therefore, under the optimized reaction conditions, the EC2_3-catalyzed conversion of CHOL to HMD was carried out. **Fig. 5c** shows the time course of this biocatalytic cascade reaction in a one-pot, two-step manner. CHOL was completely converted to HDO within 8 h. After that, cell module 3 was added to transform HDO into HMD, and the product HMD was gradually produced and reached a maximum of 12.6 mM at 32 h with almost no intermediate accumulation. It is worth mentioning that the lower product concentration (compared to the theoretical value) could be attributed to the loss of product caused by sampling as well as the dilution arising from the addition of the cell suspension of module 3.

**Figure 5.**
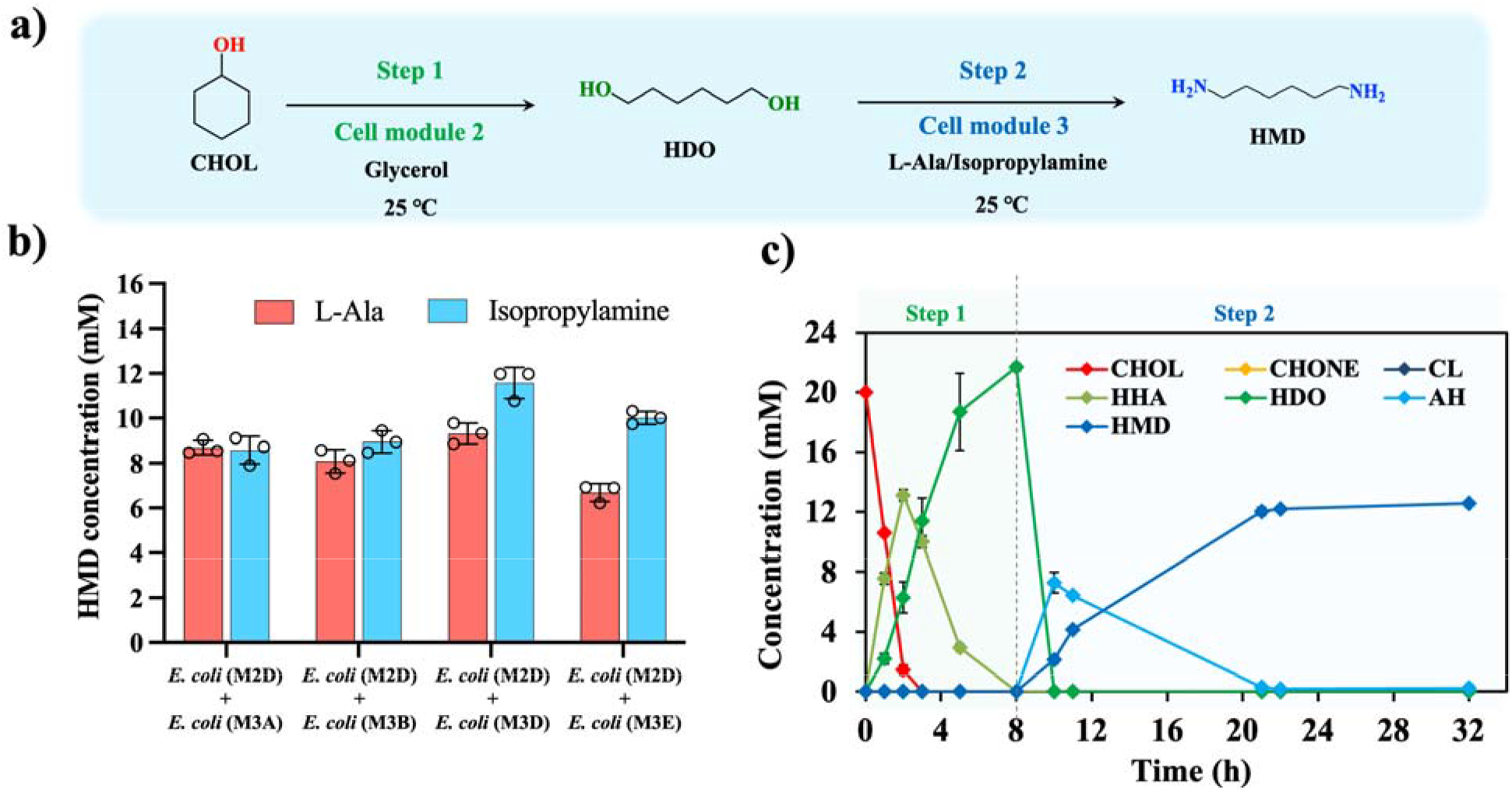
Construction of *E. coli* consortia for HMD production in a one-pot, two-step manner. a) Scheme of *E. coli* consortia EC2_3-catalyzed conversion of CHOL to HMD in a one-pot, two-step manner. b) Comparison of different EC2_3 in the biotransformation of CHOL to HMD in a one-pot, two-step manner. Reaction conditions: *E. coli* cell modules 2 and 3 at a ratio of 4:3 was resuspended in phosphate buffer (pH 8.0, 100 mM) with a total cell density of 14 g CDW L^-1^, 20 mM CHOL, reactions were performed at 25 °C, 220 rpm for 22 h, cofactor NAD(P)H/ATP was provided by the E. coli cells using 68 mM glycerol as an energy source and 60 mM L-Ala or isopropylamine was employed as amine donor. Cell module 3 and amine donor were added after 8 h reaction when the first step reaction was completed. c) Time course of EC2_3-catalyzed conversion of CHOL to HMD in one-pot, two-step manner. Reaction conditions: *E. coli* consortia 2_3 composed of *E. coli* (M2D) and *E. coli* (M3D) at a ratio of 4:3 was resuspended in phosphate buffer (pH 8.0, 100 mM) with a total cell density of 14 g CDW L^-1^, 20 mM CHOL, reactions were performed at 25 °C, 220 rpm for 32 h, cofactor NAD(P)H/ATP was provided by the *E. coli* cells using 68 mM glycerol as an energy source and 80 mM isopropylamine was added as amine donor. Cell module 3 and amine donor were added after 8 h reaction when the first step reaction was completed.

Next, encouraged by the positive results of HMD synthesis from CHOL in a one-pot, two-step manner, a more directed and efficient one-pot, one-step approach was attempted (**Fig. 6a**). During this process, both cell modules 2 and 3 were simultaneously added at the beginning of the reaction. First, we rescreened the different combinations of *E. coli* (M2D) with different cell module 3 types and found that the *E. coli* consortium consisting of *E. coli* (M2D) and *E. coli* (M3A) displayed the best productivity, producing 6~6.3 mM HMD within 22 h with isopropylamine or L-Ala as the amine donor (**Fig. 6b**). Therefore, this *E. coli* consortium was chosen for the subsequent experiments. However, one concern raised from this one-pot, one-step biocatalytic system is that the intermediate CHONE, which is produced from the module 2-catalyzed oxidation of CHOL, could be used as a substrate of transaminase in module 3 to produce cyclohexylamine as a byproduct. To verify this, the whole reaction process of this EC2_3-catalyzed conversion of CHOL to HMD was monitored. Interestingly, no cyclohexylamine was detected during the reaction, although cell module 3 expressing the corresponding transaminases clearly showed catalytic activity towards CHONE (**Supplementary Figs. 8 & 9**). We speculate that a reason for this could be the fortunate allocation of different enzymes among different host cells, which prevented cross-contamination by confining the intermediates inside the cells, meaning that the intermediate CHONE is quickly transformed into CL before being secreted out of the cell. Subsequently, various reaction conditions were likewise conducted in a one-pot, one-step manner (**Supplementary Fig. 10**). Of these, at a fixed CHOL substrate concentration of 20 mM, the optimal glycerol and isopropylamine concentrations proved to be 102 mM and 80 mM, respectively. The optimal ratio of modules 2 and 3 was 2:1, and higher cell loadings led to a higher HMD concentration. Finally, the one-pot, one-step HMD production from CHOL was carried out under the optimized conditions with the pH maintained at 8.0. As shown in **Fig. 6c**, complete CHOL consumption was quickly achieved within 4 h, and the main intermediates HDO, HHA and 6-amino-1-hexanol (AH) were first formed and then converted. The highest HMD concentration reached 16.5 mM with very little intermediate accumulation, which corresponds to a titre of 1.91 g/L (**Fig. 6c**).

**Figure 6.**
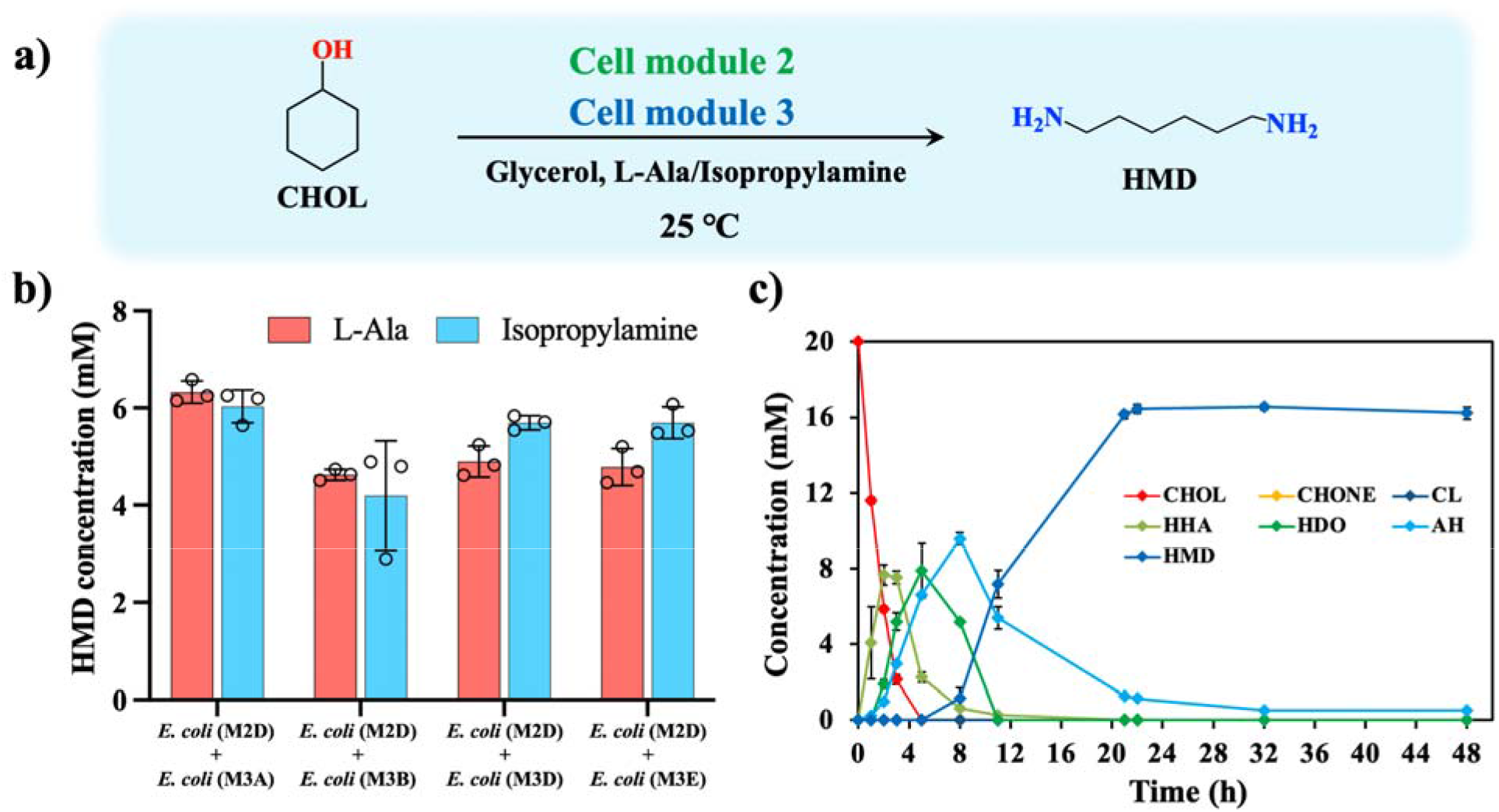
Construction of *E. coli* consortia for HMD production in a one-pot, one-step manner. a) Scheme of EC2_3-catalyzed conversion of CHOL to HMD in a one-pot, one-step manner. b) Comparison of different EC2_3 for the biotransformation of CHOL to HMD in a one-pot, one-step manner. Reaction conditions: *E. coli* cell modules 2 and 3 at a ratio of 4:3 was resuspended in phosphate buffer (pH 8.0, 100 mM) with a total cell density of 14 g CDW L^-1^, 20 mM CHOL, reactions were performed at 25 °C, 220 rpm for 22 h, cofactor NAD(P)H/ATP was provided by the *E. coli* cells using 68 mM glycerol as an energy source and 60 mM isopropylamine was employed as amine donor. c) Time course of EC2_3-catalyzed conversion of CHOL to HMD in one-pot, one-step manner. Reaction conditions: *E. coli* consortia 2_3 composed of *E. coli* (M2D) and *E. coli* (M3A) at a ratio of 2:1 was resuspended in phosphate buffer (pH 8.0, 100 mM) with a total cell density of 14 g CDW L^-1^, 20 mM CHOL, reactions were performed at 25 °C, 220 rpm for 48 h, cofactor NAD(P)H/ATP was provided by the *E. coli* cells using 102 mM glycerol as an energy source and 80 mM isopropylamine was added as amine donor.

Finally, to examine the scalability of the developed *E. coli* consortium-catalyzed HMD production from CHOL, a scale-up reaction was conducted in a 5 L fermenter with a 600 mL reaction mixture containing 20 mM CHOL in a one-pot, one-step manner. As presented in **Supplementary Fig. 11**, without further optimization, 13.5 mM HMD was produced with very little intermediate accumulation, which was comparable to the results obtained using shaking flasks. Therefore, to the best of our knowledge, our work offers the highest HMD production, which is over 4.5-fold higher than that offered by employing adipic acid as a substrate (3 mM for HMD)^4^, in a biological manner. In addition, there are almost no intermediates left after the overall reaction in our developed biocatalytic system. In contrast, when using adipic acid as a substrate, 70% 6-aminocaproic acid (6-ACA) intermediate remained in the reaction mixture^4^, which is not beneficial for downstream product separation.

### One-pot HMD Synthesis from CH

Having achieved efficient HMD synthesis from CHOL, we turned our attention to HMD synthesis using CH as a less expensive and more readily available starting material. The previously engineered *E. coli* expressing P450_BM3_19A12^36^ (for SDS-PAGE analysis **see Supplementary Fig. 12**) was used in cell module 1 for converting CH to CHOL. With the three optimally functioning cell modules 1, 2 and 3 in hand, the *E. coli* consortium 1_2_3 (EC1_2_3) was constructed to transform CH into HMD. Encouraged by the positive result from the EC2_3-catalyzed conversion of CHOL to HMD in a one-pot, one-step manner, the one-pot, one-step transformation of CH into HMD was first attempted. However, only 0.23 mM HMD was produced after reacting for 18 h (**Supplementary Fig. 13**). This low productivity might be attributed to the sensitivity of P450 to the poor oxygen transfer caused by the increased viscosity at a high cell density or the loss of P450 enzyme activity due to product inhibition. This hypothesis needs to be addressed in future studies. Thus, to minimize potentially adverse effects, biocatalytic cascade reactions of CH to HMD were performed in a one-pot, two-step approach. This is the combination of cell modules 1 and 2 for catalysing the first step of CH to CHOL, followed by the addition of module 3 for initiating the transformation of CHOL into HMD as the second step. Then, the reaction conditions, including the cell density and the ratio of the three modular cells, were optimized. We first rescreened the different combinations of *E. coli* (M1) and *E. coli* (M2D) with different module 3 types *(E. coli* (M3A) or *E. coli* (M3D)) (**Fig. 7b)**. The result showed that EC1_2_3 containing *E. coli* (M1), *E. coli* (M2D) and *E. coli* (M3D) at a ratio of 1:2:2 with a total cell density of 20 g CDW L^-1^ yielded 3 mM HMD after 32 h. Finally, the time course of EC1_2_3-catalyzed conversion of CH to HMD was carried out under the optimized conditions (**Supplementary Fig. 14 & 15**) in a one-pot, two-step approach. As shown in **Fig. 7c**, HDO was first produced and reached a maximum of 10 mM after 8 h. Subsequently, cell module 3 was added to transform CHOL into HMD, and the highest product concentration of 7.6 mM was obtained within 22 h, which is 33-fold higher than that of the one-pot, one-step process.

**Figure 7.**
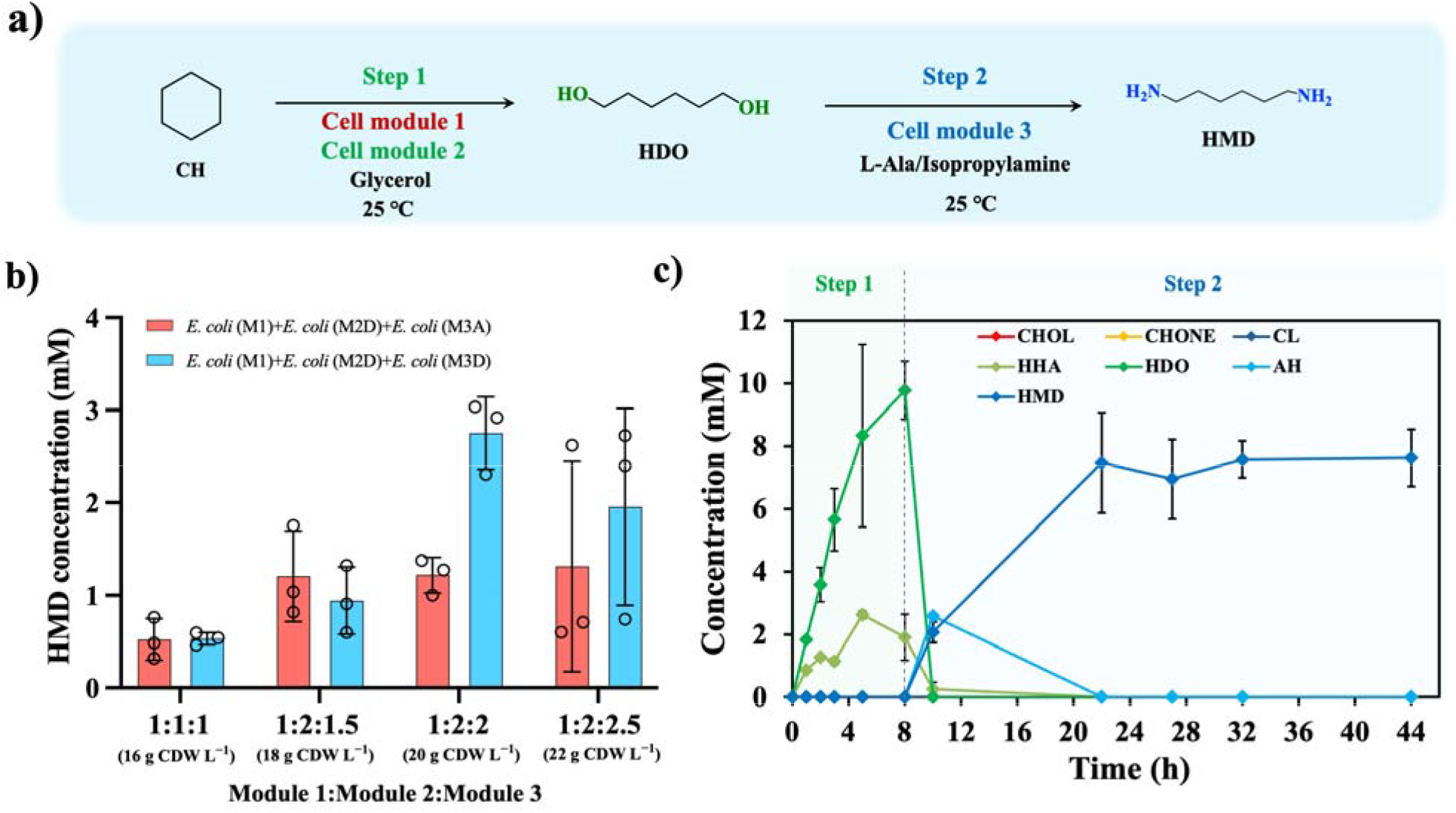
Construction of *E. coli* consortia for HMD production in a one-pot, two-step manner. a) Scheme of EC1_2_3-catalyzed conversion of CH to HMD in a one-pot, two-step manner. b) Cell density and module ratio for EC1_2_3-catalyzed conversion of CH to HMD in a one-pot, two-step manner. Reaction conditions: *E. coli* consortia consisting of modules 1, 2 and 3 at a designed ratio was resuspended in phosphate buffer (pH 8.0, 100 mM) with a specified cell density, 30 mM CH, reactions were performed at 25 °C, 220 rpm for 32 h, cofactor NAD(P)H/ATP was provided by the *E. coli* host cells using 136 mM glycerol as an energy source and 80 mM isopropylamine was employed as amine donor. Cell module 3 and amine donor isopropylamine were added after 7 h reaction. c) Time course of EC1_2_3 catalyzed conversion of CH to HMD in a one-pot-two-step manner. Reaction conditions: *E. coli* consortium consisting of *E. coli* (M1), *E. coli* (M2D) and *E. coli* (M3D) at a ratio of 3:3:4 was resuspended in phosphate buffer (pH 8.0, 100 mM) at a cell density of 20 g CDW L^-1^, 30 mM CH, reactions were performed at 25 °C, 220 rpm for 44 h, cofactor NAD(P)H/ATP was provided by the *E. coli* cells using 54 mM glycerol as an energy source and 80 mM isopropylamine was employed as amine donor. Cell module 3 and amine donor isopropylamine were added after 8 h reaction.

### Substrate Scope Investigation for the Synthesis of Different α,ω-diamines

In order to expand the substrate scope and demonstrate the generality of the developed *E. coli* consortia system, several cycloalkanols and cycloalkanes with varying carbon numbers (C6 to C8) were tested. The results are presented in **Fig. 8**. For all of the substrates (cycloalkanols or cycloalkanes), using the corresponding *E. coli* consortium as the catalyst, the respective diamines were produced with varying degrees of catalytic e□ciencv. For all substrates tested using EC2_3 as the catalyst, efficient DAs synthesis was achieved from the corresponding cycloalkanols. The product concentrations exceeded 12 mM with high product yields (72%~81%). For the EC1_2_3-catalyzed conversion of cycloalkanes to corresponding diamines in a one-pot, two-step approach, somewhat lower product concentrations (1.0-6.7 mM) were achieved, possibly due to the low catalytic efficiency in the first step of the P450-catalyzed reaction. The products were confirmed by GC□MS analysis (**Supplementary Figs. 16-21**). To the best of our knowledge, the biosynthesis of HMD, 1,7-heptanediamine or 1,8-octanediamine from either cycloalkanes or cycloalkanols have not been reported to date.

**Figure 8.**
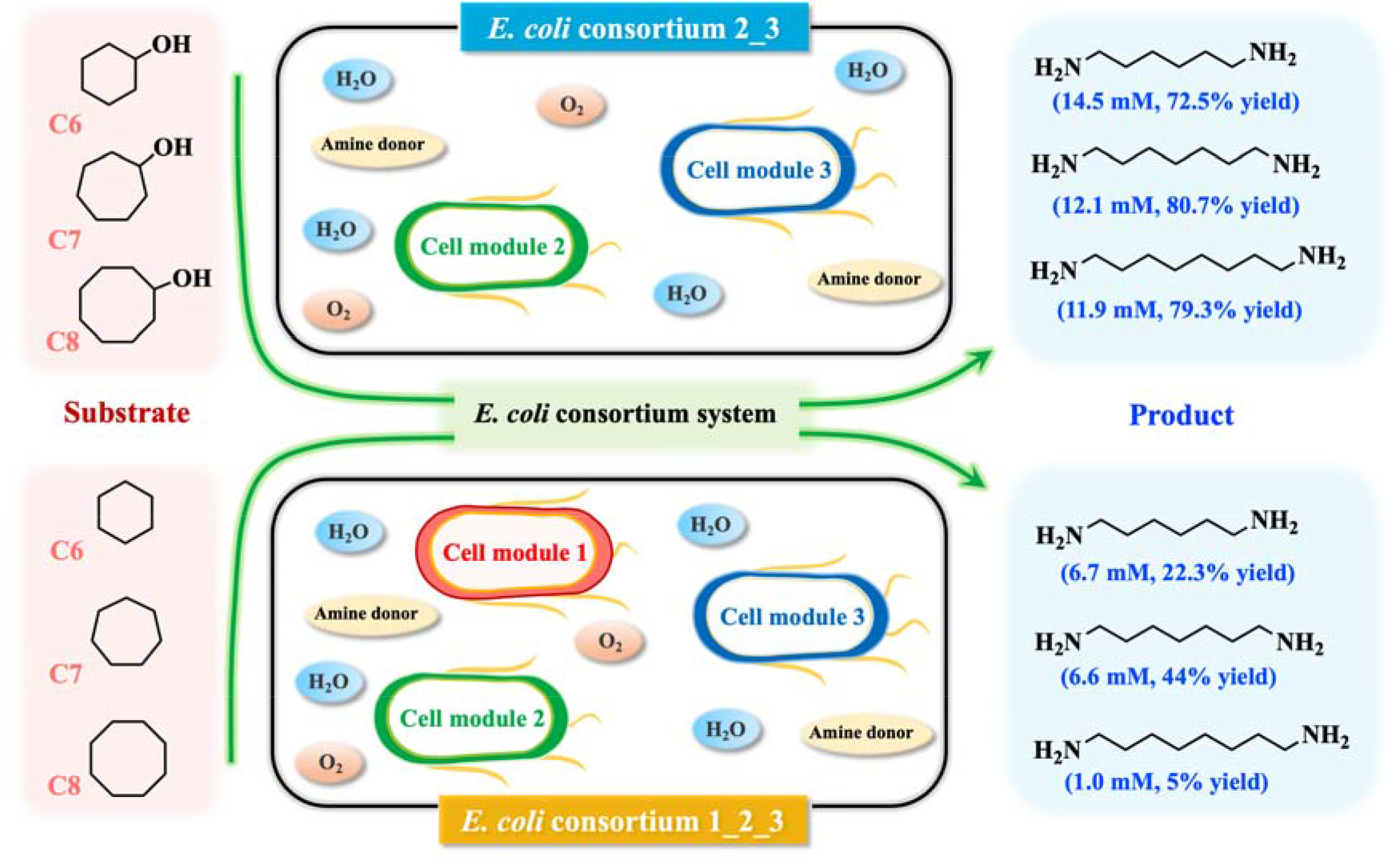
One-pot DAs production with designed *E. coli* consortia from cycloalkanes or cycloalkanols. The DAs production from the corresponding cycloalkanols was conducted in one-pot, one-step manner (reactions conditions see legend of Figure 6c), while for the DAs production from the respective cycloalkanes was conducted in one-pot, two-step process (reactions conditions see legend of Figure 7c).

## Conclusions

In summary, we have developed a general one-pot multienzyme biocatalytic cascade for α,ω-diamines production as important nylon monomers, starting from cheap and readily available cycloalkanes or cycloalkanols. This biocatalytic system was established with advanced techniques such as bioretrosynthesis tools, enzyme mining and microbial consortia-mediated pathway reconstruction. Using this strategy, the production of α,ω-diamines was successfully achieved, especially for the most prominent α,ω-diamine HMD. As high as 16.5 mM and 7.6 mM product concentrations were obtained when using CHOL and CH as substrates, respectively. To the best of our knowledge, this represents the highest HMD biosynthesis productivity known to date, providing an ideal solution to the problems encountered in industrial chemical processes. The current study also provides solutions and guidance for the further development of *in vivo* artificial biocatalytic cascades for challenging transformations. To further improve the efficiency of this biocatalytic system for DA production, future work will focus on engineering rate-limiting enzymes, such as P450, fine-tuning protein expression by promoter and RBS engineering, and scaling up bioreactor processes with precisely controlled parameters.

## Experimental section

### Construction of recombinant *E. coli* cells

DNA fragments of enzyme genes and the linear plasmid backbone were amplified by PCR using primers with 15 to 20 bp homologous arms that enabled the subsequent recombination. Full length of enzyme genes was assembled *via* overlap PCR and cloned into linear vector in the presence of T5 exonuclease to generate 15 bp or 20 bp sticky ends to promote recombination efficiency. Reaction mixtures were 5 μL containing linear vector, enzyme genes, buffer 4.0 (New England Biolabs) and T5 exonuclease, incubated in ice-water for 5 min, followed by quick addition of 50 μL competent cells (*E. coli* DH5α) for transformation and plating on LB agar containing appropriate antibiotics. Resulting transformants were picked and DNA sequenced for confirmation. Plasmids containing targeted enzyme genes were transformed into *E. coli* BL21 cells for protein expression and whole-cell biocatalyst preparation. The detailed information of the strain and plasmids, primers and synthetic gene sequences are listed in **Supplementary Tables 1-4**.

### Typical procedure for protein expression and preparation of whole-cell catalysts

Constructed *E. coli* cells were inoculated into 4 mL LB medium containing antibiotics (50 μg mL^-1^ kanamycin, 100 μg mL^-1^ ampicillin or both), and cultured at 37 °C, 220 rpm for 6 h. Precultures (4 mL) were transferred into 400 mL TB medium with appropriate antibiotics in 1 L shaking flasks and cultured at 37 °C, 220 rpm for 2~3 h until the OD_600_ reached 0.6~0.8, then IPTG was added to give a final concentration of 0.2 mM. The temperature was shifted to 25 °C for 14~16 h for protein expression. The cells were harvested by centrifugation at 5000 × *g*, 10 °C for 10 min, washed with 100 mM potassium phosphate buffer (pH 8.0) and used as whole-cell biocatalysts in the subsequent reactions.

### Gene editing in the *Escherichia coli* Genome via the CRISPR-Cas9 System

The ldhA gene on the genome of *E. coli* BL21 (DE3) was replaced by lactonase gene with a CRISPR-Cas9-mediated gene editing system by following the procedure reported previously^37^. Briefly, the system consisted of two plasmids (pCas and pTarget) and a donor DNA (lactonase gene with 500 bp upstream and downstream homologous arms of ldhA assembled via overlap PCR). Cell harbouring plasmid pCas was firstly cultured at 30 °C in the presence of kanamycin (50 mg/mL) and L-arabinose (20 mM) for λ-Red induction for 6 h and then the obtained cell was used to make competent cells. In addition, the pTarget and donor DNA were then co-transformed into *E. coli* BL21 (DE3) containing pCas by electroporation. After 3 h recovery at 30 °C, cells were spread onto LB agar (50 mg/mL kanamycin and 100 mg/mL spectinomycin) and incubated at 30 °C overnight, and the successful integration of lactonase gene into the host genome was identified by colony PCR and DNA sequencing. Finally, after a two-round plasmid curing, the engineered *E. coli* with lactonase inserted into the genome was obtained and used for subsequent experiments. The oligonucleotide sequences for genome engineering are given in **Supplementary Table 2**.

### Typical procedure for cell module 3 converting HDO to HMD

The substrate HDO (final concentration was 20 mM) was added to a 3 mL suspension of modular *E. coli* cells expressing enzymes of module 3 (final cell density was 8 g CDW L^-1^) in potassium phosphate buffer (0.1 M, pH 8.0) containing 100 mM isopropylamine as an amine donor. The reactions were performed at 25 °C, 220 rpm in 50 mL shaking flasks for a specified reaction time. Afterwards, samples were taken at appropriate intervals and prepared for HPLC analysis. In order to determine HMD concentration, each reaction sample was prepared for HPLC analysis with the HPLC-C18 column as follows: 208 μL of potassium phosphate buffer (0.1 M, pH 8.0) and 250 μL of dansyl chloride (dissolved in acetone, 6 mg/mL) containing 75 μL saturated NaHCO_3_ (pH=9.5) were added to 42 μL of the reaction sample, followed by ultrasonic treatment for 10 min and stand for 10 min. Subsequently, 500 mL methanol was added to this mixture. After filtration with a 0.22 μm membrane filter, reaction samples were analysed by HPLC. All experiments were performed in triplicate.

### Typical procedure for cell module 2 converting CHOL to HDO

The substrate CHOL (final concentration was 20 mM) was added to a 3 mL suspension of modular *E. coli* cells expressing enzymes of module 2 (final cell density was 6 g CDW L^-1^) in potassium phosphate buffer (0.1 M, pH 8.0) containing 68 mM glycerol to facilitate NADPH regeneration. The reactions were performed at 25 °C, 220 rpm in 50 mL shaking flasks for a specified reaction time. Afterwards, samples were taken at appropriate intervals and prepared for GC analysis. In order to determine HDO concentration, each reaction sample was prepared for GC analysis with the SH-Rtx-WAX column as follows: 375 μL of potassium phosphate buffer (0.1 M, pH 8.0) and500 μL of ethyl acetate containing 4 mM *n*-decane (internal standard) were added to 125 μL of the reaction sample saturated with NaCl, followed by vortexing and centrifugation (13,680 × *g*, 1 min). The organic phase was dried over anhydrous Na_2_SO_4_ and then directly used for GC analysis. All experiments were performed in triplicate.

### *E. coli* consortium 2_3-catalyzed transformation of CHOL to HMD

One-pot, one-step biosynthesis of HMD: For a 10 mL reaction system. The substrate CHOL (final concentration was 20 mM) was added to a 10 mL suspension of *E. coli* consortium 2_3 (final CDW was 24 g L^-1^, ratio of cell module 2 and cell module 3 was 2: 1) in potassium phosphate buffer (0.1 M, pH 8.0) containing 102 mM glycerol to facilitate NADPH regeneration and 80 mM isopropylamine as an amine group donor. The reactions were performed at 25 °C, 220 rpm in 250 mL shaking flasks. For the optimization experiments, the reaction system was 3 mL. All experiments were performed in triplicate, and the error bars indicate the standard deviations.

One-pot, two-step biosynthesis of HMD: The substrate CHOL (final concentration was 20 mM) was added to a 10 mL suspension of *E. coli* consortium 2_3 (final CDW was 14 g L ratio of cell module 2 and cell module 3 was 4: 3) in potassium phosphate buffer (0.1 M, pH 8.0) containing 68 mM glycerol to facilitate NADPH regeneration and 80 mM isopropylamine as an amine group donor. Among them, cell module 3 and isopropylamine were added after reaction at 25 □ for 8 h. The reactions were performed at 25 °C, 220 rpm in 250 mL shaking flasks. For the optimization experiments, the reaction system was 3 mL.

Samples were taken at appropriate intervals and prepared for HPLC analysis of HMD and AH as described in the section of typical procedure of cell module 3 converting HDO to HMD. In order to determine CHOL, CHONE, CL, HDO concentration, each reaction sample was prepared for GC analysis with the SH-Rtx-5 column as follows: 375 μL of potassium phosphate buffer (0.1 M, pH 8.0) and 500 μL of ethyl acetate containing 4 mM *n*-decane (internal standard) were added to 125 μL of the reaction sample saturated with NaCl, followed by vortexing and centrifugation (13,680 × *g*, 1 min). The organic phase was dried over anhydrous Na_2_SO_4_ and then directly used for GC analysis. In order to determine HHA concentration, each reaction sample was prepared for GC analysis with the SH-Rtx-5 column as follows: 375 μL of potassium phosphate buffer (0.1 M, pH 8.0), 50 μL HCl (4 M) and 500 μL of ethyl acetate were added to 125 μL of the reaction sample, followed by vortexing and centrifugation (13,680 × *g*, 1 min). The organic phase was dried over anhydrous Na_2_SO_4_ and followed with the derivatization step then used for GC analysis. All experiments were performed in triplicate.

### *E. coli* consortium 1_2_3-catalyzed transformation of CH to HMD

The substrate CH (final concentration was 30 mM) was added to a 10 mL suspension of *E. coli* consortium 1_2_3 (final CDW was 20 g L^-1^, ratio of module 1: 2: 3 was 3: 3: 4) in potassium phosphate buffer (0.1 M, pH 8.0) containing 68 mM glycerol to facilitate NADPH regeneration and 80 mM isopropylamine as an amine donor. Cell module 3 and isopropylamine were added after reaction at 25 □ for 8 h. The reactions were performed in a one-pot, two-step process at 25 °C, 220 rpm in 250 mL shaking flasks. Samples were taken at appropriate intervals and prepared for HPLC and GC analysis as described in the section of *E. coli* consortium 2_3-catalyzed transformation of CHOL to HMD. All experiments were performed in triplicate.

### Scale-up of *E. coli* consortium 2_3-catalyzed transformation of CHOL to HMD

The reaction was performed at 25 °C, 400 rpm in 5 L fermentor with the *E. coli* consortium 2_3 (final CDW was 20.8 g L^-1^, ratio of cell module 2: 3 was 2: 1) at a 600 mL reaction system. Other conditions were as described in the section of *E. coli* consortium 2_3-catalyzed transformation of CHOL to HMD (One-pot, one-step process). Samples were taken at appropriate intervals and prepared for HPLC and GC analysis as described in the section of *E. coli* consortium 2_3-catalyzed transformation of CHOL to HMD. All experiments were performed in triplicate.

### Typical procedure for *E. coli* consortium 2_3 converting cycloalkanol to α, ω-diamine

The conditions were as described in the section of *E. coli* consortium 2_3-catalyzed transformation of CHOL to HMD (One-pot-one-step). Samples were taken at appropriate intervals and prepared for HPLC and GC analysis as described in the section of *E. coli* consortium 2_3-catalyzed transformation of CHOL to HMD. All experiments were performed in triplicate.

### Typical procedure for *E. coli* consortium 1_2_3 converting cycloalkane to α, ω-diamine

The conditions were as described in the section of *E. coli* consortium 1_2_3-catalyzed transformation of CH to HMD. Samples were taken at appropriate intervals and prepared for HPLC and GC analysis as described in the section of *E. coli* consortium 2_3-catalyzed transformation of CHOL to HMD. All experiments were performed in triplicate.

### GC analysis

The procedure for GC analysis with the SH-Rtx-WAX column: the obtained mixtures were analyzed using the SH-Rtx-WAX column (30 m × 0.25 mm, 0.25 μm). The temperature program was as follows: 5 °C min^-1^ from 50 °C to 120 °C, 40 °C min^-1^ to 240 °C, and held at 240 °C for 3 min.

The procedure for GC analysis with the SH-Rtx-5 column: the obtained mixtures were analyzed using the SH-Rtx-5 column (30 m × 0.25 mm, 0.25 μm). The temperature program was as follows: For HHA analysis, 5 °C min^-1^ from 60 °C to 100 °C, 20 °C min^-1^ to 240 °C, and held at 240 °C for 1 min. For CHOL, CHONE, CL, HDO analysis, held at 80 °C for 3 min and then 12 °C min^-1^ from 80 °C to 165 °C and held at 165 °C for 1 min, 80 °C min^-1^ to 280 °C, and held at 280 °C for 2 min.

### Derivatization

The previously obtained mixtures (products in ethyl acetate) was centrifuged at 13,680×*g* for 10 min to remove Na_2_SO_4_ and then 200 μL supernatant solutions were transferred to new 1.5 mL tubes. The resulting solid was dissolved in 60 μL of pyridine and 30 μL of N-methyl-N-(trimethylsilyl) trifluoroacetamide (MSTFA) after ethyl acetate was evaporated completely. The derivatization reaction was performed at 65 °C for 1 h and then the mixtures were applied for GC analysis with the SH-Rtx-5 column to analyze HHA.

### HPLC analysis

The conversion analysis of aliphatic α, ω-amine alcohol and α, ω-diamine was performed *via* reversed phase HPLC technique using a Shimadzu LC-2030C Plus system equipped with a Shim-pack GIST C18 column (4.6 × 250 mm, 5 μm, SHIMADZU, Japan). The mobile phase consisted of 17% water (solvent A) and 83% acetonitrile (solvent B), flow rate was 0.8 mL/min. The injection volume was 10 μL and monitored absorption wavelength was 254 nm.

## Data availability

The data that support the plots in this paper and other study findings are available from the corresponding author upon request.

## Acknowledgments

This study was funded by the National Key Research and Development Program of China (2019YFA0905000), the Distinguished Young Scholars of Hubei Province (2020CFA072), and was Supported by the Innovation Base for Introducing Talents of Discipline of Hubei Province (2019BJH021) as well as the Research Program of State Key Laboratory of Biocatalysis and Enzyme Engineering.

## Author Contributions

A.L. conceived and supervised the project, Z.Z., L.F. and F.W. performed the experiments; Z.Z., L.F., F.W., Y.D. and Z. J. analysed the data; A.L., Y.D. and Z.J. wrote the manuscript; all authors checked and modified the manuscript.

## Additional information

Supplementary information is available in the online version of the paper. Correspondence and requests for materials should be addressed to A.L.

## References

1. Wang, L., Li, G. & Deng, Y. Diamine Biosynthesis: Research Progress and Application Prospects. Appl. Environ. Microbiol. 86, e01972–01920 (2020).

2. Sattler, J. H. et al. Redox self-sufficient biocatalyst network for the amination of primary alcohols. Angew. Chem. Int. Ed. Engl. 51, 9156–9159 (2012).

3. Alini, S. et al. The catalytic hydrogenation of adiponitrile to hexamethylenediamine over a rhodium/alumina catalyst in a three phase slurry reactor. J. Mol. Catal. A: Chem. 206, 363–370 (2003).

4. Fedorchuk, T. P. et al. One-Pot Biocatalytic Transformation of Adipic Acid to 6-Aminocaproic Acid and 1,6-Hexamethylenediamine Using Carboxylic Acid Reductases and Transaminases. J. Am. Chem. Soc. 142, 1038–1048 (2020).

5. Wang, F. et al. One-pot biocatalytic route from cycloalkanes to α,ω-dicarboxylic acids by designed *Escherichia coli* consortia. Nat. Commun. 11, 5035 (2020).

6. Li, Q., Zhang, Z., Zhao, J. & Li, A. Recent advances in the sustainable production of α,ω-C6 bifunctional compounds enabled by chemo-/biocatalysts. Green Chem. 24, 4270–4303 (2022).

7. Breikss, A. I. Process for Hydrocyanation. US5523453A (1996).

8. Rapoport, M. Hydrocyanation of Olefins. US4330483 (1982).

9. Yu, X., Li, H. & Deng, J. F. Selective hydrogenation of adiponitrile over a skeletal Ni–P amorphous catalyst (Raney Ni–P) at 1 atm pressure. Appl. Catal. A-Gen. 199, 191–198 (2000).

10. Pellegatta, J. L. et al. Catalytic investigation of rhodium nanoparticles in hydrogenation of benzene and phenylacetylene. J. Mol. Catal. A: Chem. 178, 55–61 (2002).

11. Mormul, J. et al. Synthesis of Adipic Acid, 1,6-Hexanediamine, and 1,6-Hexanediol via Double-n-Selective Hydroformylation of 1,3-Butadiene. Acs Catal 6, 2802–2810 (2016).

12. Wang, T., Ide, M. S., Nolan, M. R., Davis, R. J. & Shanks, B. H. Renewable Production of Nylon-6,6 Monomers from Biomass-Derived 5-Hydroxymethylfurfural (HMF). Energy Environment Focus 5, 13–17 (2016).

13. Harshal, C. A High Yield Route for the Production of Compounds from Renewable Sources. WO 2015/042201A2 (2015).

14. Zhang, Z. et al. One-pot biosynthesis of 1,6-hexanediol from cyclohexane by *de novo* designed cascade biocatalysis. Green Chem. 22, 7476–7483 (2020).

15. Song, W. et al. Asymmetric assembly of high-value alpha-functionalized organic acids using a biocatalytic chiral-group-resetting process. Nat. Commun. 9, 3818 (2018).

16. Wu, S. et al. Highly regio-and enantioselective multiple oxy-and amino-functionalizations of alkenes by modular cascade biocatalysis. Nat. Commun. 7, 11917 (2016).

17. Luo, Z. W. & Lee, S. Y. Biotransformation of *p*-xylene into terephthalic acid by engineered Escherichia coli. Nat. Commun. 8, 15689 (2017).

18. Benitez-Mateos, A. I., Roura Padrosa, D. & Paradisi, F. Multistep enzyme cascades as a route towards green and sustainable pharmaceutical syntheses. Nat. Chem. 14, 489–499 (2022).

19. Huffman, M. A. et al. Design of an in vitro biocatalytic cascade for the manufacture of islatravir. Science 366, 1255–1259 (2019).

20. Gao, D. et al. Efficient Production of L-homophenylalanine by Enzymatic-Chemical Cascade Catalysis. Angew. Chem. Int. Ed. Engl., e202207077 (2022).

21. Bachmann, B. O. Biosynthesis: is it time to go retro? Nat. Chem. Biol. 6, 390–393 (2010).

22. Delepine, B., Duigou, T., Carbonell, P. & Faulon, J. L. RetroPath2.0: A retrosynthesis workflow for metabolic engineers. Metab. Eng. 45, 158–170 (2018).

23. Finnigan, W., Hepworth, L. J., Flitsch, S. L. & Turner, N. J. RetroBioCat as a computer-aided synthesis planning tool for biocatalytic reactions and cascades. Nat Catal. 4, 98–104 (2021).

24. Bornscheuer, U. T. et al. Engineering the third wave of biocatalysis. Nature 485, 185–194 (2012).

25. Lovelock, S. L. et al. The road to fully programmable protein catalysis. Nature 606, 49–58 (2022).

26. Reetz, M. Making Enzymes Suitable for Organic Chemistry by Rational Protein Design. Chembiochem, e202200049 (2022).

27. Song, H., Ding, M. Z., Jia, X. Q., Ma, Q. & Yuan, Y. J. Synthetic microbial consortia: from systematic analysis to construction and applications. Chem. Soc. Rev. 43, 6954–6981 (2014).

28. Zhou, K., Qiao, K., Edgar, S. & Stephanopoulos, G. Distributing a metabolic pathway among a microbial consortium enhances production of natural products. Nat. Biotechnol. 33, 377–383 (2015).

29. Betke, T., Maier, M., Gruber-Wolfler, H. & Groger, H. Biocatalytic production of adiponitrile and related aliphatic linear alpha,omega-dinitriles. Nat. Commun. 9, 5112 (2018).

30. Cheng, Q., Thomas, S. M., Kostichka, K., Valentine, J. R. & Nagarajan, V. Genetic analysis of a gene cluster for cyclohexanol oxidation in *Acinetobacter* sp. strain SE19 by in vitro transposition. J. Bacteriol. 182, 4744–4751 (2000).

31. van der Vlugt-Bergmans, C. J. B. & van der Werf, M. J. Genetic and biochemical characterization of a novel monoterpene □-lactone hydrolase from *Rhodococcus erythropolis* DCL14. Appl. Environ. Microbiol. 67, 733–741 (2001).

32. Yoo, H. W. et al. Multi-enzymatic cascade reactions with Escherichia coli-based modules for synthesizing various bioplastic monomers from fatty acid methyl esters. Green Chem. 24, 2222–2231 (2022).

33. Opperman, D. J. & Reetz, M. T. Towards practical Baeyer-Villiger-monooxygenases: design of cyclohexanone monooxygenase mutants with enhanced oxidative stability. Chembiochem 11, 2589–2596 (2010).

34. Rodriguez, C. et al. Steric vs. electronic effects in the *Lactobacillus brevis* ADH-catalyzed bioreduction of ketones. Org Biomol Chem 12, 673–681 (2014).

35. Khusnutdinova, A. N. et al. Exploring bacterial carboxylate reductases for the reduction of bifunctional carboxylic acids. Biotechnol. J. 12, 1600751 (2017).

36. Yu, H. L. et al. Bioamination of alkane with ammonium by an artificially designed multienzyme cascade. Metab. Eng. 47, 184–189 (2018).

37. Jiang, Y. et al. Multigene editing in the *Escherichia coli* genome via the CRISPR-Cas9 system. Appl. Environ. Microbiol. 81, 2506–2514 (2015).

