## Supporting Information for "Transforming inert cycloalkanes into α,ω-diamines through designed enzymatic cascade catalysis"

| **Materials** |  |
| --- | --- |
| *Commercial materials* | S4 |
| *Strains and plasmids* | S4 |
| **Supplementary Tables** |  |
| ***Table S1****. Strains and plasmids* | S5 |
| ***Table S2****. Oligonucleotide sequences* | S5 |
| ***Table S3****. Transaminases* | S7 |
| ***Table S4.*** *Synthetic gene sequences* | S10 |
| **Supplementary Figures** |  |
| ***Figure S1.*** *Screening of the ω-transaminases* | S21 |
| ***Figure S2.*** *SDS-PAGE analysis of some high-performance transaminases (TAs) from the transaminases library expressed in E. coli* | S22 |
| ***Figure S3.*** *SDS-PAGE analysis of whole-cell proteins of cell module 3 expressed in E. coli* | S23 |
| ***Figure S4.*** *Previous work by our group using eight enzymes to synthesize HDO* | S23 |
| ***Figure S5.*** *Engineering of E. coli cell module 2 for the conversion of CHOL to HDO* | S24 |
| ***Figure S6.*** *SDS-PAGE analysis of whole-cell proteins of cell module 2 expressed in E. coli.* | S24 |
| ***Figure S7.*** *Optimization of experimental conditions for one-pot, two-step method with CHOL as substrate*  ***Figure S8.*** *Biocatalytic synthesis of cyclohexylamine from CHONE*  ***Figure S9.*** *GC analysis of cyclohexylamine during the conversion of CH or CHOL to HMD*  ***Figure S10.*** *Optimization of experimental conditions for one-pot, one-step method with CHOL as substrate*  ***Figure S11.*** *Biosynthesis of HMD from CHOL in 600 mL scale inone-pot, one-step process*  ***Figure S12.*** *SDS-PAGE analysis of whole-cell proteins of cell module 1 expressed in E. coli*  ***Figure S13.*** *Optimization of cell module ratio of E. coli (M1), E. coli (M2D) and E. coli (M3A) from CH to HMD in a one-pot, one-step process*  ***Figure S14.*** *Optimization of the cell module ratio of E. coli (M1) and E. coli (M2D) from CH to HMD in a one-pot, two-step process*  ***Figure S15.*** *Optimization of glycerol concentration from CH to HMD in a one-pot, two-step process*  ***Figure S16.*** *GC-MS analysis of reaction mixtures from E. coli consortium 2_3 catalyzed conversion of cyclohexanol to HMD*  ***Figure S17.*** *GC-MS analysis of reaction mixtures from E. coli consortium 1_2_3 catalyzed conversion of cyclohexane to HMD*  ***Figure S18.*** *GC-MS analysis of reaction mixtures from E. coli consortium 2_3 catalyzed conversion of cycloheptanol to 1,7-heptanediamine*  ***Figure S19****. GC-MS analysis of reaction mixtures from E. coli consortium 1_2_3 catalyzed conversion of cycloheptane to 1,7-heptanediamine* | S25  S26  S26  S27  S28  S28  S29  S29  S30  S31  S32  S33  S34 |
| ***Figure S20****. GC-MS analysis of reaction mixtures from E. coli consortium 2_3 catalyzed conversion of cyclooctanol to 1,8-octanediamine*  ***Figure S21****. GC-MS analysis of reaction mixtures from E. coli consortium 1_2_3 catalyzed conversion of cyclooctane to 1,8-octanediamine* | S35  S36 |
| ***References*** | S37 |

**Materials**

**Commercial materials.** Isopropyl *β*-D-1-thiogalactopyranoside (IPTG, >99%) and

required antibiotics were purchased from Sangon Biotech (Shanghai, China). Dansyl chloride, *n*-butanol, cyclohexanone, cyclohexylamine, 1,6-hexanediol and N-Methyl-

N-(trimethylsilyl) trifluoroacetamide with 1% trimethylchlorosilane were purchased from Macklin (Shanghai, China). Cycloheptane, cyclooctane, cyclohexanol, cycloheptanol, ε-caprolactone, 1,6-hexanediamine, 1,7-heptanediamine, 1,8-octanediamine, pyridine, isopropylamine and L-alanine were purchased from aladdin (Shanghai, China). Cyclooctanol and 6-amino-1-hexanol were purchased from 9 Ding Chemistry (Shanghai, China). Tryptone and yeast extract were purchased from OXOID (Shanghai, China). 6-Hydroxycaproic acid was purchased from Bidepharm (Shanghai, China). Primer STAR Max DNA Polymerase was purchased from Takara (Shanghai, China). T5 exonuclease and restriction enzymes were obtained from New England Biolabs (Beverley, MA, USA). Plasmid Miniprep Purification kit and DNA Clean/Extraction kit were purchased from Genemark (USA). Gene synthesis was performed by Wuhan GeneCreate Biological Engineering Co., Ltd. (Wuhan, China). Synthesis of oligonucleotides and DNA sequencing were performed by TSINGKE Biological Technology (Wuhan, China). All other chemicals were of chemical purity and commercially available from Sinopharm Chemical Reagent Co., Ltd (Shanghai, China).

**Strains and plasmids.** *E. coli* DH5α and *E. coli* BL21 (DE3) were used as hosts for

gene cloning and protein expression, respectively. The genes encoding alcohol

dehydrogenase (ADH) from *Lactobacillus brevis*^1^, lactonase from *Rhodococcus* sp. HI-31^2^, double mutant (C376L/M400I) of Baeyer-Villiger monooxygenase (BVMO) from *Acinetobacter* sp. NCIMB9871^3^, Carboxylic acid reductase (CAR)^4^ and Phosphopantetheinyl transferase (SFP)^4^ were chemically synthesized and ligated to plasmid pRSFDuet-1. Transaminases such as CV, PAKωTA, SPO3471, PP2159 and SAV2614 and alcohol dehydrogenases (ChnD) from *Acinetobacter sp.* NCIMB9871^5^ were chemically synthesized and ligated to plasmid pETDuet-1. P450BM3 variants 19A12^6^ was provided by professor Huilei Yu in East China University of Science and technology. Details of strains and plasmids used in this study are summarized in Table S1.

**Supplementary Tables**

**Supplementary Table 1**. Strains and plasmids

| **Strain** | **Description** | **Source** |
| --- | --- | --- |
| *E. coli* BL21 (DE3) | *F^–^ompT gal dcm lon hsdS_B_* (*r_B_^–^ m_B_^–^*) *λ(DE3 [lacI lacUV5-T7 gene 1 ind1 sam7 nin5])* | Invitrogen |

| **Plasmid** | **Description** | **Reference or source** |
| --- | --- | --- |
| M1 | pRSFDuet-1 carrying P450_BM3_19A12 | This study |
| M2A | pRSFDuet-1 carrying ADH, CAR, SFP and BVMO. Genome (IdHA**::**P_T7_-lactonase) | This study |
| M2B | pETDuet-1 carrying ADH, BVMO, CAR and SFP. Genome (IdHA**::**P_T7_-lactonase) | This study |
| M2C | pRSFDuet-1 carrying BVMO, ADH, CAR and SFP. Genome (IdHA**::**P_T7_-lactonase) | This study |
| M2D | pRSFDuet-1 carrying CAR, SFP, BVMO and ADH. Genome (IdHA**::**P_T7_-lactonase) | This study |
| M2E | pRSFDuet-1 carrying CAR, SFP, ADH and BVMO. Genome (IdHA**::**P_T7_-lactonase) | This study |
| M2F | pRSFDuet-1 carrying CAR-linker-SFP, ADH and BVMO. Genome (IdHA**::**P_T7_-lactonase) | This study |
| LBL | pRSFDuet-1 carrying ADH, BVMO, lactonase. | ^7^ |
| CSA | pRSFDuet-1 carrying CAR, SFP, AKR | ^7^ |
| M3A | pETDuet-1 carrying ChnD and CV | This study |
| M3B | pETDuet-1 carrying ChnD and PP2159 | This study |
| M3C | pETDuet-1 carrying ChnD and SAV2614 | This study |
| M3D | pETDuet-1 carrying ChnD and PAKωTA | This study |
| M3E | pETDuet-1 carrying ChnD and SPO3471 | This study |
| pETDuet-1 | Double T7 promoters, pBR322 ori, Amp^R^ | Novagen |
| pRSFDuet-1 | Double T7 promoters, RSF ori, Kan^R^ | Novagen |

**Supplementary Table 2**. Oligonucleotide sequences

| **Name** | **Sequence (5’ →3’)** |
| --- | --- |
| pETDuet-ChnD-CV-F | AAGCACGTGGTCTGGCATAAAGCCAGGATCCGAATTCGAGCTCG |
| pETDuet-ChnD-CV-R | TTATGCCAGACCACGTGCTTTCAGGGTC |
| pETDuet-ChnD-RA-F | AGCCAGGATCCGAATTCGAGCTCG |
| pETDuet-ChnD-RA-R | GATATATCTCCTTAGGTACCTTAGTTCTCGTGCATCAGAACG |
| SAV2614 to pETDuet-ChnD-OL-F | GGTACCTAAGGAGATATATCATGGGTAATCCGATTGCAGTGAGCAAAGATC |
| SAV2614 to pETDuet-ChnD-OL-R | CTCGAATTCGGATCCTGGCTTTACAGTTTGGTCCATGCTTCGGTCAGAACC |
| PP2159 to pETDuet-ChnD-OL-F | GGTACCTAAGGAGATATATCATGAGTGAACAGAATAGTCAGACCCTGGC |
| PP2159 to pETDuet-ChnD-OL-R | CTCGAATTCGGATCCTGGCTTTAGCGCACTGCTTCATAGGTCAGATCC |
| SPO3471 to pETDuet-ChnD-OL-F | GGTACCTAAGGAGATATATCATGGCCACCATTACCAATCATATGCCGAC |
| SPO3471 to pETDuet-ChnD-OL-R | CTCGAATTCGGATCCTGGCTTTAGGCGGCACTTTTCATCAGACCCTG |
| PAKωTA to pETDuet-ChnD-OL-F | GGTACCTAAGGAGATATATCATGAACAGTCAGATTACCAATGCAAAAACCCGC |
| PAKωTA to pETDuet-ChnD-OL-R | CTCGAATTCGGATCCTGGCTTTAGGCCAGAACTGCGGCGGC |
| PRSF-CSLB-RA-F | GGTACCTAAGGAGATATATCATGAGCAATCGTCTGGATGGTAAAGTTGC |
| PRSF-CSLB-RA-R | GTGCTAATGGTTTCGGTCATGTGGTGATGATGGTGATGGCTGC |
| PRSF-CSBL-RA-F | GGTACCTAAGGAGATATATCATGTCACAAAAAATGGATTTTGATGCTATCGTG |
| PRSF-CSBL-RA-R | GTGCTAATGGTTTCGGTCATGTGGTGATGATGGTGATGGCTGC |
| CAR-SFP to PRSF-CSLB/BL-OL-F | ATGACCGAAACCATTAGCACCGC |
| CAR-SFP to PRSF-CSLB/BL-OL-R | GATATATCTCCTTAGGTACCTTACAGCAGTTCTTCATAGCTAACCATGG |
| PRSF-LBCS-RA-F | GCTATGAAGAACTGCTGTAAGCCAGGATCCGAATTCGAGCTC |
| PRSF-LBCS-RA-R | GATATATCTCCTTAGGTACCTTAGGCATTGGCAGGTTGCTTGATATCTG |
| PRSF-BLCS-RA-F | GCTATGAAGAACTGCTGTAAGCCAGGATCCGAATTCGAGCTC |
| PRSF-BLCS-RA-R | GATATATCTCCTTAGGTACCTTACTGTGCGGTATAACCACCATCCAC |
| CAR-SFP to PRSF-LB/BLCS-OL-F | GGTACCTAAGGAGATATATCATGACCGAAACCATTAGCACCGC |
| CAR-SFP to PRSF-LB/BLCS-OL-R | TTACAGCAGTTCTTCATAGCTAACCATGGTAATATCTTC |
| PRSF-LCSB-RA-F | AGCAACCTGCCAATGCCTAAGCCAGGATCCGAATTCGAGCTCGG |
| PRSF-LCSB-RA-R | GATATATCTCCTTAGGTACCTTACTGTGCGGTATAACCACCATCCACAAC |
| CAR-SFP to PRSF-LCSB-OL-F | GGTACCTAAGGAGATATATCATGACCGAAACCATTAGCACCGCAGC |
| CAR-SFP to PRSF-LCSB-OL-R | TTAGGCATTGGCAGGTTGCTTGATATCTGAACG |
| pETDuet-RA-F | AAGACGTGTGGGCTGGGTAAAAGCTTGCGGCCGCATAATGCTTAAG |
| pETDuet-RA-R | GGCATTTCTTTAATTGCCATCGAATTCGGATCCTGGCTGTGGTG |
| PRSF-C-Linker-SLB-RA-F | GCTCTGGCGGACCCGGCTCTATGAAAATCTATGGCATCTATATGGATCGTCCGC |
| PRSF-C-Linker-SLB-RA-R | AGAGCCGGGTCCGCCAGAGCCTCCGCCGCCAACTAAACCCAGCAGCTGAATATCGCTGG |
| **Oligonucleotide sequences for genome engineering** | |
| **Homology A** |  |
| ldhA-A-F | TATAAGTTAATGTCTGTTTCGCGGTCGCC |
| ldhA-A-T7-R | GGGAGAGCGTCGAGATCCCGTCTTGCCGCTCCCCTGCAACC |
| **Amplify Lac** |  |
| T7-Lactonase-F | CGGGATCTCGACGCTCTCCC |
| Lactonase-T7ter-R | CCTGAGGTTTCAGCAAAAAACCCCTCAAG |
| **Homology B** |  |
| T7-ldhA-B-F | TTTTTTGCTGAAACCTCAGGAAGACTTTCTCCAGTGATGTTGAATCACATTTAAGC |
| ldhA-B-R | CAAGCAGAATCAAGTTCTACCATGCCGA |
| sgRNA for ldhA | CGAGTCCTTTGGCTTTGAGC |

**Supplementary Table 3.** **Sources of different transaminases**

| Number | Protein name | Gene name | Organism | Source |
| --- | --- | --- | --- | --- |
| T1 | PA5313 | PA5313 | *Pseudomonas aeruginosa* | Q9HTP1^a^ |
| T2 | SM2404 | SMc01534 | *Rhizobium meliloti* | Q92N34 ^a^ |
| T3 | ectB | ectB | *Bacillus clausii (strain KSM-K16)* | Q5WL78 ^a^ |
| T4 | ω-(R)-transaminase | XP_001209325 | *Aspergillus terreus NIH2624* | Q0C8G1 ^a^ |
| T5 | Rha04845 | gabT2 | *Rhodococcus jostii* | Q0S806 ^a^ |
| T6 | SAV2585 | gabT | *Streptomyces avermitilis* | Q82K21 ^a^ |
| T7 | SM5064 | SM_b20379 | *Rhizobium meliloti* | Q92WH4 ^a^ |
| T8 | PA4805 | PA4805 | *Pseudomonas aeruginosa* | Q9HV04 ^a^ |
| T9 | SM4420 | SMa1855 | *Rhizobium meliloti* | Q92Y66 ^a^ |
| T10 | TA-(T7C/S47C/Q380L/V379L/M166F/R416D/Y168A) | TA-(T7C/S47C/Q380L/V379L/M166F/R416D/Y168A) | *Chromobacterium violaceum* | (CN 110592042 B) ^b^ |
| T11 | SPO3471 | SPO3471 | *Ruegeria pomeroyi* | Q5LMU1 ^a^ |
| T12 | pyruvate transaminase [Vibrio fluvialis] |  | *Vibrio fluvialis* | F2XBU9 ^a^ |
| T13 | Pc22g00160 protein | PCH_Pc22g00160 | *Penicillium rubens Wisconsin* | B6HP76 ^a^ |
| T14 | SC5440 | SCO5676 | *Streptomyces coelicolor* | O86823 ^a^ |
| T15 | 3FCR_Y59W/Y87L/T231A/L382M/G429A (3FCR_WLAMA) | TM1040_2691 | *Ruegeria sp. TM1040* | Q1GD43 ^a^ |
| T16 | CV | CV_2025 | *Chromobacterium violaceum* | Q7NWG4 ^a^ |
| T17 | MAR0012 | Maqu_0007 | *Marinobacter hydrocarbonoclasticus* | A1TWJ6 ^a^ |
| T18 | HEWT | spuC (HELO 1904) | *Halomonas elongata DSM 2581* | E1V913 ^a^ |
| T19 | GabT | gabT | *Escherichia coli* | P22256 ^a^ |
| T20 | BAS4776 | GBAA_5138 | *Bacillus anthracis* | A0A0F7R517 ^a^ |
| T21 | SAV2614 | SAVERM_2612 | *Streptomyces avermitilis* | Q82JZ2 ^a^ |
| T22 | PAKω-TA | pakω-ta | *Pseudomonas aeruginosa PAK* | ARI05650.1 ^c^ |
| T23 | TA-WT | TA-WT | *Chromobacterium violaceum* | (CN 110592042 B) ^b^ |
| T24 | MLL7127 | mll7127 | *Mesorhizobium japonicum* | Q987B2 ^a^ |
| T25 | Rha07987 | RHA1_ro05386 | *Rhodococcus jostii* | Q0S5M0 ^a^ |
| T26 | PA0221 | PA0221 | *Pseudomonas aeruginosa* | Q9I6R7 ^a^ |
| T27 | PP2159 | spuC-I | *Pseudomonas putida* | Q88KV9 ^a^ |
| T28 | 2'-deamino-2'-hydroxyneamine transaminase | kacL | *Streptomyces kanamyceticus* | Q6L741 ^a^ |
| T29 | PatA | patA | *Escherichia coli* | P42588 ^a^ |
| **T30** | SM3293 | SMc04388 | *Rhizobium meliloti* | Q92L05 ^a^ |
| T31 | Branched-chain-amino-acid aminotransferase | ilvE | *Escherichia coli (strain K12)* | P0AB80 ^a^ |
| T32 | D-alanine aminotransferase | dat | *Bacillus sp. (strain YM-1)* | P19938 ^a^ |
| T33 | Aminotransferase | PH1371 | *Pyrococcus horikoshii (strain ATCC 700860 / DSM 12428 / JCM 9974 / NBRC 100139 / OT-3)* | O59096 ^a^ |
| T34 | Putative branched-chain amino acid aminotransferase protein | msi247 | *Mesorhizobium japonicum R7A* | Q8KJC1 ^a^ |
| T35 | Aromatic-amino-acid aminotransferase | tyrB | *Paracoccus denitrificans* | P95468 ^a^ |
| T36 | branched-chain amino acid aminotransferase, putative | AFUA_7G06900 | *Aspergillus fumigatus Af293* | Q4WH08 ^a^ |
| T37 | Aminotransferase | TTHA0940 | *Thermus thermophilus (strain HB8 / ATCC 27634 / DSM 579)* | Q5SJR8 ^a^ |
| T38 | aminotransferase, class IV | An04g08630 | *Aspergillus niger CBS 513.88* | A2QJX8 ^a^ |
| T39 | 3-oxo-glucose-6-phosphate:glutamate aminotransferase | ntdA | *Bacillus subtilis (strain 168)* | O07566 ^a^ |
| T40 | AT |  | *Aspergillus terreus* | XP_001209325.1 ^d^ |
| T41 | Putative alanine aminotransferase | SPBC582.08 | *Schizosaccharomyces pombe (strain 972 / ATCC 24843) (Fission yeast)* | Q10334 ^a^ |
| T42 | Aromatic-amino-acid aminotransferase 1 | OCC_04335 | *Thermococcus litoralis (strain ATCC 51850 / DSM 5473 / JCM 8560 / NS-C)* | H3ZPL1 ^a^ |
| T43 | branched chain amino acid: 2-keto-4-methylthiobutyrate aminotransferase |  | *Burkholderia cenocepacia HI2424* | ABK12047.1 ^c^ |
| T44 | PP3681 | PP_3718 | *Pseudomonas putida* | Q88GK3 ^a^ |
| T45 | hypothetical protein | AO090308000020 | *Aspergillus oryzae RIB40* | Q2UPF1 ^a^ |
| T46 | Branched-chain-amino-acid aminotransferase | ilvE | *Thermus thermophilus (strain HB8 / ATCC 27634 / DSM 579)* | Q5SM19 ^a^ |
| T47 | Branched-chain amino acid aminotransferase/4-amino-4-deoxychorismate lyase | SADFL11_359 | *Labrenzia alexandrii DFL-11* | A0A5E8GTH8 ^a^ |
| T48 | Histidinol-phosphate aminotransferase 2 | hisC2 | *Pseudomonas fluorescens (strain Pf0-1)* | Q3K8U2 ^a^ |
| T49 | aminotransferase class IV | HNE_2507 | *Hyphomonas neptunium* | Q0BZ92 ^a^ |
| T50 | Aminotransferase | Pden_3984 | *Paracoccus denitrificans (strain Pd 1222)* | A1B956 ^a^ |
| T51 | MULTISPECIES: aminotransferase class IV | Mvan_4516 | *Mycolicibacterium* | A1TDP1 ^a^ |
| T52 | branched-chain amino acid--2-keto-4-methylthiobutyrate aminotransferase | Jann_1122 | *Jannaschia sp. CCS1* | Q28TC3 ^a^ |
| T53 | branched-chain amino acid--2-keto-4-methylthiobutyrate aminotransferase | Bcep18194_C6784 | *Burkholderia lata* | Q39NY5 ^a^ |
| T54 | branched-chain amino acid aminotransferase | ilvE_3 | *Luminiphilus syltensis NOR5-1B* | B8KQT8 ^a^ |
| T55 | Beta-alanine--pyruvate aminotransferase | bauA | *Pseudomonas aeruginosa (strain ATCC 15692 / DSM 22644 / CIP 104116 / JCM 14847 / LMG 12228 / 1C / PRS 101 / PAO1)* | Q9I700 ^a^ |
| T56 | The (R)-selective transaminase from Nectria haematococca | NECHADRAFT_29730 | *Fusarium vanettenii (strain ATCC MYA-4622 / CBS 123669 / FGSC 9596 / NRRL 45880 / 77-13-4) (Fusarium solani subsp. pisi)* | C7YVL8 ^a^ |
| T57 | D-amino acid aminotransferase | mlr1594 | *Mesorhizobium japonicum* | Q98K82 ^a^ |
| T58 | Aminodeoxychorismate lyase | pabC | *Escherichia coli (strain K12)* | P28305 ^a^ |
| T59 | MesAT | N/A | *Mesorhizobium sp. LUK* | A3EYF7 ^a^ |

^a^ Uniport ID, ^b^ Chinese patent, ^c^ GenBank ID, ^d^ NCBI accession.

**Supplementary Table 4**. **Synthetic gene sequences**

**Alcohol dehydrogenase (ChnD) from *Acinetobacter* sp. NCIMB9871**

ATGCATTGTTATTGCGTTACCCATCATGGTCAGCCGCTGGAAGATGTTGAAAAAGAAATTCCGCAGCCGAAAGGCACCGAAGTTCTGCTGCATGTTAAAGCAGCAGGTCTGTGTCATACCGATCTGCATCTGTGGGAAGGTTATTATGATTTAGGTGGTGGTAAACGTCTGAGCCTGGCAGATCGTGGTCTGAAACCGCCTCTGACACTGAGCCATGAAATTACCGGTCAGGTTGTTGCAGTTGGTCCGGATGCAGAAAGCGTTAAAGTTGGTATGGTTAGCCTGGTTCATCCGTGGATTGGTTGTGGTGAATGTAATTATTGTAAACGCGGTGAAGAAAACCTGTGTGCAAAACCGCAGCAGCTGGGTATTGCAAAACCTGGTGGTTTTGCAGAATACATTATTGTTCCGCATCCGCGTTATCTGGTTGATATTGCAGGTCTGGATCTGGCCGAAGCAGCACCGCTGGCATGTGCCGGTGTTACCACCTATAGCGCACTGAAAAAATTCGGTGATCTGATTCAGAGCGAACCGGTTGTTATTATTGGTGCCGGTGGTCTGGGTCTGATGGCACTGGAACTGCTGAAAGCAATGCAGGCAAAAGGTGCAATTGTTGTGGATATCGATGATAGCAAACTGGAAGCAGCCCGTGCAGCCGGTGCACTGAGCGTGATTAATAGCCGTAGCGAAGATGCAGCACAGCAGCTGATTCAGGCCACCGATGGTGGTGCACGTCTGATTCTGGACCTGGTTGGTAGCAATCCGACACTGAGTCTGGCACTGGCAAGCGCAGCACGTGGTGGTCATATTGTTATTTGTGGCCTGATGGGTGGTGAAATCAAACTGAGCATTCCGGTTATTCCGATGCGTCCGCTGACCATTCAGGGTAGCTATGTTGGCACCGTTGAAGAACTGCGTGAACTGGTTGAGCTGGTTAAAGAAACCCATATGAGCGCAATTCCGGTGAAAAAACTGCCGATTAGCCAGATTAATAGTGCCTTTGGCGATCTGAAAGATGGTAATGTTATTGGTCGTATCGTTCTGATGCACGAGAACTAA

**CV from *Chromobacterium violaceum***

ATGCAGAAACAGCGTACCACCAGTCAGTGGCGTGAACTGGATGCAGCACATCATCTGCATCCGTTTACCGATACCGCAAGCCTGAATCAGGCAGGCGCACGTGTTATGACCCGTGGTGAAGGTGTTTATCTGTGGGATAGCGAAGGCAACAAAATTATCGATGGTATGGCAGGTCTGTGGTGTGTTAATGTTGGTTATGGTCGTAAAGATTTTGCAGAAGCAGCACGTCGTCAGATGGAAGAACTGCCGTTTTATAACACCTTTTTCAAAACCACACATCCGGCAGTTGTTGAACTGAGCAGCCTGCTGGCAGAAGTTACACCGGCAGGTTTTGATCGTGTGTTTTATACCAATAGCGGTAGCGAAAGCGTTGATACCATGATTCGTATGGTTCGTCGTTATTGGGATGTTCAGGGTAAACCGGAAAAAAAGACCCTGATTGGTCGTTGGAATGGTTATCATGGTAGCACCATTGGTGGTGCAAGCTTAGGTGGTATGAAATATATGCATGAACAGGGTGATCTGCCGATTCCTGGTATGGCCCATATTGAACAGCCGTGGTGGTATAAACATGGCAAAGATATGACACCGGATGAATTTGGTGTTGTTGCAGCACGTTGGCTGGAAGAAAAAATTCTGGAAATTGGTGCAGATAAAGTGGCAGCATTTGTTGGTGAACCGATTCAAGGTGCCGGTGGTGTTATTGTTCCGCCTGCAACCTATTGGCCTGAAATTGAACGTATTTGCCGTAAATATGACGTTCTGCTGGTTGCCGATGAAGTTATTTGCGGTTTTGGTCGTACCGGTGAATGGTTTGGTCATCAGCATTTTGGTTTTCAGCCGGACCTGTTTACCGCAGCAAAAGGTCTGAGCAGCGGTTATCTGCCTATTGGTGCCGTTTTTGTTGGTAAACGTGTTGCCGAAGGTCTGATTGCCGGTGGCGATTTTAATCATGGTTTTACCTATAGCGGTCATCCGGTTTGTGCAGCAGTTGCACATGCAAATGTTGCAGCCCTGCGTGATGAAGGTATTGTTCAGCGTGTTAAAGATGATATCGGTCCGTATATGCAAAAACGTTGGCGTGAAACCTTTAGCCGTTTTGAACATGTTGATGATGTTCGTGGTGTTGGTATGGTTCAGGCATTTACCCTGGTTAAAAACAAAGCAAAACGTGAACTGTTTCCGGATTTTGGTGAAATTGGCACCCTGTGTCGTGATATCTTTTTTCGCAATAACCTGATTATGCGTGCCTGCGGTGATCATATTGTTAGCGCACCGCCTCTGGTTATGACACGTGCCGAAGTTGATGAAATGCTGGCAGTTGCCGAACGCTGTCTGGAAGAATTTGAACAGACCCTGAAAGCACGTGGTCTGGCATAA

**PAKωTA**

ATGAACAGTCAGATTACCAATGCAAAAACCCGCGAATGGCAGGCACTGAGCCGTGATCATCATCTGCCGCCGTTTACCGATTATAAACAGCTGAATGAAAAGGGTGCCCGTATTATTACCAAAGCAGAAGGCGTTTATATTTGGGATAGCGAAGGTAATAAAATCCTGGATGCCATGGCCGGTCTGTGGTGTGTGAATGTTGGTTATGGCCGTGAAGAACTGGTTCAGGCCGCAACCCGTCAGATGCGCGAACTGCCGTTTTATAATCTGTTTTTTCAGACCGCACATCCGCCGGTTGTTGAACTGGCCAAAGCCATTGCCGATGTGGCCCCGGAAGGTATGAATCATGTTTTTTTTACCGGCAGCGGCAGCGAAGCAAATGATACCGTTCTGCGTATGGTTCGTCATTATTGGGCCACCAAAGGTCAGCCGCAGAAAAAAGTGGTGATTGGCCGCTGGAATGGCTATCATGGTAGCACCGTGGCCGGTGTTAGTCTGGGCGGCATGAAAGCCCTGCATGAACAGGGTGATTTTCCGATTCCGGGTATTGTGCATATTGCCCAGCCGTATTGGTATGGCGAAGGCGGTGATATGAGTCCGGATGAATTTGGCGTTTGGGCCGCAGAACAGCTGGAAAAAAAAATTCTGGAAGTGGGCGAAGAAAATGTGGCAGCATTTATTGCCGAACCGATTCAGGGCGCAGGCGGTGTGATTGTTCCGCCGGATACCTATTGGCCGAAAATTCGCGAAATTCTGGCAAAATATGATATTCTGTTCATCGCCGATGAAGTTATTTGCGGCTTTGGCCGCACCGGCGAATGGTTTGGCAGCCAGTATTATGGCAATGCACCGGATCTGATGCCGATTGCAAAAGGTCTGACCAGCGGTTATATTCCGATGGGCGGCGTTGTTGTGCGTGATGAAATTGTTGAAGTGCTGAATCAGGGTGGCGAATTTTATCATGGTTTTACCTATAGTGGCCATCCGGTTGCAGCAGCCGTTGCCCTGGAAAATATTCGCATTCTGCGCGAAGAAAAAATTATTGAAAAGGTGAAAGCCGAAACCGCCCCGTATCTGCAGAAACGTTGGCAGGAACTGGCCGATCATCCGCTGGTGGGTGAAGCACGTGGTGTTGGTATGGTTGCAGCACTGGAACTGGTTAAAAATAAAAAAACCCGTGAACGCTTTACCGATAAAGGTGTTGGCATGCTGTGTCGCGAACATTGCTTTCGCAATGGTCTGATTATGCGCGCCGTTGGCGATACCATGATTATTAGTCCGCCGCTGGTGATTGATCCGAGCCAGATTGATGAACTGATTACCCTGGCCCGTAAATGTCTGGATCAGACCGCCGCCGCAGTTCTGGCCTAA

**PP2159 from *Pseudomonas* *putida***

ATGAGTGAACAGAATAGTCAGACCCTGGCATGGCAGACCATGAGTCGTGATCATCATCTGGCACCGTTTAGCGATGTTCGTCAGCTGGCAGAAAAAGGTCCGCGCATTATTACCAGCGCCAAAGGTGTTTATCTGTGGGATAGCGAAGGTAATAAAATTCTGGATGGCATGGCCGGTCTGTGGTGCGTTGCAGTTGGTTATGGCCGTGAAGAACTGGCCGAAGTGGCAAGTCAGCAGATGAAACAGCTGCCGTATTATAATCTGTTTTTTCAGACCGCCCATCCGCCGGCCCTGGAACTGGCAAAAGCCATTGCCGAAGTTGCACCGCAGGGCATGAATCATGTTTTTTTTACCGGCAGCGGTAGTGAAGGTAATGATACCGTTCTGCGCATGGTGCGTCATTATTGGGCACTGAAAGGTCAGAAAAATAAAAAAGTGATCATCGGTCGTATCAATGGCTATCATGGTAGCACCGTTGCAGGTGCCGCACTGGGTGGTATGAGCGGTATGCATCAGCAGGGTGGCGTTATTCCGGATGTTGTGCATATTCCGCAGCCGTATTGGTTTGGTGAAGGCGGTGATATGACCGAAGCCGATTTTGGCGTTTGGGCCGCCGAACAGCTGGAAAAAAAAATTCTGGAAGTGGGTGTGGATAATGTGGCAGCCTTTATTGCAGAACCGATTCAGGGTGCAGGTGGCGTTATCATTCCGCCGCAGACCTATTGGCCGAAAATTAAAGAAATTCTGGCACGCTATGATATTCTGTTTGTTGCCGATGAAGTGATTTGCGGTTTTGGTCGCACCGGCGAATGGTTTGGTACCGATTATTATGATCTGAAACCGGATCTGATGACCATTGCAAAAGGTCTGACCAGCGGTTATATTCCGATGGGTGGCGTGATTGTGCGCGATGAAGTTGCCAAAGTTATTAGCGAAGGCGGTGACTTTAATCATGGTTTTACCTATAGCGGTCATCCGGTTGCAGCCGCCGTTGGCCTGGAAAATCTGCGTATTCTGCGCGATGAACAGATTATTCAGCAGGTTCATGATAAAACCGCCCCGTATCTGCAGCAGCGCCTGCGCGAACTGGCCGATCATCCGCTGGTGGGCGAAGTGCGCGGCCTGGGTATGCTGGGTGCCATTGAACTGGTGAAAGATAAAGCAACCCGCGCACGCCATGAAGGCAAAGGCGTGGGTATGATTTGCCGTCAGCATTGCTTTGATAATGGCCTGATTATGCGCGCAGTTGGTGATACCATGATTATTGCCCCGCCGCTGGTTATTAGCATTGAAGAAATTGATGAACTGGTGGAAAAAGCACGCAAATGCCTGGATCTGACCTATGAAGCAGTGCGCTAA

**SAV2614 from *Streptomyces avermitilis***

ATGGGTAATCCGATTGCAGTGAGCAAAGATCTGAGTCGTACCGCCTATGATCATCTGTGGATGCATTTTACCCGTATGAGCAGTTATGAAAATGCCCCGGTTCCGACCATTGTTCGTGGTGAAGGCACCTATATTTATGATGATAAAGGTAAACGCTACCTGGATGGCCTGAGCGGTCTGTTTGTTGTGCAGGCAGGCCATGGTCGCACCGAACTGGCCGAAACCGCATTTAAACAGGCACAGGAACTGGCCTTTTTTCCGGTGTGGAGTTATGCACATCCGAAAGCCGTTGAACTGGCCGAGCGCCTGGCCAATTATGCACCGGGTGATCTGAATAAAGTGTTTTTTACCACCGGTGGCGGCGAAGCCGTTGAAACCGCATGGAAACTGGCCAAACAGTATTTTAAACTGCAGGGCAAACCGACCAAATATAAAGTTATTAGCCGCGCCGTTGCATATCATGGTACCCCGCAGGGTGCCCTGAGCATTACCGGCCTGCCGGCACTGAAAGCACCGTTTGAACCGCTGGTGCCGGGCGCACATAAAGTGCCGAATACCAATATTTATCGCGCCCCGCTGTTTGGCGATGATCCGGAAGCCTTTGGTCGTTGGGCAGCCGATCAGATTGAACAGCAGATTCTGTTTGAAGGCCCGGAAACCGTTGCAGCCGTGTTTCTGGAACCGGTGCAGAATGCAGGTGGCTGTTTTCCGCCGCCGCCGGGTTATTTTCAGCGCGTGCGCGAAATTTGCGATCAGTATGATGTTCTGCTGGTTAGCGATGAAGTTATTTGCGCATTTGGCCGTCTGGGCACCATGTTTGCCTGCGATAAATTTGGCTATGTGCCGGATATGATTACCTGCGCCAAAGGTATGACCAGTGGTTATAGCCCGATTGGTGCCTGCATTGTGAGTGATCGCATTGCAGAACCGTTTTATAAAGGCGATAATACCTTTCTGCATGGTTATACCTTTGGTGGCCATCCGGTGAGCGCAGCAGTGGGTGTTGCCAATCTGGATCTGTTTGAACGTGAAGGTCTGAATCAGCATGTGCTGGATAATGAAAGTGCATTTCTGACCACCCTGCAGAAACTGCATGATCTGCCGATTGTTGGCGATGTTCGCGGTAATGGCTTTTTTTATGGCATTGAACTGGTTAAAGACAAAGCCACCAAAGAAACCTTTACCGATGAAGAAAGTGAACGTGTGCTGTATGGTTTTGTTAGCAAAAAACTGTTTGAGTATGGTCTGTATTGTCGCGCAGATGATCGTGGTGATCCGGTGATTCAGCTGAGCCCGCCGCTGATTAGCAATCAGAGTACCTTTGATGAAATTGAAAGCATTATCCGCCAGGTTCTGACCGAAGCATGGACCAAACTGTAA

**SPO3471 from *Ruegeria pomeroyi***

ATGGCCACCATTACCAATCATATGCCGACCGCCGAACTGCAGGCCCTGGATGCAGCCCATCATCTGCATCCGTTTAGCGCAAATAATGCACTGGGTGAAGAAGGTACCCGTGTGATTACCCGTGCACGCGGTGTGTGGCTGAATGATAGCGAAGGCGAAGAAATTCTGGATGCAATGGCCGGCCTGTGGTGTGTGAATATTGGCTATGGTCGCGATGAACTGGCAGAAGTTGCCGCACGTCAGATGCGTGAACTGCCGTATTATAATACCTTTTTTAAGACCACCCACGTGCCGGCAATTGCCCTGGCCCAGAAACTGGCCGAACTGGCCCCGGGCGATCTGAATCATGTGTTTTTTGCAGGTGGTGGTAGTGAAGCAAATGATACCAATATTCGCATGGTGCGTACCTATTGGCAGAATAAAGGCCAGCCGGAAAAAACCGTTATTATTAGCCGTAAAAACGCATATCATGGCAGTACCGTGGCCAGCAGCGCACTGGGTGGTATGGCCGGCATGCATGCACAGAGTGGCCTGATTCCGGATGTTCATCATATTAATCAGCCGAATTGGTGGGCAGAAGGCGGTGATATGGACCCTGAAGAATTTGGTCTGGCCCGTGCACGTGAACTGGAAGAAGCCATTCTGGAACTGGGCGAAAATCGTGTGGCAGCCTTTATTGCAGAACCGGTTCAGGGCGCCGGCGGTGTGATTGTGGCCCCTGATAGTTATTGGCCGGAAATTCAGCGCATTTGCGATAAATATGATATTCTGCTGATCGCAGATGAAGTGATTTGCGGCTTTGGCCGCACCGGCAATTGGTTTGGCACCCAGACCATGGGTATTCGTCCGCATATTATGACCATTGCAAAAGGTCTGAGCAGTGGTTATGCACCGATTGGCGGTAGTATTGTGTGCGATGAAGTGGCACATGTGATTGGTAAAGATGAATTTAATCACGGCTATACCTATAGCGGCCATCCGGTTGCCGCCGCCGTGGCATTAGAAAATCTGCGCATTCTGGAAGAAGAAAATATTCTGGATCATGTGCGCAATGTTGCAGCACCGTATCTGAAAGAAAAATGGGAAGCACTGACCGATCATCCGCTGGTGGGCGAAGCAAAAATTGTGGGTATGATGGCAAGTATTGCCCTGACCCCGAATAAAGCAAGCCGCGCAAAATTTGCCAGTGAACCGGGCACCATTGGCTATATTTGCCGCGAACGCTGTTTTGCAAATAATCTGATTATGCGCCATGTTGGTGATCGCATGATTATTAGCCCGCCGCTGGTGATTACCCCGGCAGAAATTGATGAAATGTTTGTTCGTATTCGTAAGAGTCTGGATGAAGCCCAGGCAGAAATTGAAAAACAGGGTCTGATGAAAAGTGCCGCCTAA

**Alcohol dehydrogenase (ADH) from *Lactobacillus brevis***

ATGAGCAATCGTCTGGATGGTAAAGTTGCAATTATTACCGGTGGCACCTTAGGTATTGGTCTGGCAATTGCAACCAAATTTGTTGAAGAGGGTGCCAAAGTTATGATTACCGGTCGTCATAGTGATGTTGGTGAAAAAGCAGCAAAAAGCGTTGGTACACCGGATCAGATTCAGTTTTTTCAGCATGATAGCAGTGATGAAGATGGTTGGACCAAACTGTTTGATGCAACCGAAAAAGCATTTGGTCCGGTTAGCACCCTGGTTAATAATGCAGGTATTGCAGTGAATAAGAGCGTTGAAGAAACCACCACCGCAGAATGGCGTAAACTGCTGGCAGTTAATCTGGATGGCGTTTTTTTTGGTACACGTCTGGGTATTCAGCGCATGAAAAACAAAGGTCTGGGTGCAAGCATTATCAACATGAGCAGCATTGAAGGTTTTGTTGGTGATCCGAGCCTGGGTGCATATAATGCAAGCAAAGGTGCAGTTCGTATTATGAGCAAAAGCGCAGCACTGGATTGTGCACTGAAAGATTATGATGTTCGTGTGAATACCGTTCATCCGGGTTATATCAAAACACCGCTGGTTGATGATCTGCCTGGTGCCGAAGAAGCAATGAGCCAGCGTACAAAAACCCCGATGGGTCATATTGGTGAACCGAATGATATTGCCTATATCTGTGTTTATCTGGCCAGCAACGAAAGTAAATTTGCAACCGGTAGCGAATTTGTTGTGGATGGTGGTTATACCGCACAGTAA

**Baeyer-Villiger monooxygenase (BVMO) from *Acinetobacter* sp. NCIMB9871**

atgtcacaaaaaatggattttgatgctatcgtgattggtggtggttttggcggactttatgcagtcaaaaaattaagagacgagctcgaacttaaggttcaggcttttgataaagccacggatgtcgcaggtacttggtactggaaccgttacccaggtgcattgacggatacagaaacccacctctactgctattcttgggataaagaattactacaatcgctagaaatcaagaaaaaatatgtgcaaggccctgatgtacgcaagtatttacagcaagtggctgaaaagcatgatttaaagaagagctatcaattcaataccgcggttcaatcggctcattacaacgaagcagatgccttgtgggaagtcaccactgaatatggtgataagtacacggcgcgtttcctcatcactgctttaggcttattgtctgcgcctaacttgccaaacatcaaaggcattaatcagtttaaaggtgagctgcatcataccagccgctggccagatgacgtaagttttgaaggtaaacgtgtcggcgtgattggtacgggttccaccggtgttcaggttattacggctgtggcacctctggctaaacacctcactgtcttccagcgttctgcacaatacagcgttccaattggcaatgatccactgtctgaagaagatgttaaaaagatcaaagacaattatgacaaaatttgggatggtgtatggaattcagcccttgcctttggcctgaatgaaagcacagtgccagcaatgagcgtatcagctgaagaacgcaaggcagtttttgaaaaggcatggcaaacaggtggcggtttccgtttcatgtttgaaactttcggtgatattgccaccaatatggaagccaatatcgaagcgcaaaatttcattaagggtaaaattgctgaaatcgtcaaagatccagccattgcacagaagcttatgccacaggatttgtatgcaaaacgtccgttgtgtgacagtggttactacaacacctttaaccgtgacaatgtccgtttagaagatgtgaaagccaatccgattgttgaaattaccgaaaacggtgtgaaactcgaaaatggcgatttcgttgaattagacatgctgataCTGgccacaggttttgatgccgtcgatggcaactatgtgcgcatggacattcaaggtaaaaacggcttggccATTaaagactactggaaagaaggtccgtcgagctatatgggtgtcaccgtaaataactatccaaacatgttcatggtgcttggaccgaatggcccgtttaccaacctgccgccatcaattgaatcacaggtggaatggatcagtgataccattcaatacacggttgaaaacaatgttgaatccattgaagcgacaaaagaagcggaagaacaatggactcaaacttgcgccaatattgcggaaatgaccttattccctaaagcgcaatcctggatttttggtgcgaatatcccgggcaagaaaaacacggtttacttctatctcggtggtttaaaagaatatcgcagtgcgctagccaactgcaaaaaccatgcctatgaaggttttgatattcaattacaacgttcagatatcaagcaacctgccaatgcctaa

**Lactonase from *Rhodococcus* sp. HI-31**

ATGACCAATATTAGCGAAACCCTGAGCACCGCACCTGGTGGTGCAGCAGGTCCGGATGTTCTGCGTGATCTGTATGCAGATTGGAGCGAAATTATGGCAGCAACACCGGATCTGACCATTCGTCTGCTGCGTAGCCTGTTTGATGAATGGCATCAGCCGACCGTTGAACCGGAAGGTGTTACCTATCGTGAAGAAACCGTTGGTGGTGTTCCTGGTATTTGGTGTCTGCCGCAGGGTGCAGATGGTAGCAAAGTTCTGCTGTATACCCATGGTGGTGGTTTTGCAGTTGGTAGCGCAGCAAGCCATCGTAAACTGGCAGGTCATGTTGCAAAAGCACTGGGTGCCGTTGGTTTTGTTCTGGATTATCGTCGTGCACCGGAATTTCAGCATCCGGCACAGATTGAAGATGGTGTTGCAGCATTTGATGCACTGGTTGCAAATGGTATTGCACCGCAGGATATTACCACCATTGGTGATAGTGCCGGTGGTAATCTGGCAGTTGCAATTGCCCTGAGCCTGCGTGAACAGGGTAAACAAGGTCCGGGTAGCGTTATTGCATTTAGCCCGTGGCTGGATATGGAAAATAAAGGTGAAACCCTGGCCACCAATAATGATACCGATGCACTGATTACACCGGAACTGCTGGAAGGCATGATTGCCGGTGTGCTGGGTGATACCATTGATCCGAAAACACCGCTGGCAAATCCGCTGTATGCCGATTTTACCGGTTTTCCGCGTCTGTATATCACCGCAGGTAGCGTTGAAAGCCTGCTGGATAATGCAACCCGTCTGGAAAAATTAGCAGCATCTGCCGGTGTTGATGTTACCCTGAGTATTGGTGAAGGTCAGCAGCATGTTTATCCGTTTCTGGCAGGCCGTAGCGCACTGGTGGATGATGAATTTGCAAAGCTGGCAGCATGGTATCAGAAAGCCCTGGAATAA

**Carboxylic acid reductase from *Mycobacterium abscessus* ATCC 19977**

ATGACCGAAACCATTAGCACCGCAGCAGTTCCGACCACCGATCTGGAAGAACAGGTTAAACGTCGTATTGAACAGGTTGTTAGCAATGATCCGCAGCTGGCAGCCCTGCTGCCGGAAGATAGCGTTACCGAAGCAGTTAATGAACCGGATCTGCCGCTGGTTGAAGTTATTCGTCGTCTGCTGGAAGGTTATGGTGATCGTCCGGCACTGGGTCAGCGTGCATTTGAATTTGTTACCGGTGATGATGGTGCAACCGTTATTGCACTGAAACCGGAATACACCACCGTTAGCTATCGTGAACTGTGGGAACGTGCCGAAGCAATTGCAGCAGCATGGCATGAACAGGGTATTCGTGATGGTGATTTTGTTGCACAGCTGGGTTTTACCAGCACCGATTTTGCAAGCCTGGATGTTGCAGGTCTGCGTCTGGGTACAGTTAGCGTTCCGCTGCAGACCGGTGCCAGCCTGCAGCAGCGTAATGCAATCCTGGAAGAAACCCGTCCGGCAGTTTTTGCAGCAAGCATTGAATATCTGGATGCAGCAGTTGATAGCGTTCTGGCAACCCCGAGCGTGCGTCTGCTGAGCGTTTTTGATTATCATGCCGAAGTGGATAGCCAGCGTGAAGCACTGGAAGCAGTTCGTGCACGTCTGGAAAGCGCAGGTCGTACCATTGTTGTTGAAGCCCTGGCCGAAGCGCTGGCACGTGGTCGTGATCTGCCAGCAGCACCGCTGCCGAGTGCCGATCCGGATGCACTGCGCCTGCTGATTTATACCAGCGGTAGCACCGGTACACCGAAAGGTGCCATGTATCCGCAGTGGCTGGTTGCAAATCTGTGGCAGAAAAAATGGCTGACCGATGATGTTATTCCGAGCATTGGTGTGAATTTTATGCCGATGAGCCATCTGGCAGGTCGTCTGACCCTGATGGGCACCCTGAGCGGTGGTGGCACCGCCTATTATATCGCAAGCAGCGATCTGAGCACCTTTTTTGAAGATATTGCCCTGATTCGTCCGAGCGAAGTTCTGTTTGTTCCGCGTGTTGTGGAAATGGTTTTTCAGCGTTTTCAGGCAGAACTGGATCGTAGCCTGGCTCCGGGTGAAAGCAATAGCGAAATTGCAGAACGTATTAAAGTGCGTATTCGCGAACAGGATTTTGGTGGTCGTGTGCTGAGCGCAGGTAGCGGTAGTGCACCGCTGAGTCCGGAAATGACCGAATTTATGGAAAGCCTGCTGCAAGTGCCGCTGCGTGATGGTTATGGTAGTACCGAAGCCGGTGGTGTTTGGCGTGATGGCGTTCTGCAGCGTCCGCCTGTTACCGATTATAAACTGGTTGATGTGCCGGAACTGGGTTATTTTACCACCGATAGTCCGCATCCGCGTGGTGAACTGCGTCTGAAAAGCGAAACCATGTTTCCGGGTTATTACAAACGTCCGGAAACCACCGCAGATGTTTTTGATGATGAAGGCTATTACAAAACGGGTGATGTGGTTGCTGAACTGGGTCCTGATCATCTGAAATACCTGGATCGTGTTAAAAACGTTCTGAAACTGGCACAGGGTGAATTTGTGGCAGTTAGCAAACTGGAAGCCGCATATACCGGTAGTCCGCTGGTTCGTCAGATTTTTGTTTATGGTAATAGCGAACGTAGCTTTCTGCTGGCCGTTGTTGTTCCGACACCGGAAGTTCTGGAACGTTATGCAGATAGTCCGGATGCGCTGAAACCGCTGATTCAGGATAGTCTGCAGCAGGTTGCAAAAGATGCAGAACTGCAGAGCTATGAAATTCCGCGTGATTTTATTGTTGAAACCGTTCCGTTTACCGTTGAAAGCGGACTGCTGAGTGATGCACGTAAACTGTTACGTCCGAAACTGAAAGATCATTATGGTGAACGCCTGGAAGCCCTGTATGCCGAACTGGCAGAAAGCCAGAATGAACGTCTGCGTCAGCTGGCACGCGAAGCAGCAACACGTCCGGTTCTGGAAACCGTTACCGATGCCGCAGCCGCACTGCTGGGTGCAAGCAGCTCCGATCTGGCACCAGATGTTCGTTTTATTGATTTAGGTGGTGATAGCCTGAGCGCACTGAGCTATAGCGAGCTGCTGCGCGATATTTTTGAAGTTGATGTTCCGGTTGGTGTGATTAATAGCGTTGCAAATGATCTGGCAGCAATTGCCCGTCATATTGAAGCACAGCGTACAGGTGCAGCAACCCAGCCGACCTTTGCAAGCGTTCATGGTAAAGATGCCACCGTTATTACCGCAGGCGAACTGACCCTGGATAAATTTCTGGATGAAAGTCTGCTGAAAGCAGCCAAAGATGTTCAGCCTGCGACAGCAGATGTTAAAACCGTGCTGGTGACCGGTGGTAATGGCTGGTTAGGTCGTTGGCTGGTTCTGGATTGGCTGGAACGTCTGGCACCGAATGGTGGTAAAGTTTATGCACTGATTCGTGGTGCAGATGCGGAAGCAGCACGCGCACGCCTGGATGCCGTTTATGAAAGCGGTGATCCTAAACTGAGTGCACATTATCGTCAACTGGCCCAGCAGAGCCTGGAAGTTATTGCAGGCGATTTTGGCGATCAGGATCTGGGTCTGAGCCAAGAAGTTTGGCAGAAACTGGCGAAAGATGTTGATCTGATTGTTCATAGCGGTGCCCTGGTTAATCATGTTCTGCCGTATAGCCAGCTGTTTGGTCCGAATGTTGCCGGTACAGCAGAAATTATCAAACTGGCAATTAGCGAACGCCTGAAACCTGTTACCTATCTGAGTACCGTTGGTATTGCAGATCAGATTCCGGTTACCGAATTTGAAGAGGATAGTGATGTTCGCGTTATGAGCGCAGAACGTCAGATTAATGATGGCTATGCAAATGGCTATGGCAATAGCAAATGGGCTGGTGAAGTTCTGCTGCGTGAAGCCCATGATTTAGCCGGTCTGCCGGTTCGTGTTTTTCGTAGCGATATGATTCTGGCACATAGCGATTATCACGGTCAGCTGAATGTGACCGATGTTTTTACCCGTAGCATTCAGAGTCTGCTGCTGACAGGTGTTGCACCGGCAAGTTTTTATGAACTGGATGCGGATGGTAATCGCCAGCGTGCCCATTATGATGGTGTTCCAGGTGATTTTACCGCAGCCAGCATTACCGCAATTGGTGGTGTTAATGTTGTGGATGGTTATCGCAGCTTTGATGTGTTTAATCCGCATCACGATGGTGTTAGCATGGATACCTTTGTTGATTGGCTGATTGATGCCGGTTACAAAATTGCACGCATCGATGATTATGATCAGTGGTTAGCACGTTTTGAACTGGCCCTGAAAGGCCTGCCTGAACAGCAGCGTCAGCAGAGCGTTCTGCCACTGCTGAAAATGTATGAAAAACCGCAGCCTGCAATTGATGGTAGCGCACTGCCGACCGCAGAATTTAGCCGTGCAGTTCATGAAGCAAAAGTGGGTGATAGTGGTGAAATCCCGCATGTTACCAAAGAACTGATTCTGAAATATGCCAGCGATATTCAGCTGCTGGGTTTAGTTTAA

**Phosphopantetheinyl transferase from *Bacillus subtilis***

ATGAAAATCTATGGCATCTATATGGATCGTCCGCTGAGCCAAGAAGAAAACGAACGTTTTATGAGCTTTATCAGTCCGGAAAAACGTGAAAAATGCCGTCGCTTTTATCACAAAGAAGATGCACATCGTACCCTGCTGGGTGATGTTCTGGTTCGTAGCGTTATTAGCCGTCAGTATCAGCTGGATAAATCCGATATTCGTTTTAGCACCCAAGAATATGGTAAACCGTGTATTCCGGATCTGCCGGATGCACATTTTAACATTAGCCATAGCGGTCGTTGGGTTATTTGTGCATTTGATAGCCAGCCGATTGGCATTGATATCGAAAAAACCAAACCGATCAGCCTGGAAATTGCCAAACGTTTTTTTAGCAAAACCGAGTATAGCGATCTGCTGGCCAAAGATAAAGATGAACAGACCGATTATTTCTATCATCTGTGGTCCATGAAAGAGAGCTTCATTAAACAAGAAGGTAAAGGTCTGAGCCTGCCGCTGGATAGCTTTAGCGTTCGTCTGCATCAGGATGGTCAGGTTAGCATTGAACTGCCGGATAGCCATAGTCCGTGTTATATCAAAACCTATGAAGTTGATCCGGGTTACAAAATGGCAGTTTGTGCAGCACATCCGGATTTTCCGGAAGATATTACCATGGTTAGCTATGAAGAACTGCTGTAA

**P450_BM3_19A12**

ATGGCAATTAAAGAAATGCCTCAGCCAAAAACGTTTGGAGAGCTTAAAAATTTACCGTTATTAAACACAGATAAACCGGTTCAAGCTTTGATGAAAATTGCGGATGAATTAGGAGAAATCTTTAAATTCGAGGCGCCTGGTCGTGTAACGCGCTACTTATCAAGTCAGCGTCTAATTAAAGAAGCATGCGATGAATCACGCTTTGATAAAAACTTAAGTCAAGCGCTTAAATTTGTACGTGATTTTTTCGGAGACGGGTTATTTACAAGCTGGACGCATGAAAAAAATTGGAAAAAAGCGCATAATATCTTACTTCCAAGCTTCAGTCAGCAGGCAATGAAAGGCTATCATGCGATGATGGTCGATATCGCCGTGCAGCTTGTTCAAAAGTGGGAGCGTCTAAATGCAGATGAGCATATTGAAGTACCGGAAGACATGACACGTTTAACGCTTGATACAATTGGTCTTTGCGGCTTTAACTATCGCTTTAACAGCTTTTACCGAGATCAGCCTCATCCATTTATTACAAGTATGGTCCGTGCACTGGATGAAGCAATGAACAAGCTGCAGCGAGCAAATCCAGACGACCCAGCTTATGATGAAAACAAGCGCCAGTTTCAAGAAGATATCAAGGTGATGAACGACCTAGTAGATAAAATTATTGCAGATCGCAAAGCAAGCGGTGAACAAAGCGATGATTTATTAACGCATATGCTAAACGGAAAAGATCCAGAAACGGGTGAGCCGCTTGATGACGAGAACATTCGCTATCAAATTATTACATTCTTAATTGCGGGACACGAAACAACAAGTGGTCTTTTATCATTTGCGCTGTATTTCTTAGTGAAAAATCCACATGTATTACAAAAAGCAGCAGAAGAAGCAGCACGAGTTCTAGTAGATCCTGTTCCAAGCTACAAACAAGTCAAACAGCTTAAATATGTCGGCATGGTCTTAAACGAAGCGCTGCGCTTATGGCCAACTTTTCCTGCGTTTTCCCTATATGCAAAAGAAGATACGGTGCTTGGAGGAGAATATCCTTTAGAAAAAGGCGACGAACTAATGGTTCTGATTCCTCAGCTTCACCGTGATAAAACAATTTGGGGAGACGATGTGGAAGAGTTCCGTCCAGAGCGTTTTGAAAATCCAAGTGCGATTCCGCAGCATGCGTTTAAACCGTTTGGAAACGGTCAGCGTGCGTGTATCGGTCAGCAGTTCGCTCTTCATGAAGCAACGCTGGTACTTGGTATGATGCTAAAACACTTTGACTTTGAAGATCATACAAACTACGAGCTGGATATTAAAGAAACTTTAACGTTAAAACCTGAAGGCTTTGTGGTAAAAGCAAAATCGAAAAAAATTCCGCTTGGCGGTATTCCTTCACCTAGCACTGAACAGTCTGCTAAAAAAGTACGCAAAAAGGCAGAAAACGCTCATAATACGCCGCTGCTTGTGCTATACGGTTCAAATATGGGAACAGCTGAAGGAACGGCGCGTGATTTAGCAGATATTGCAATGAGCAAAGGATTTGCACCGCAGGTCGCAACGCTTGATTCACACGCCGGAAATCTTCCGCGCGAAGGAGCTGTATTAATTGTAACGGCGTCTTATAACGGTCATCCGCCTGATAACGCAAAGCAATTTGTCGACTGGTTAGACCAAGCGTCTGCTGATGAAGTAAAAGGCGTTCGCTACTCCGTATTTGGATGCGGCGATAAAAACTGGGCTACTACGTATCAAAAAGTGCCTGCTTTTATCGATGAAACGCTTGCCGCTAAAGGGGCAGAAAACATCGCTGACCGCGGTGAAGCAGATGCAAGCGACGACTTTGAAGGCACATATGAAGAATGGCGTGAACATATGTGGAGTGACGTAGCAGCCTACTTTAACCTCGACATTGAAAACAGTGAAGATAATAAATCTACTCTTTCACTTCAATTTGTCGACAGCGCCGCGGATATGCCGCTTGCGAAAATGCACGGTGCGTTTTCAACGAACGTCGTAGCAAGCAAAGAACTTCAACAGCCAGGCAGTGCACGAAGCACGCGACATCTTGAAATTGAACTTCCAAAAGAAGCTTCTTATCAAGAAGGAGATCATTTAGGTGTTATTCCTCGCAACTATGAAGGAATAGTAAACCGTGTAACAGCAAGGTTCGGCCTAGATGCATCACAGCAAATCCGTCTGGAAGCAGAAGAAGAAAAATTAGCTCATTTGCCACTCGCTAAAACAGTATCCGTAGAAGAGCTTCTGCAATACGTGGAGCTTCAAGATCCTGTTACGCGCACGCAGCTTCGCGCAATGGCTGCTAAAACGGTCTGCCCGCCGCATAAAGTAGAGCTTGAAGCCTTGCTTGAAAAGCAAGCCTACAAAGAACAAGTGCTGGCAAAACGTTTAACAATGCTTGAACTGCTTGAAAAATACCCGGCGTGTGAAATGAAATTCAGCGAATTTATCGCCCTTCTGCCAAGCATACGCCCGCGCTATTACTCGATTTCTTCATCACCTCGTGTCGATGAAAAACAAGCAAGCATCACGGTCAGCGTTGTCTCAGGAGAAGCGTGGAGCGGATATGGAGAATATAAAGGAATTGCGTCGAACTATCTTGCCGAGCTGCAAGAAGGAGATACGATTACGTGCTTTATTTCCACACCGCAGTCAGAATTTACGCTGCCAAAAGACCCTGAAACGCCGCTTATCATGGTCGGACCGGGAACAGGCGTCGCGCCGTTTAGAGGCTTTGTGCAGGCGCGCAAACAGCTAAAAGAACAAGGACAGTCACTTGGAGAAGCACATTTATACTTCGGCTGCCGTTCACCTCATGAAGACTATCTGTATCAAGAAGAGCTTGAAAACGCCCAAAGCGAAGGCATCATTACGCTTCATACCGCTTTTTCTCGCATGCCAAATCAGCCGAAAACATACGTTCAGCACGTAATGGAACAAGACGGCAAGAAATTGATTGAACTTCTTGATCAAGGAGCGCACTTCTATATTTGCGGAGACGGAAGCCAAATGGCACCTGCCGTTGAAGCAACGCTTATGAAAAGCTATGCTGACGTTCACCAAGTGAGTGAAGCAGACGCTCGCTTATGGCTGCAGCAGCTAGAAGAAAAAGGCCGATACGCAAAAGACGTGTGGGCTGGGTAA

**Supplementary figures**

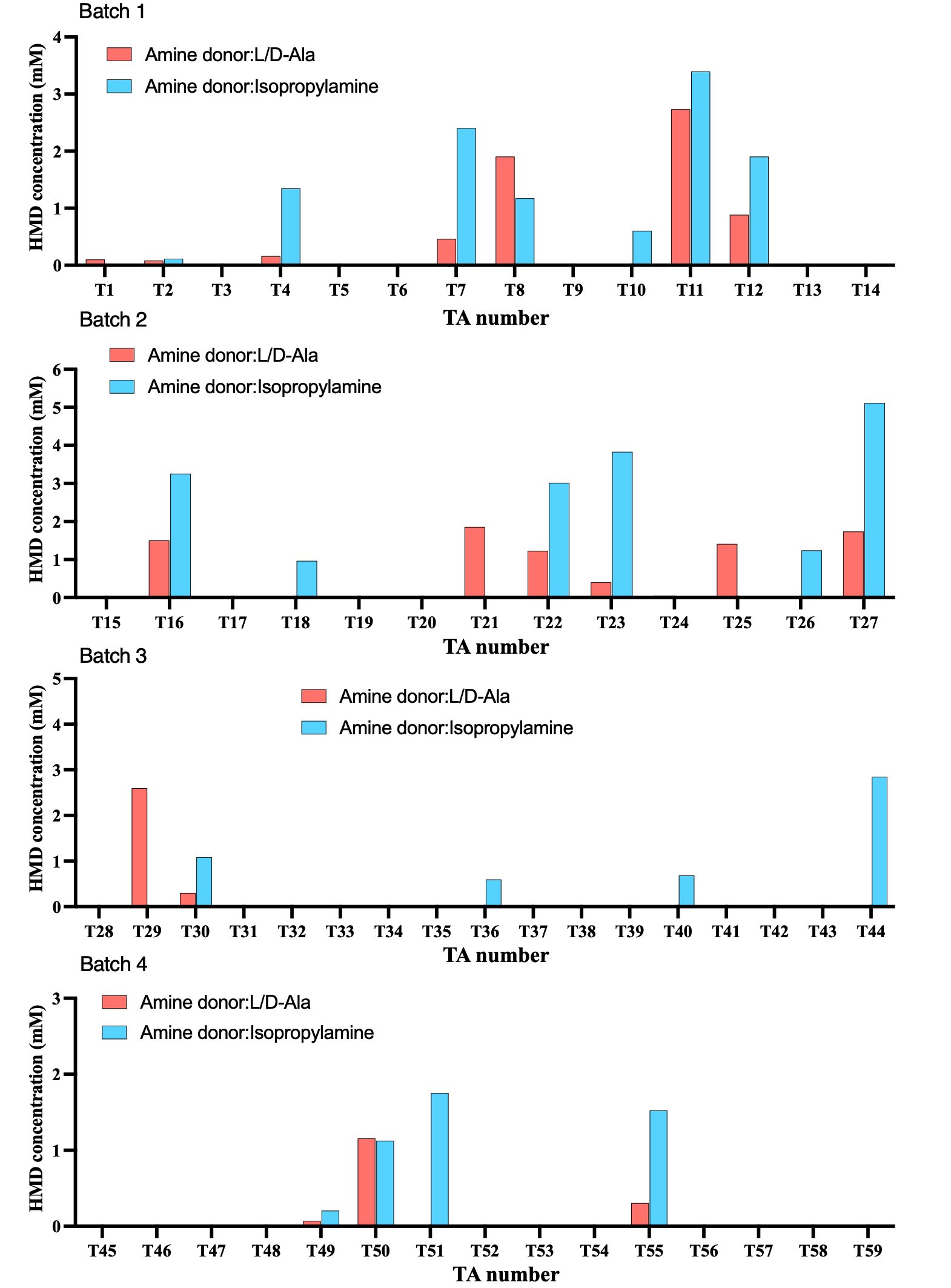

**Supplementary Figure 1.** **Screening of the ω-transaminases.** The screening process for the existing 59 transaminases in the library was divided into four batches with the same screening conditions. Two *E. coli* cells expressing transaminases (TAs) and alcohol dehydrogenase (ADH) were resuspended at a ratio of 1:1 in 3 mL phosphate buffer (pH 8.0, 100 mM) with a cell density of 8 g CDW L^-1^, 30 mM HDO. Reactions were performed at 25 °C, 220 rpm for 18 h, L/D-Ala (100 mM L-Ala and 100 mM D-Ala) or isopropylamine (100 mM) was employed as amine donor.

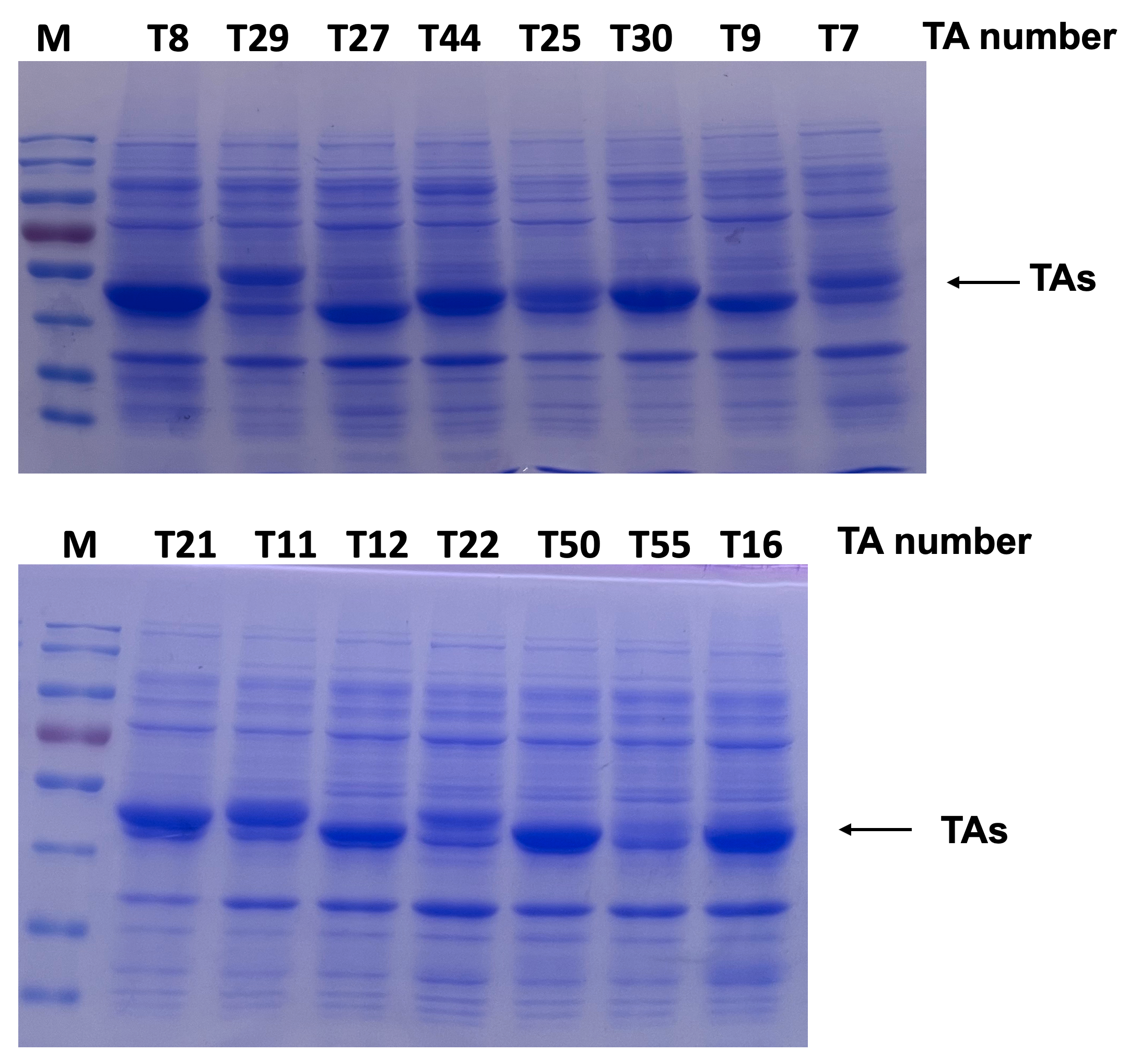

**Supplementary Figure 2. SDS-PAGE analysis of some high-performance transaminases (TAs) from the transaminases library expressed in *E. coli*.** Lane M: protein marker (kDa).

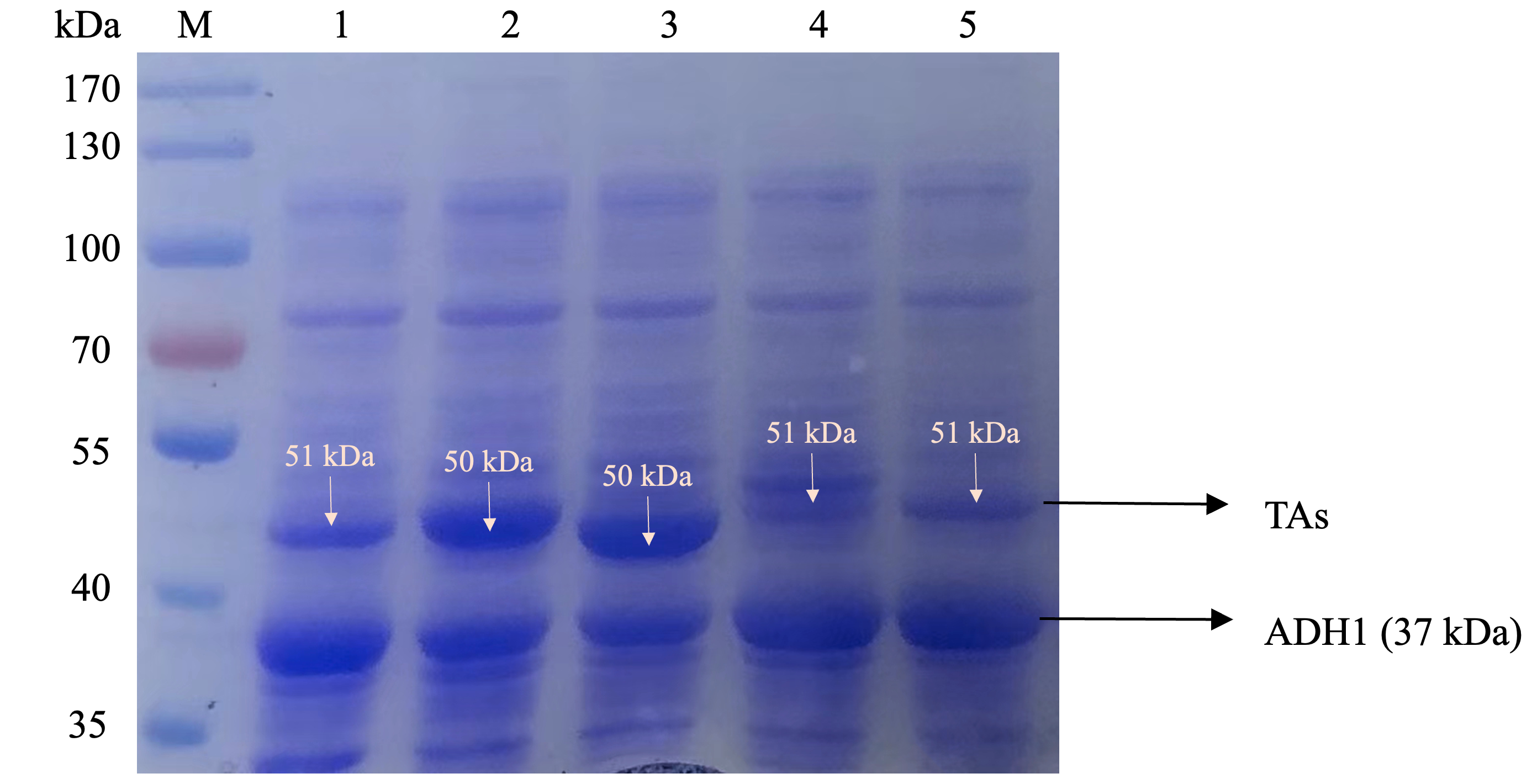

**Supplementary Figure 3. SDS-PAGE analysis of whole-cell proteins of cell module 3 expressed in *E. coli*.** Lane M: protein marker (kDa); Lane 1: *E. coli* (M3A); Lane 2: *E. coli* (M3D); Lane 3: *E. coli* (M3B); Lane 4: *E. coli* (M3E); Lane 5: *E. coli* (M3C).

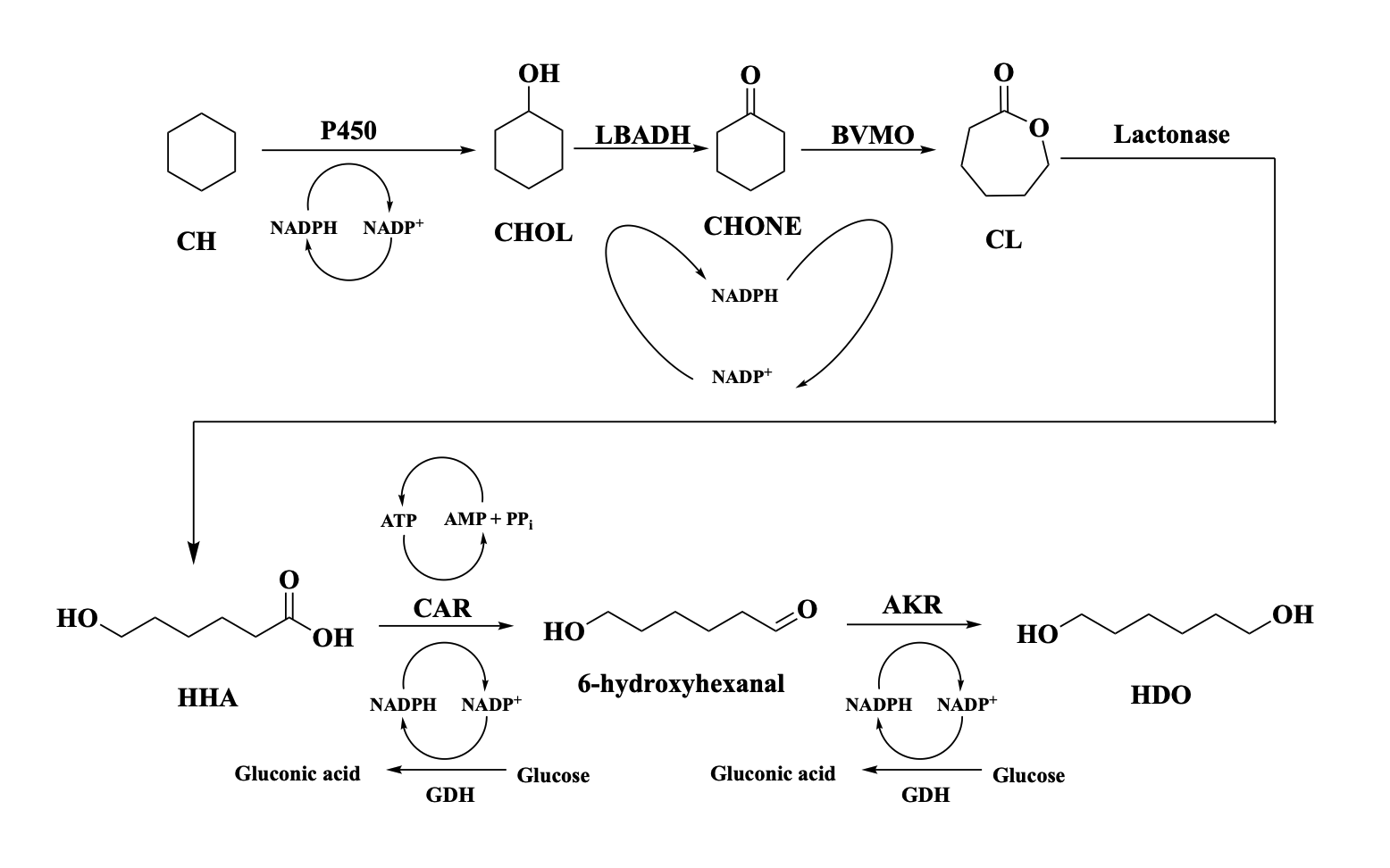

**Supplementary Figure 4．Previous work by our group using eight enzymes to synthesize HDO.**

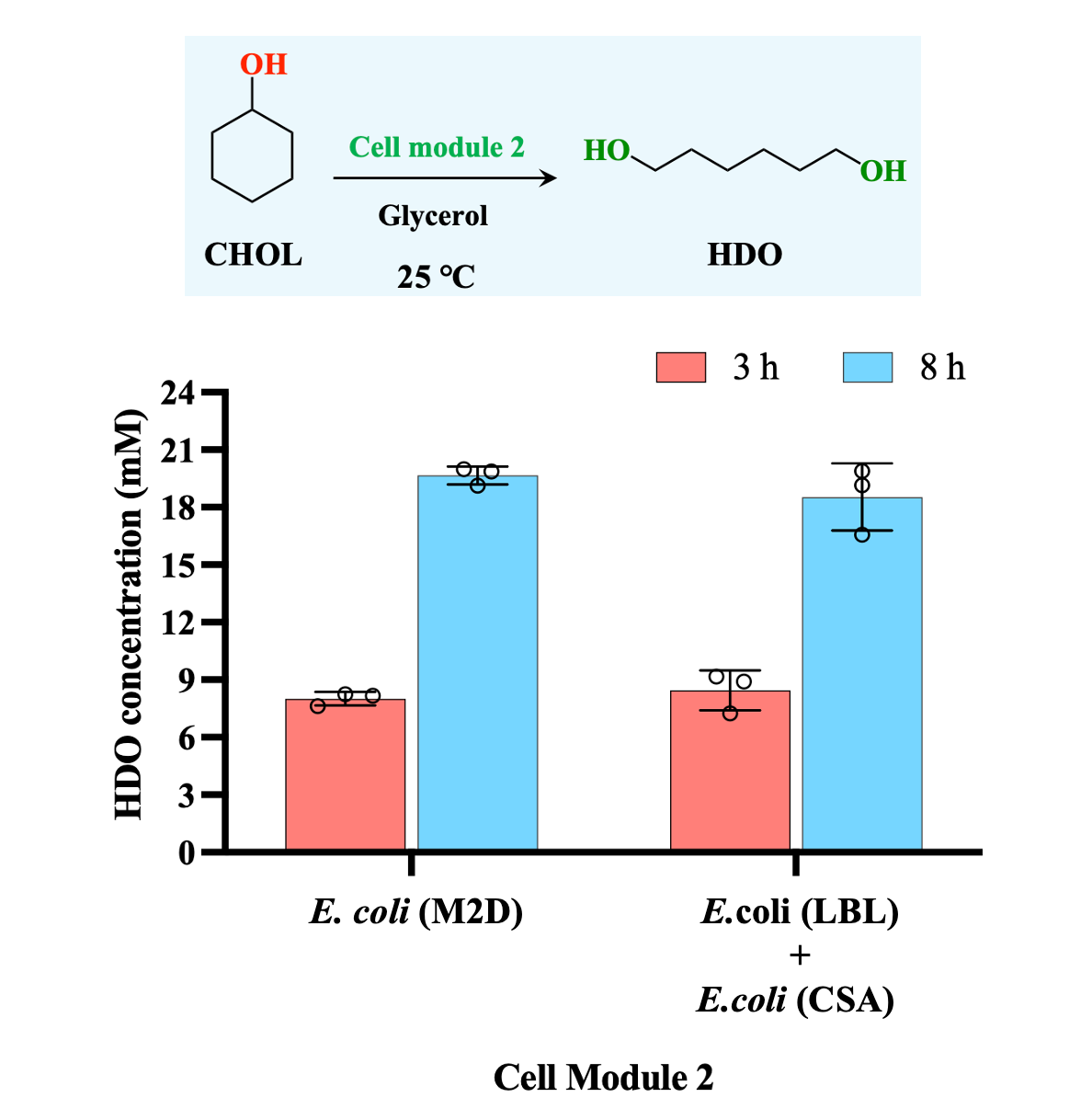

**Supplementary Figure 5. Engineering of *E. coli* cell module 2 for the conversion of CHOL to HDO.** Comparison of the catalytic performance of newly constructed *E. coli* (M2D) with two-cell system containing *E. coli* (LBL) and *E. coli* (CSA) of our previous work for the conversion of CHOL to HDO. Reaction conditions: *E. coli* (M2D) (final CDW is 6 g L^-1^) or two-cell system containing *E. coli* (LBL) and *E. coli* (CSA) (final CDW is 10.8 g L^-1^) at a ratio of 7:20 was resuspended in 3 mL phosphate buffer (pH 8.0, 100 mM), 20 mM CHOL. Reactions were performed at 25 °C, 220 rpm for 8 h, cofactor NAD(P)H/ATP was provided by the *E. coli* host cells using 68 mM glycerol as an energy source.

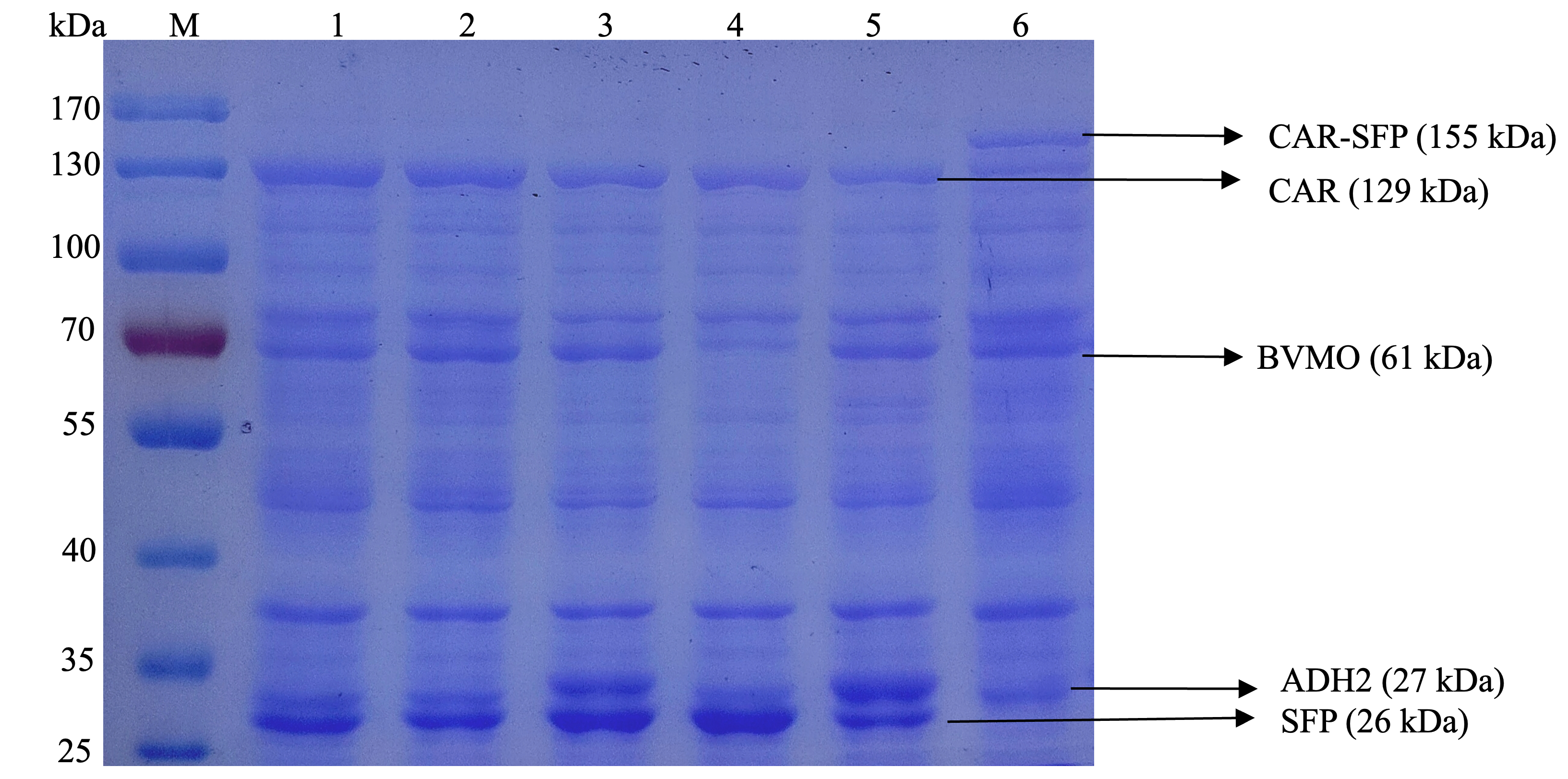

**Supplementary Figure 6. SDS-PAGE analysis of whole-cell proteins of cell module 2 expressed in *E. coli*.** Lane M: protein marker (kDa); Lane 1: *E. coli* (M2E); Lane 2: *E. coli* (M2D); Lane 3: *E. coli* (M2B); Lane 4: *E. coli* (M2C); Lane 5: *E. coli* (M2A); Lane 6: *E. coli* (M2F).

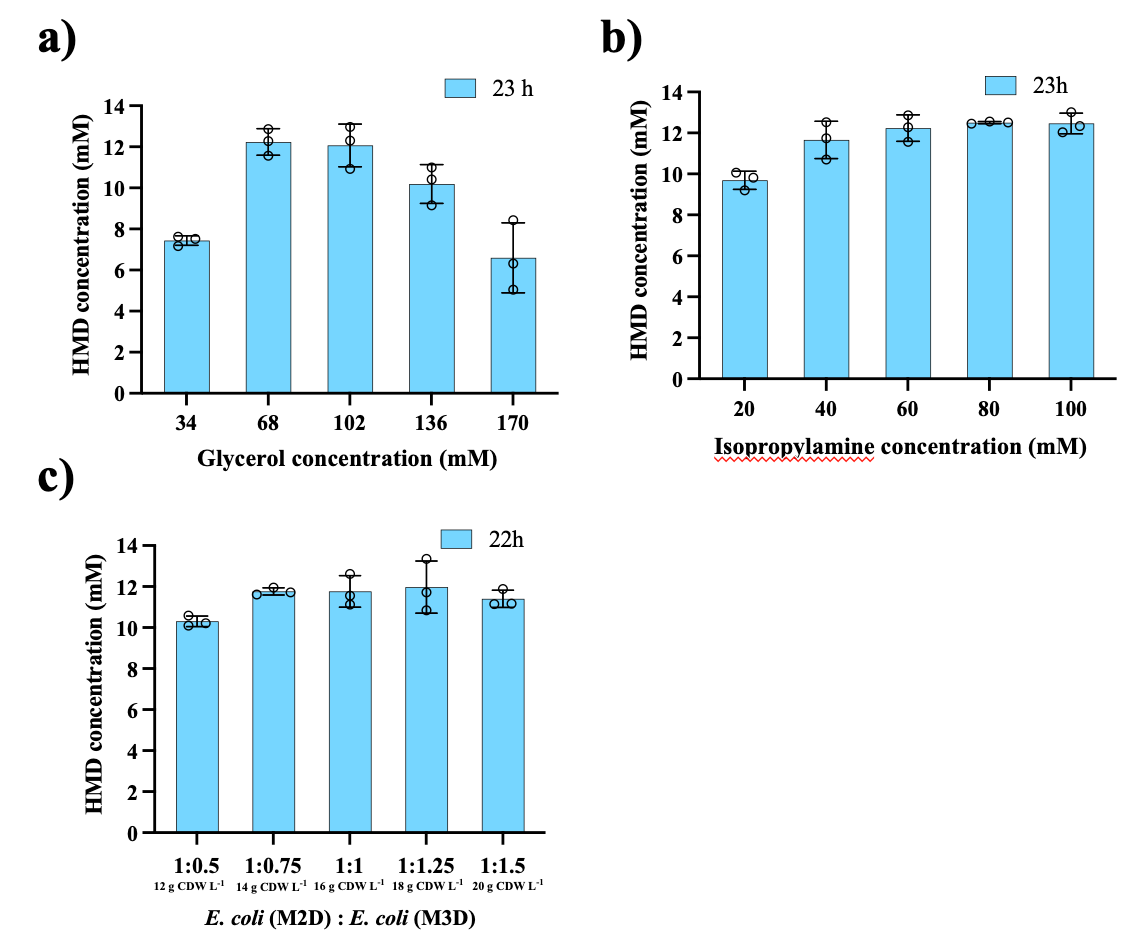

**Supplementary Figure 7.** **Optimization of experimental conditions for one-pot, two-step method with CHOL as substrate.** **a**) Optimization of glycerol concentration. Reaction conditions: *E. coli* consortium 2_3 composed of *E. coli* (M2D) and *E. coli* (M3D) at a ratio of 4:3 was resuspended in 3 mL phosphate buffer (pH 8.0, 100 mM) at a cell density of 14 g CDW L^−1^, 20 mM CHOL. Reactions were performed at 25 °C, 220 rpm for 23 h, cofactor NAD(P)H/ATP was provided by the *E. coli* host cells using glycerol as an energy source and 60 mM isopropylamine was added as amine donor. *E. coli* (M3D) and amine donor were added after 7.5 h reaction when the first step reaction was completed. b) Optimization of isopropylamine concentration. Reaction conditions: *E. coli* consortium 2_3 consisting of *E. coli* (M2D) and *E. coli* (M3D) at a ratio of 4:3 was resuspended in 3 mL phosphate buffer (pH 8.0, 100 mM) at a cell density of 14 g CDW L^-1^, 20 mM CHOL. Reactions were performed at 25 °C, 220 rpm for 23 h, cofactor NAD(P)H/ATP was provided by the *E. coli* host cells using 68 mM glycerol as an energy source and isopropylamine was employed as amine donor. *E. coli* (M3D) and amine donor were added after 7.5 h reaction. c) Optimization of cell loading ratio of *E. coli* (M2D) and *E. coli* (M3D). Reaction conditions: *E. coli* consortium 2_3 consisting of *E. coli* (M2D) and *E. coli* (M3D) at a designed ratio was resuspended in 3 mL phosphate buffer (pH 8.0, 100 mM) with a specified cell density, 20 mM CHOL. Reactions were performed at 25 °C, 220 rpm for 23 h, cofactor NAD(P)H/ATP was provided by the *E. coli* host cells using 68 mM glycerol as an energy source and 60 mM isopropylamine was employed as amine donor. *E. coli* (M3D) and amine donor were added after 7.5 h reaction.

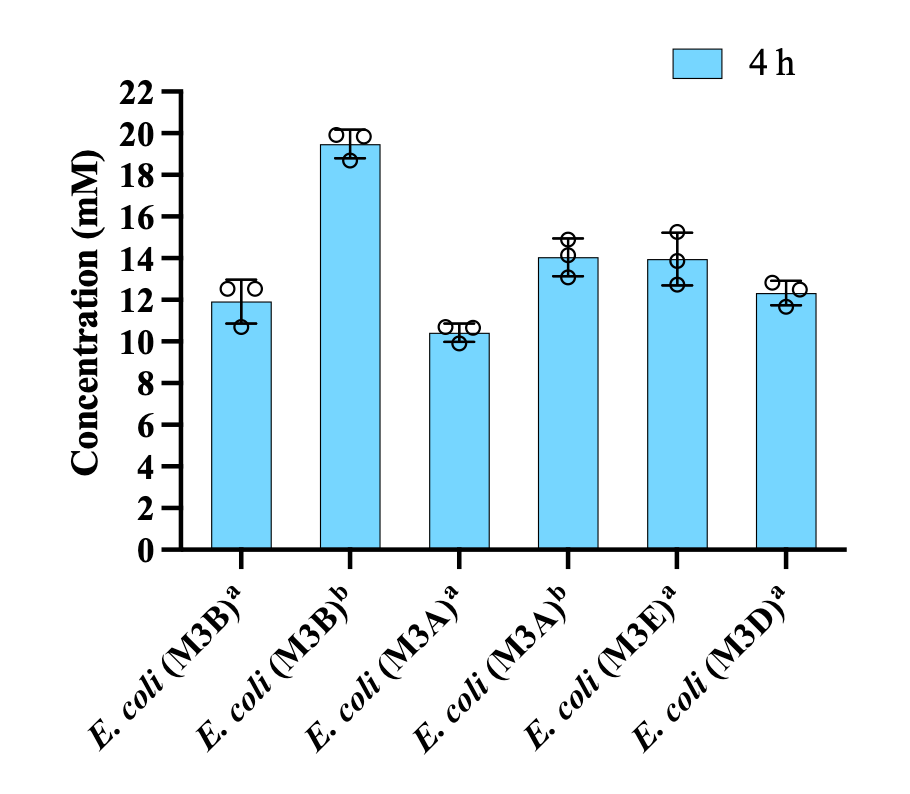

**Supplementary Figure 8. Biocatalytic synthesis of cyclohexylamine from CHONE.** Reaction conditions: *E. coli* (M3A, M3B, M3D or M3E) was resuspended in 3 mL phosphate buffer (pH 8.0, 100 mM) at a cell density of 8 g CDW L^-1^, 30 mM CHONE. Reactions were performed at 25 °C, 220 rpm for 4 h, 60 mM **^a^**L-Ala or **^b^**isopropylamine was added as amine donor. Samples were taken at appropriate intervals and prepared for GC analysis. In order to determine cyclohexylamine, each reaction sample was prepared for GC analysis with the SH-Rtx-1 column as follows: 333 µL of potassium phosphate buffer (0.1 M, pH 8.0), 500 µL of ethyl acetate containing 4 mM n-decane (internal standard) and 5 μL 10 M NaOH were added to 167 µL of the reaction sample, followed by vortexing and centrifugation (13,680 × *g*, 1 min). The organic phase was dried over anhydrous Na_2_SO_4_ and then directly used for GC analysis. The procedure for GC analysis with the SH-Rtx-1 column: the temperature program was as follows: 5 °C min^−1^ from 65 °C to 110 °C, 55 °C min^−1^ to 240 °C and held at 240 °C for 1 min. All the experiments were performed in triplicate, and the error bars indicate the standard deviations.

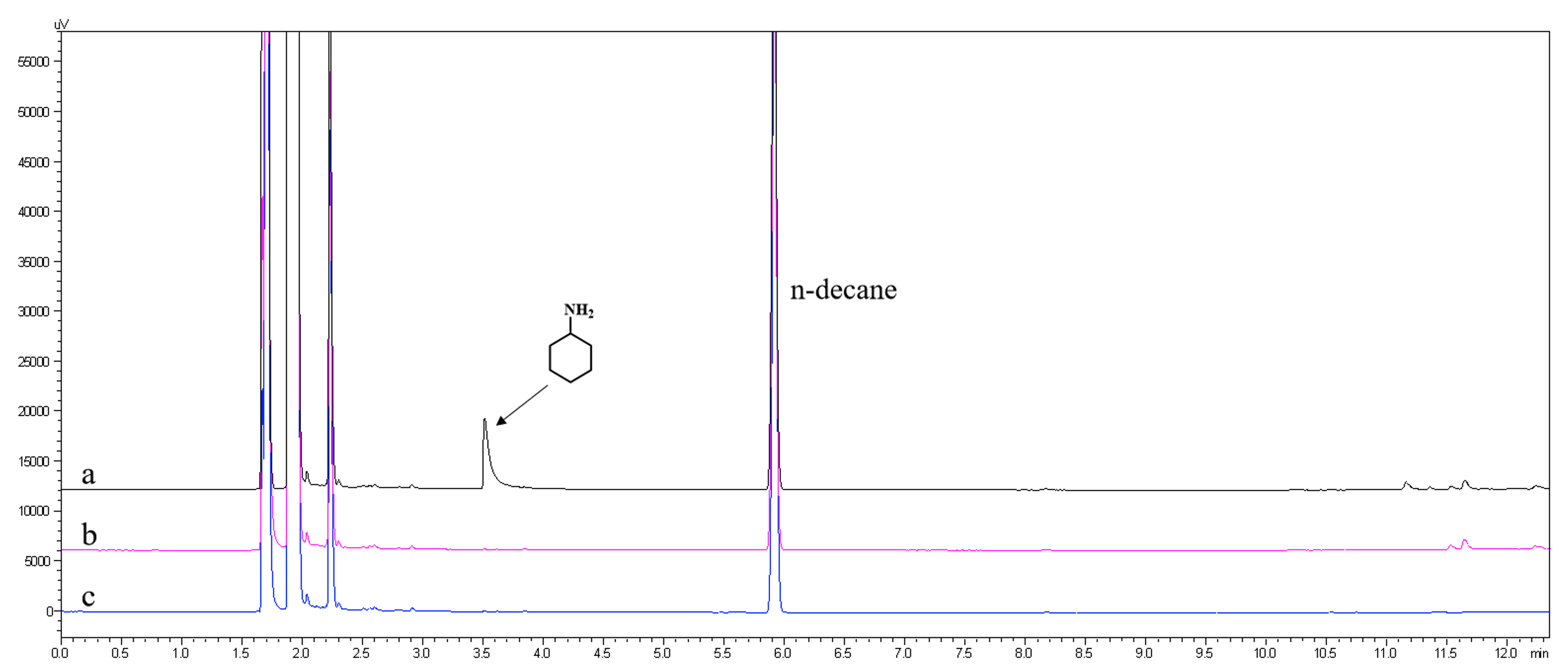

**Supplementary Figure 9. GC analysis of cyclohexylamine during the conversion of CH or CHOL to HMD.** a) standard of cyclohexylamine. b) *E. coli* consortium 1_2_3 catalyzed conversion of CH to HMD. c) *E. coli* consortium 2_3 catalyzed conversion of CHOL to HMD.

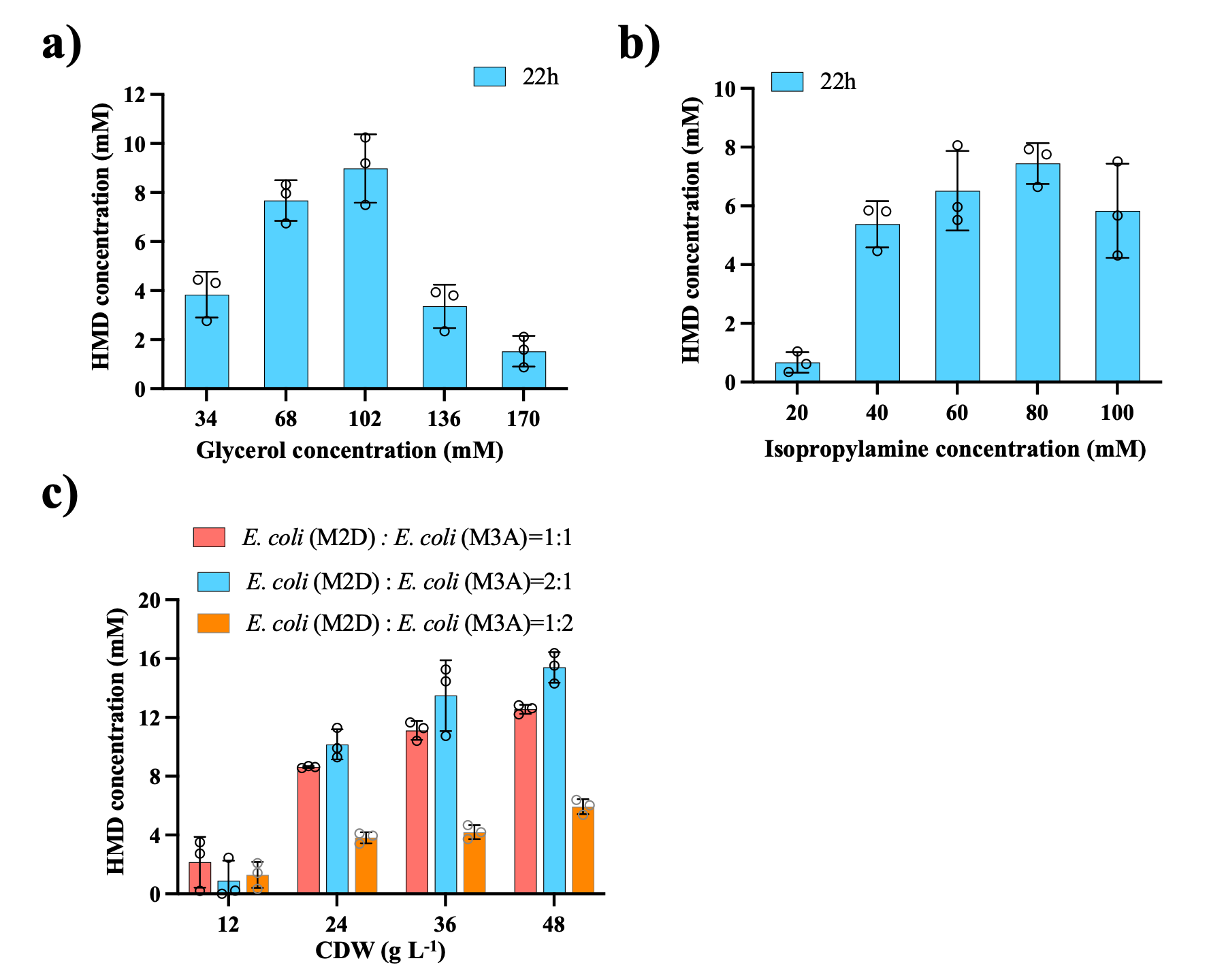

**Supplementary Figure 10.** **Optimization of experimental conditions for one-pot, one-step method with CHOL as substrate**. a) Optimization of glycerol concentration. Reaction conditions: *E. coli* consortium 2_3 composed of *E. coli* (M2D) and *E. coli* (M3A) at a ratio of 4:3 was resuspended in 3 mL phosphate buffer (pH 8.0, 100 mM) at a cell density of 14 g CDW L^−1^, 20 mM CHOL. Reactions were performed at 25 °C, 220 rpm for 22 h, cofactor NAD(P)H/ATP was provided by the *E. coli* host cells using glycerol as an energy source and 60 mM isopropylamine was added as amine donor. b) Optimization of isopropylamine concentration. Reaction conditions: *E. coli* consortium consisting of *E. coli* (M2D) and *E. coli* (M3A) at a ratio of 4:3 was resuspended in 3 mL phosphate buffer (pH 8.0, 100 mM) at a cell density of 14 g CDW L^-1^, 20 mM CHOL. Reactions were performed at 25 °C, 220 rpm for 22 h, cofactor NAD(P)H/ATP was provided by the *E. coli* host cells using 68 mM glycerol as an energy source and isopropylamine was employed as amine donor. c) Optimization of cell loading ratio of *E. coli* (M2D) and *E. coli* (M3A). Reaction conditions: *E. coli* consortium consisting of *E. coli* (M2D) and *E. coli* (M3A) at a designed ratio was resuspended in 3 mL phosphate buffer (pH 8.0, 100 mM) with a specified cell density, 20 mM CHOL. Reactions were performed at 25 °C, 220 rpm for 22 h, cofactor NAD(P)H/ATP was provided by the *E. coli* host cells using 68 mM glycerol as an energy source and 60 mM isopropylamine was employed as amine donor.

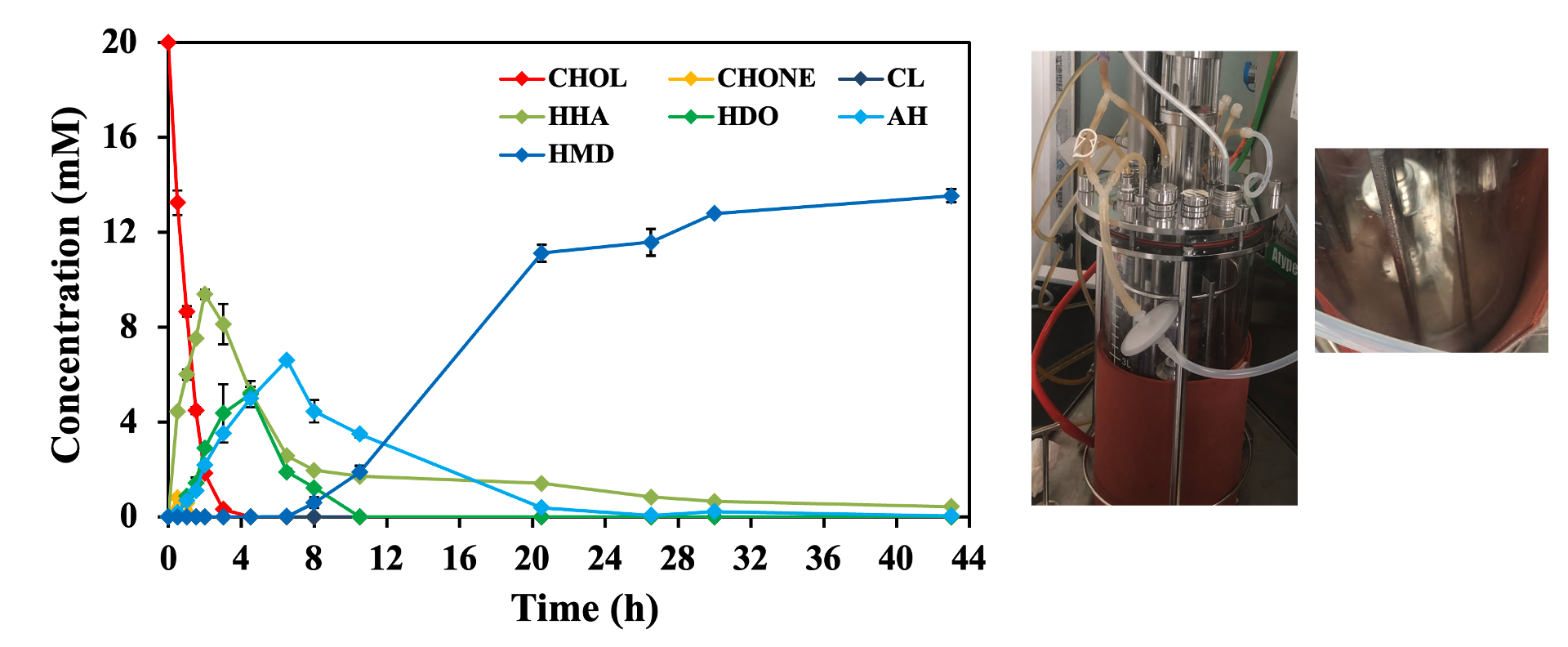

**Supplementary Figure 11. Biosynthesis of HMD from CHOL in 600 mL scale inone-pot, one-step process.** Reaction conditions: *E. coli* consortium 2_3 composed of *E. coli* (M2D) and *E. coli* (M3A) at a ratio of 2:1 was resuspended in 600 mL phosphate buffer (pH 8.0, 100 mM) at a cell density of 20 g CDW L^−1^, 20 mM CHOL. Reactions were performed at 25 °C, 400 rpm for 44 h, cofactor NAD(P)H/ATP was provided by the *E. coli* host cells using 102 mM glycerol as an energy source and 80 mM isopropylamine was added as amine donor.

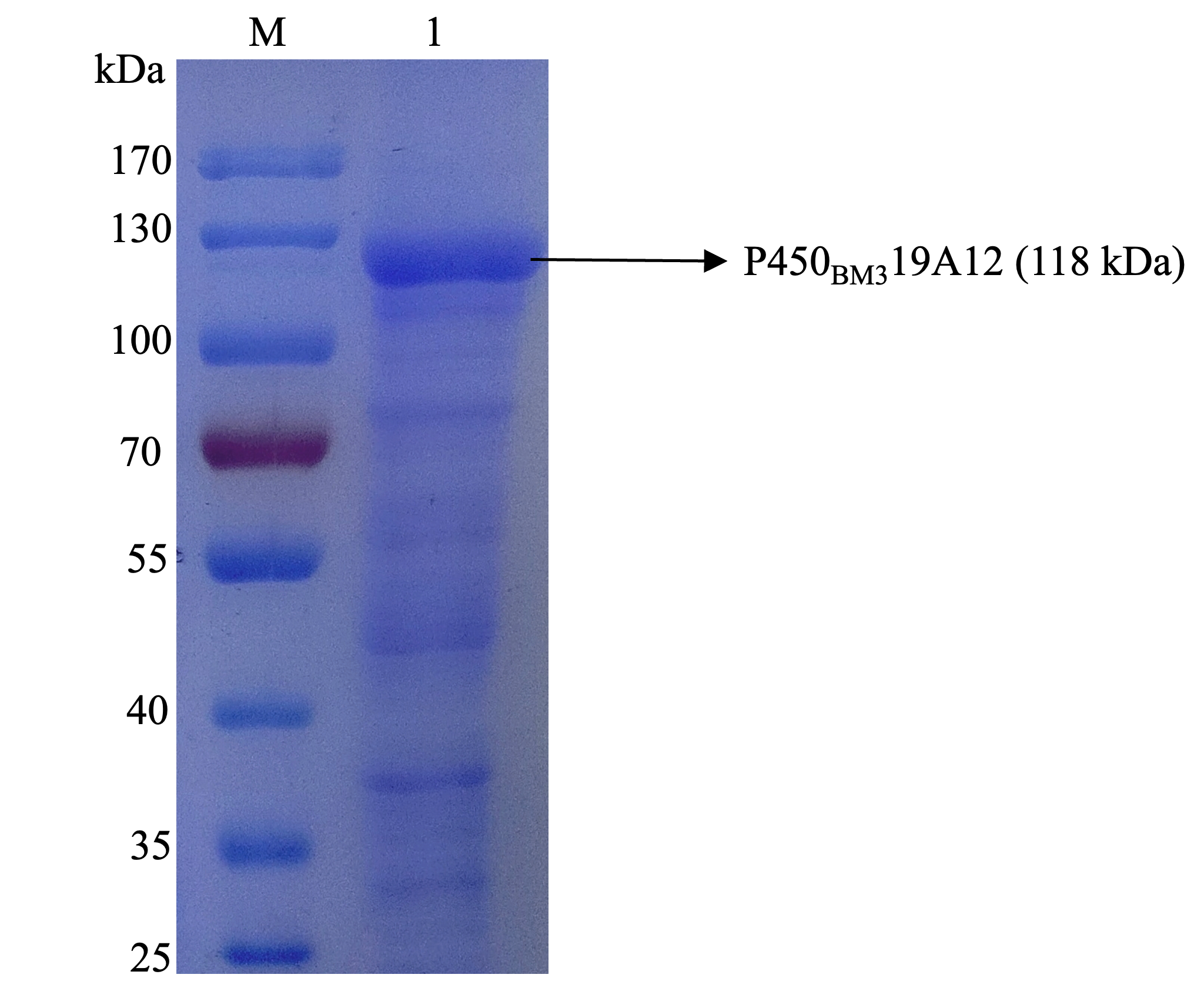

**Supplementary Figure 12. SDS-PAGE analysis of whole-cell proteins of cell module 1 expressed in *E. coli*.** Lane M: protein marker (kDa); Lane 1: *E. coli* (M1).

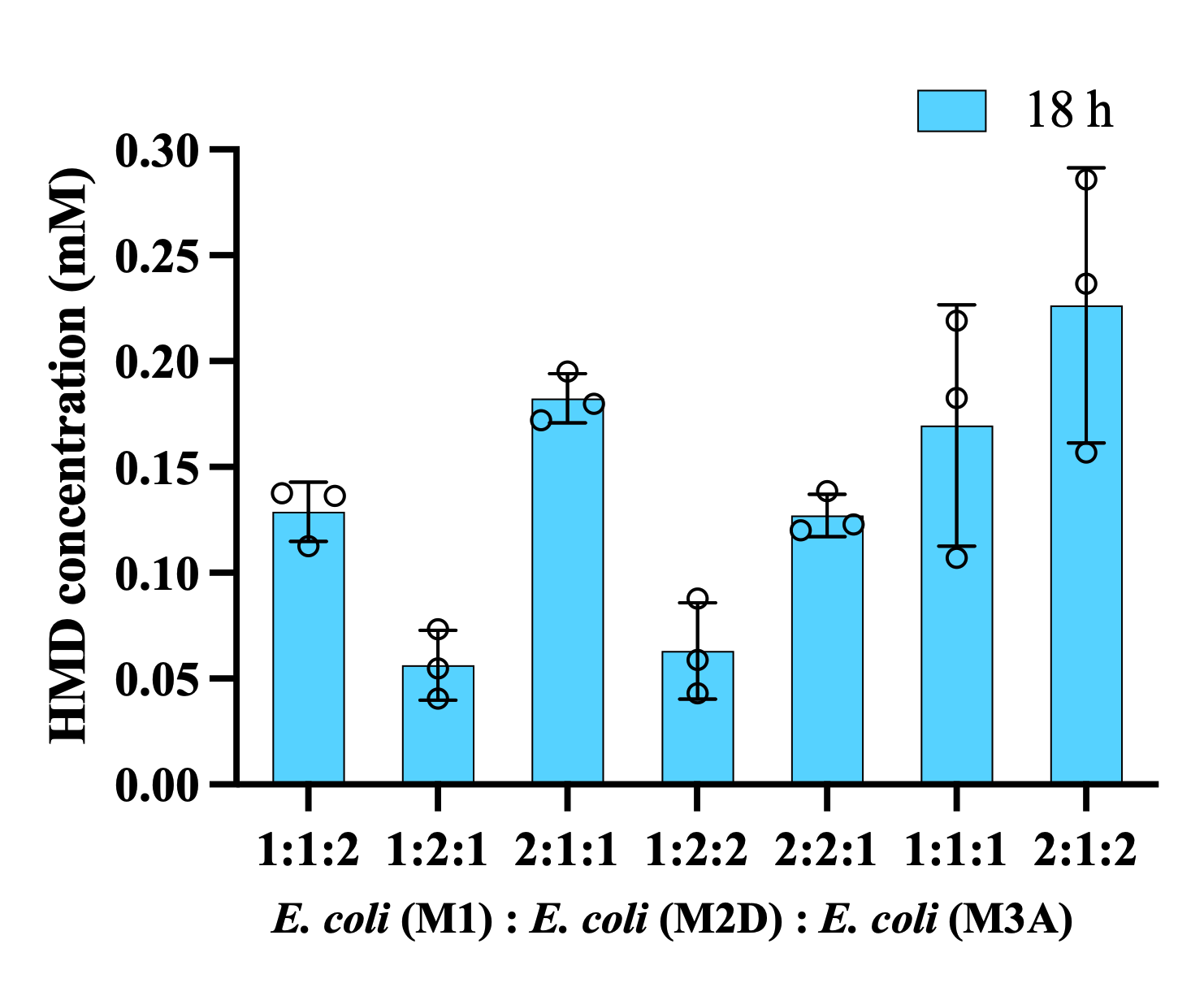

**Supplementary Figure 13.** **Optimization of cell module ratio of *E. coli* (M1), *E. coli* (M2D) and *E. coli* (M3A) from CH to HMD in a one-pot, one-step process.** Reaction conditions: *E. coli* consortium 1_2_3 composed of *E. coli* (M1), *E. coli* (M2D) and *E. coli* (M3A) at a designed ratio was resuspended in 3 mL phosphate buffer (pH 8.0, 100 mM) at a specified cell density, 30 mM CH. Reactions were performed at 25 °C, 220 rpm for 18 h, cofactor NAD(P)H/ATP was provided by the *E. coli* host cells using 136 mM glycerol as an energy source and 80 mM isopropylamine was added as amine donor.

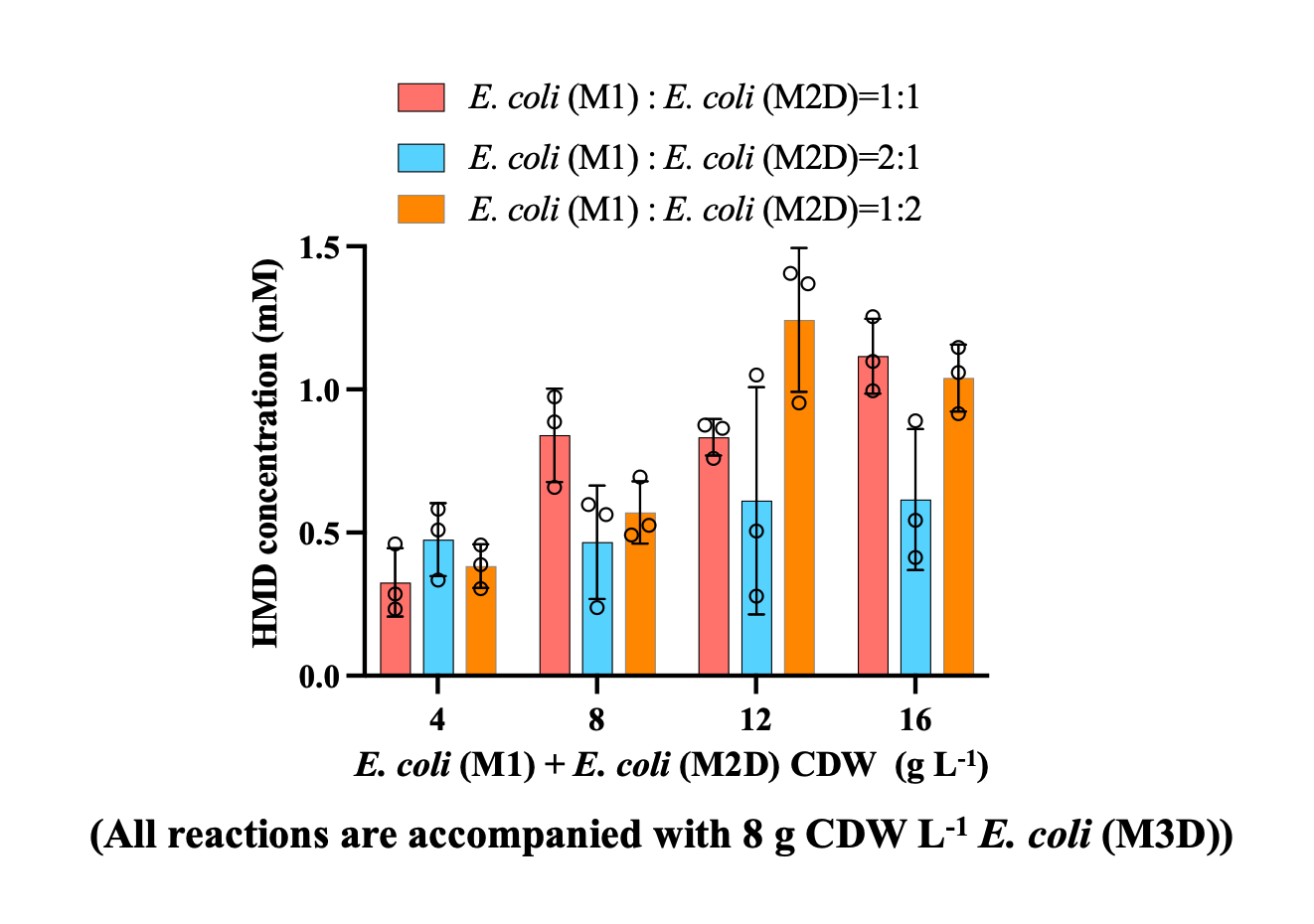

**Supplementary Figure 14.** **Optimization of the cell module ratio of *E. coli* (M1) and *E. coli* (M2D) from CH to HMD in a one-pot, two-step process.** Reaction conditions: *E. coli* consortium 1_2_3 composed of *E. coli* (M1), *E. coli* (M2D) and *E. coli* (M3D) at a designed ratio was resuspended in 3 mL phosphate buffer (pH 8.0, 100 mM) at a specified cell density, 30 mM CH. Reactions were performed at 25 °C, 220 rpm for 20 h, cofactor NAD(P)H/ATP was provided by the *E. coli* host cells using 136 mM glycerol as an energy source and 80 mM isopropylamine was added as amine donor. *E. coli* (M3D) and amine donor were added after 7 h reaction when the first step reaction was completed.

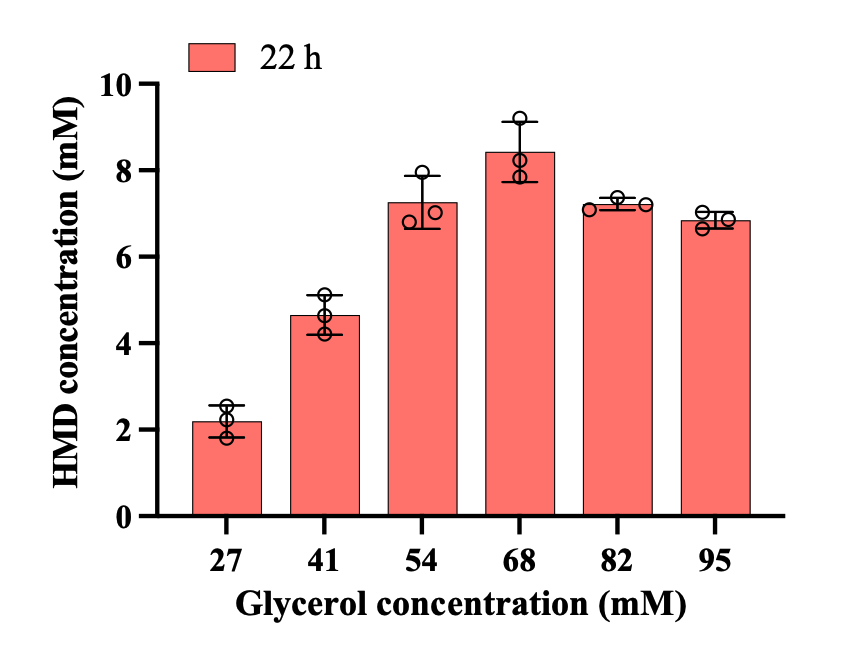

**Supplementary Figure 15. Optimization of glycerol concentration from CH to HMD in a one-pot, two-step process.** Reaction conditions: *E. coli* consortium 1_2_3 composed of *E. coli* (M1), *E. coli* (M2D) and *E. coli* (M3D) at a ratio of 3:3:4 was resuspended in 3 mL phosphate buffer (pH 8.0, 100 mM) at a cell density of 20 g CDW L^-1^, 30 mM CH. Reactions were performed at 25 °C, 220 rpm for 22 h, cofactor NAD(P)H/ATP was provided by the *E. coli* host cells using glycerol as an energy source and 80 mM isopropylamine was added as amine donor. *E. coli* (M3D) and amine donor were added after 8 h reaction when the first step reaction was completed.

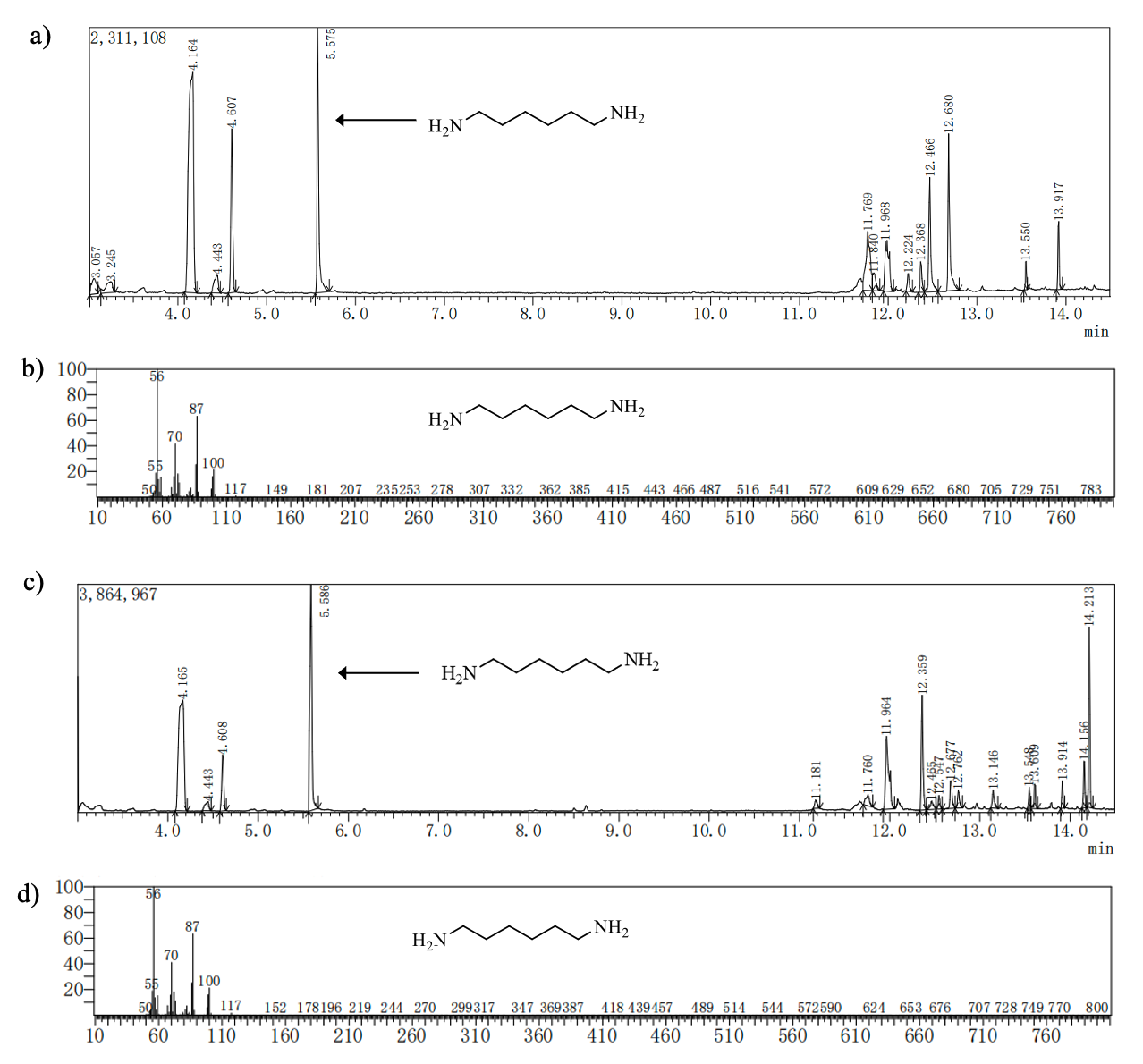

**Supplementary Figure 16. GC-MS analysis of reaction mixtures from *E. coli* consortium 2_3 catalyzed conversion of CHOL to HMD.** a) GC chromatograms of HMD standard. b) Mass spectrometry analysis of HMD standard. The fragmentation pattern was obtained for HMD standard. c) GC chromatograms of *E. coli* consortium 2_3 catalyzed conversion of cyclohexanol to HMD. d) Mass spectrometry analysis of HMD product. The fragmentation pattern was obtained for HMD product. The GC–MS analysis was performed using the SHIMADZU GCMS-QP2010 SE equipped with a Rtx-5MS column (30 m×0.25 mm, 0.25 µm).

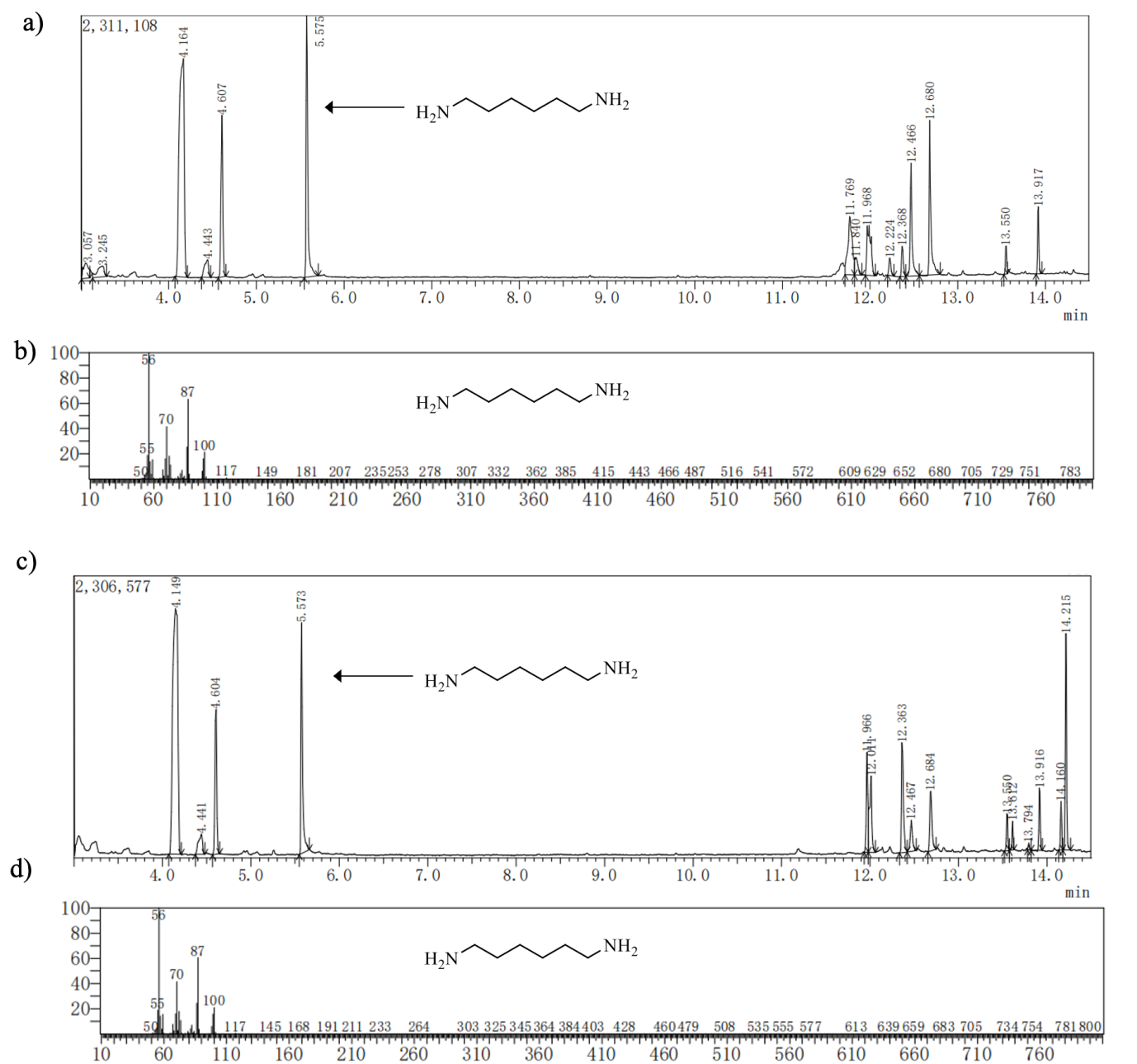

**Supplementary Figure 17. GC-MS analysis of reaction mixtures from *E. coli* consortium 1_2_3 catalyzed conversion of cyclohexane to HMD.** a) GC chromatograms of HMD standard. b) Mass spectrometry analysis of HMD standard. The fragmentation pattern was obtained for HMD standard. c) GC chromatograms of *E. coli* consortium 1_2_3 catalyzed conversion of cyclohexane to HMD. d) Mass spectrometry analysis of HMD product. The fragmentation pattern was obtained for HMD product. The GC–MS analysis was performed using the SHIMADZU GCMS-QP2010 SE equipped with a Rtx-5MS column (30 m×0.25 mm, 0.25 µm).

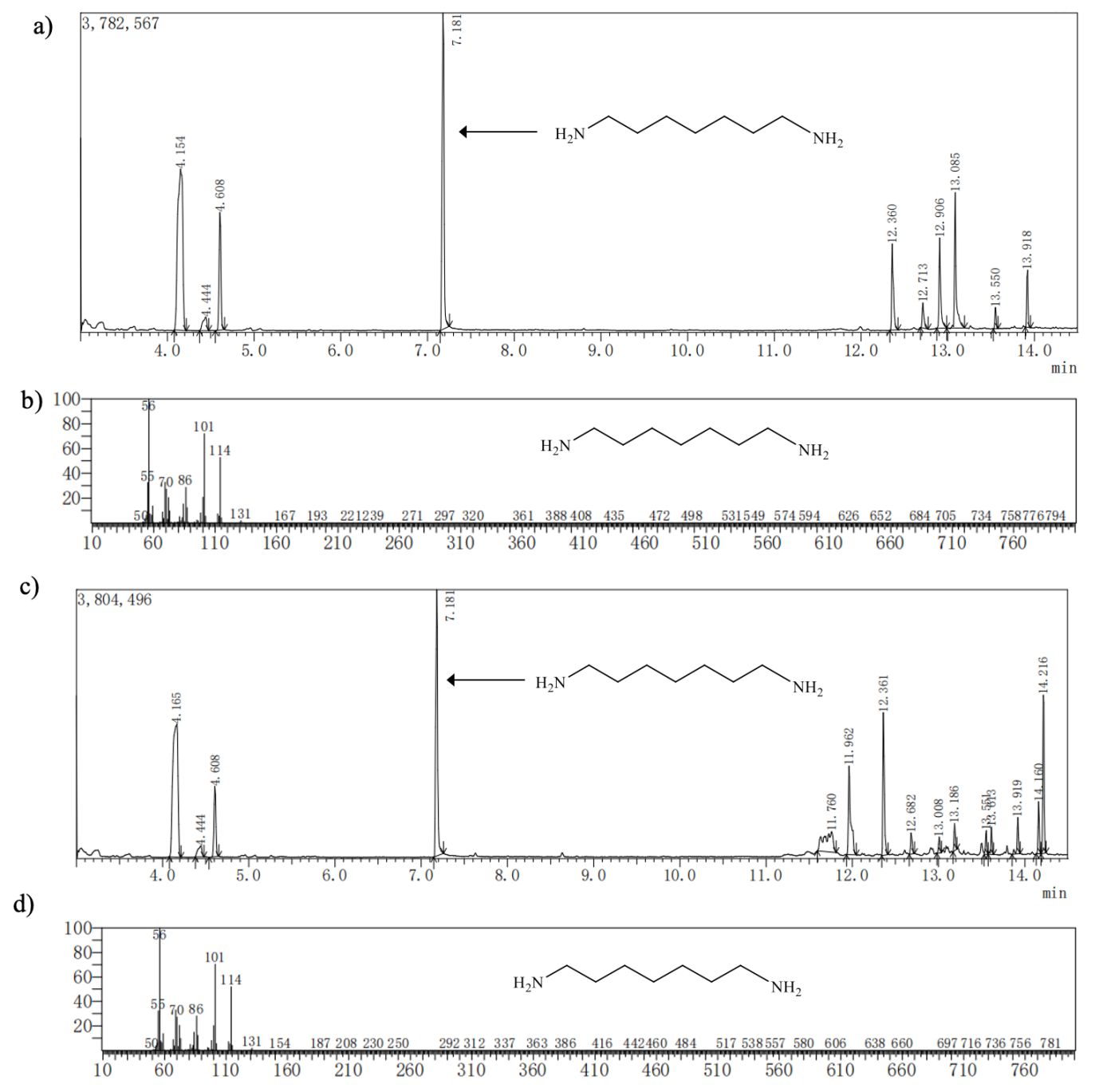

**Supplementary Figure 18. GC-MS analysis of reaction mixtures from *E. coli* consortium 2_3 catalyzed conversion of cycloheptanol to 1,7-heptanediamine.** a) GC chromatograms of 1,7-heptanediamine standard. b) Mass spectrometry analysis of 1,7-heptanediamine standard. The fragmentation pattern was obtained for 1,7-heptanediamine standard. c) GC chromatograms of *E. coli* consortium 2_3 catalyzed conversion of cycloheptanol to 1,7-heptanediamine. d) Mass spectrometry analysis of 1,7-heptanediamine product. The fragmentation pattern was obtained for 1,7-heptanediamine product. The GC–MS analysis was performed using the SHIMADZU GCMS-QP2010 SE equipped with a Rtx-5MS column (30 m×0.25 mm, 0.25 µm).

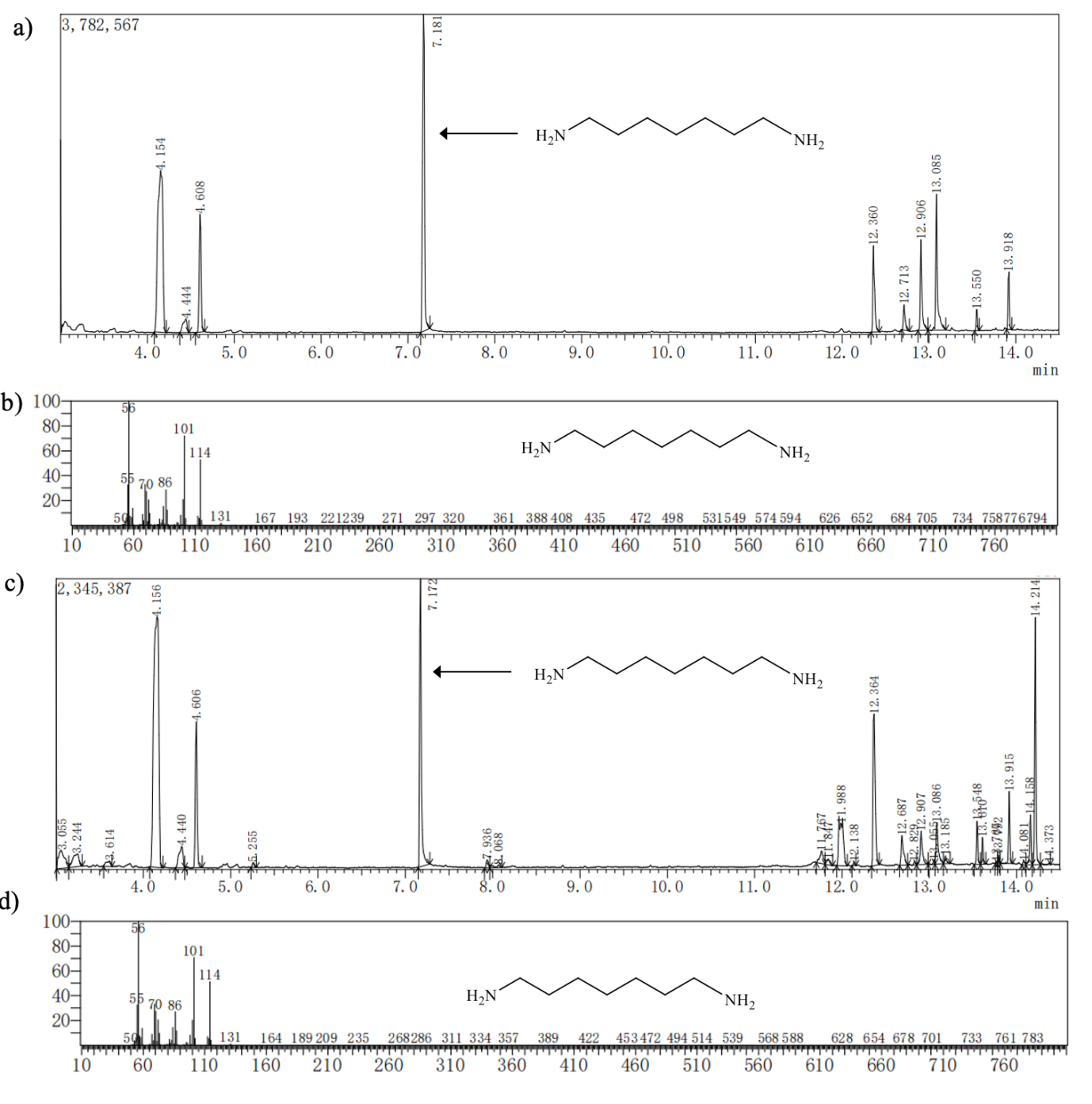

**Supplementary Figure 19. GC-MS analysis of reaction mixtures from *E. coli* consortium 1_2_3 catalyzed conversion of cycloheptane to 1,7-heptanediamine.** a) GC chromatograms of 1,7-heptanediamine standard. b) Mass spectrometry analysis of 1,7-heptanediamine standard. The fragmentation pattern was obtained for 1,7-heptanediamine standard. c) GC chromatograms of *E. coli* consortium 1_2_3 catalyzed conversion of cycloheptane to 1,7-heptanediamine . d) Mass spectrometry analysis of 1,7-heptanediamine product. The fragmentation pattern was obtained for 1,7-heptanediamine product. The GC–MS analysis was performed using the SHIMADZU GCMS-QP2010 SE equipped with a Rtx-5MS column (30 m×0.25 mm, 0.25 µm).

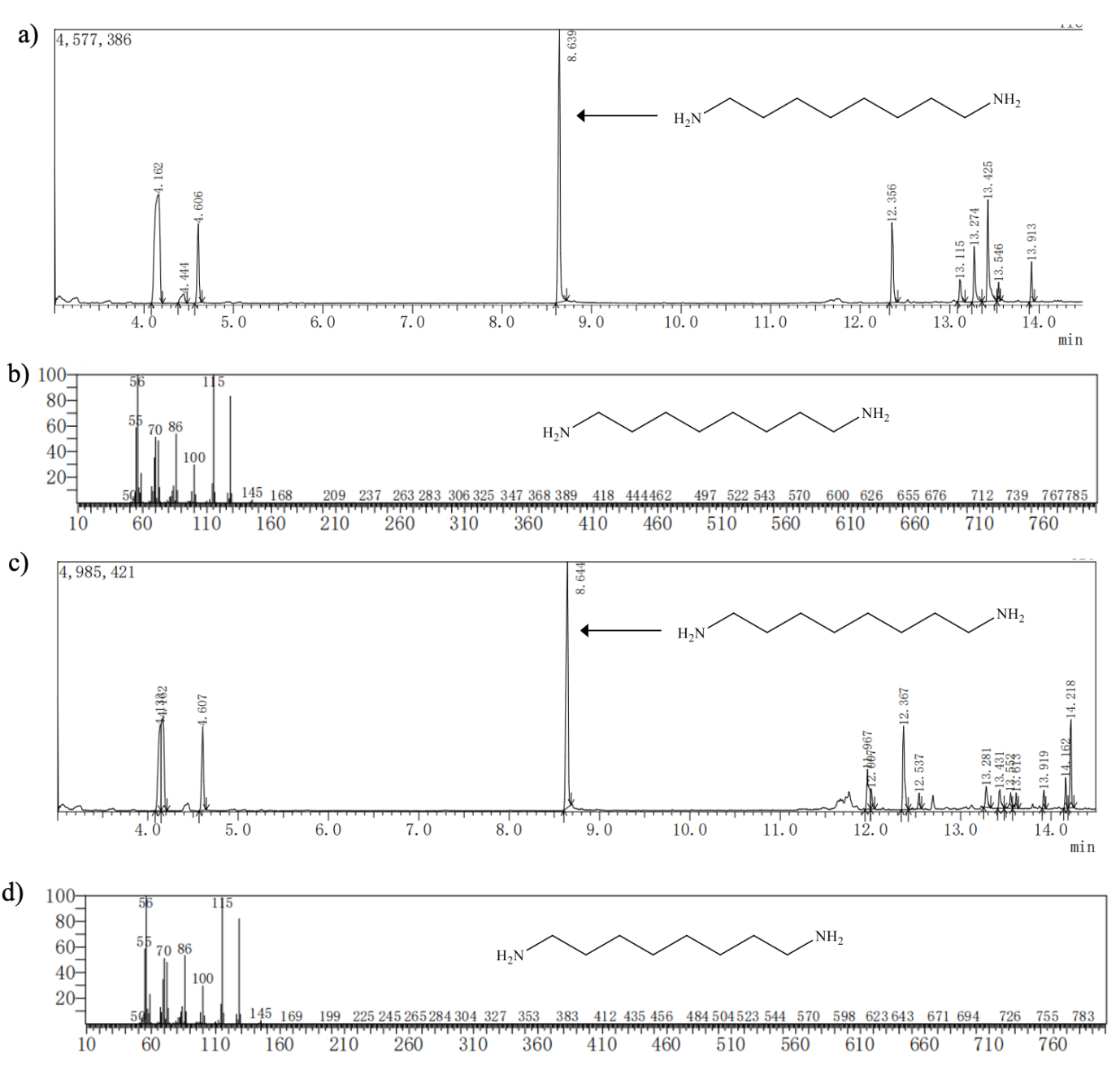

**Supplementary Figure 20. GC-MS analysis of reaction mixtures from *E. coli* consortium 2_3 catalyzed conversion of cyclooctanol to 1,8-octanediamine.** a) GC chromatograms of 1,8-octanediamine standard. b) Mass spectrometry analysis of 1,8-octanediamine standard. The fragmentation pattern was obtained for 1,8-octanediamine standard. c) GC chromatograms of *E. coli* consortium 2_3 catalyzed conversion of cyclooctanol to 1,8-octanediamine. d) Mass spectrometry analysis of 1,8-octanediamine product. The fragmentation pattern was obtained for 1,8-octanediamine product. The GC–MS analysis was performed using the SHIMADZU GCMS-QP2010 SE equipped with a Rtx-5MS column (30 m×0.25 mm, 0.25 µm).

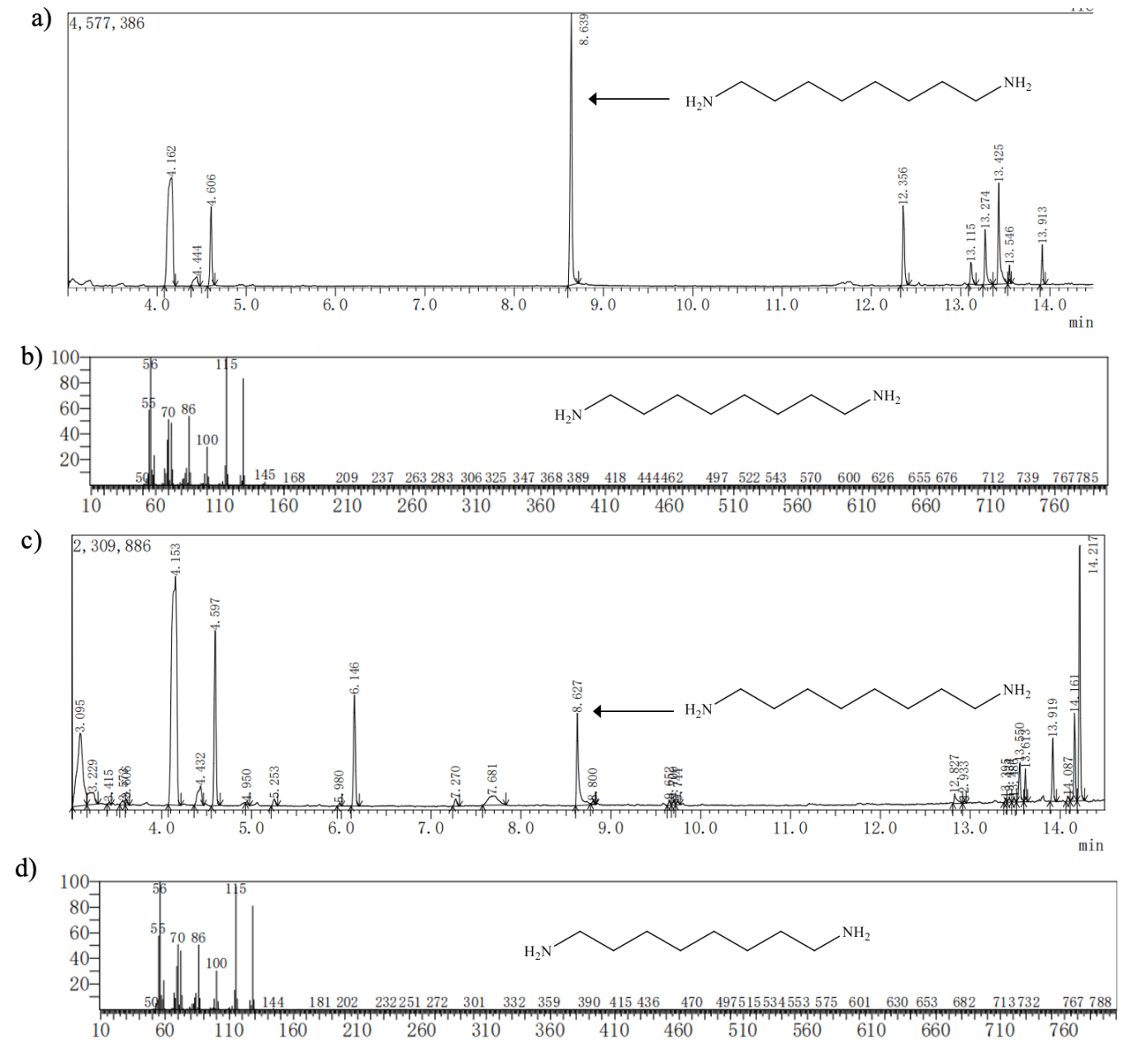

**Supplementary Figure 21. GC-MS analysis of reaction mixtures from *E. coli* consortium 1_2_3 catalyzed conversion of cyclooctane to 1,8-octanediamine.** a) GC chromatograms of 1,8-octanediamine standard. b) Mass spectrometry analysis of 1,8-octanediamine standard. The fragmentation pattern was obtained for 1,8-octanediamine standard. c) GC chromatograms of *E. coli* consortium 1_2_3 catalyzed conversion of cyclooctane to 1,8-octanediamine. d) Mass spectrometry analysis of 1,8-octanediamine product. The fragmentation pattern was obtained for 1,8-octanediamine product. The GC–MS analysis was performed using the SHIMADZU GCMS-QP2010 SE equipped with a Rtx-5MS column (30 m×0.25 mm, 0.25 µm).

**References:**

1.	Rodriguez, C. et al. Steric vs. electronic effects in the *Lactobacillus brevis* ADH-catalyzed bioreduction of ketones. *Org Biomol Chem* **12**, 673-681 (2014).

2.	van der Vlugt-Bergmans, C. J. B. & van der Werf, M. J. Genetic and biochemical characterization of a novel monoterpene ɛ-lactone hydrolase from *Rhodococcus erythropolis* DCL14. *Appl. Environ. Microbiol.* **67**, 733-741 (2001).

3.	Opperman, D. J. & Reetz, M. T. Towards practical Baeyer-Villiger-monooxygenases: design of cyclohexanone monooxygenase mutants with enhanced oxidative stability. *Chembiochem* **11**, 2589-2596 (2010).

4.	Khusnutdinova, A. N. et al. Exploring bacterial carboxylate reductases for the reduction of bifunctional carboxylic acids. *Biotechnol. J.* **12**, 1600751 (2017).

5.	Cheng, Q., Thomas, S. M., Kostichka, K., Valentine, J. R. & Nagarajan, V. Genetic analysis of a gene cluster for cyclohexanol oxidation in *Acinetobacter* sp. strain SE19 by in vitro transposition. *J. Bacteriol.* **182**, 4744-4751 (2000).

6.	Yu, H. L. et al. Bioamination of alkane with ammonium by an artificially designed multienzyme cascade. *Metab. Eng.* **47**, 184-189 (2018).

7.	Zhang, Z. et al. One-pot biosynthesis of 1,6-hexanediol from cyclohexane by *de novo* designed cascade biocatalysis. *Green Chem.* **22**, 7476-7483 (2020).
